# Evolutionary Constraints on Positional Sequence, Collective Properties and Sequence Style of Tropoelastin Dictated by Fundamental Requirements for Formation and Function of the Extracellular Elastic Matrix

**DOI:** 10.1101/2024.12.03.626628

**Authors:** Fred W. Keeley

## Abstract

Elastin is an unusual extracellular matrix protein responsible for the properties of extension and energy-efficient elastic recoil of large blood vessels, heart valves, lung parenchyma and many other vertebrate tissues requiring such resilience. Polymeric elastin is assembled from monomeric tropoelastin by a process involving liquid-liquid phase separation, followed by maturation into an extended elastic matrix, covalently cross-linked through the side chains of lysine residues in the protein and producing a robust biomaterial with the structural integrity to withstand hundreds of millions of cycles of extension and recoil without mechanical failure. Elastin functions as an entropic elastomer, whose properties are the direct result of the inability of the protein to fold into a fixed, stable structure.

Previous investigations of how the unusual properties of polymeric elastin arise from the sequence of tropoelastin have depended primarily on modeling using molecular biological and biophysical methodologies. This study takes a unique alternative approach, using a well-curated database of tropoelastin sequences from more than 80 representative species of Amniotes to identify characteristics that are conserved over more than 300 million years of evolution in order to provide assembly and conformational flexibility requirements of elastins.

Conserved characteristics included preservation not only of regions of linear or positional sequence, but also of collective or compositional characteristics, derived from the sequence but not strictly dependent on positional sequence. A plausible overall consensus sequence for Amniote tropoelastins allowed quantification of residue-by-residue, domain-by-domain and region-by-region levels of sequence conservation, identifying distinct regions of high and low positional sequence conservation. Regions of low positional sequence conservation nevertheless maintained a recognizable, low complexity sequence style characterized by tandem repeats and partial repeats of short, non-polar motifs. Analysis of these motifs indicated hPGhGG, with numerous mutations, insertions and deletions, as the underlying repeating unit in all Amniote tropoelastins.

Together these data identify significant evolutionary constraints dictated by fundamental requirements for formation and functionality of the extracellular elastin matrix. Mutations/polymorphisms in human tropoelastin affecting such well-conserved characteristics might be expected to have phenotypic consequences.

## 1. Introduction

Elastin is the remarkable polymeric protein that is responsible for the important physiological properties of extensibility and elastic recoil in vertebrate tissues. Polymeric elastin is particularly abundant in tissues such as large arteries, heart valves and lung parenchyma, all of which undergo frequent and rapid cycles of extension and recoil. Elastin is also an important, if less abundant and less studied component of a number of other tissues, including skin, periostium and pericardium, bladder, uterus, elastic tendons and cartilages, as well as Bruch’s membrane in the eye. There is good evidence for the very slow or even lack of turnover of polymeric elastin, especially in tissues such as skin, lungs and large blood vessels. This means that the same polymeric material laid down during development must maintain the ability to repeatedly and efficiently stretch and recoil over a lifetime of literally billions of cycles, without plastic deformation or more dramatic mechanical failure. For a recent general review of the biochemistry of elastin see Keeley^1^.

The property of energy-efficient extension and elastic recoil is unusual for proteins. Such properties, as well as the remarkable durability of polymeric elastin have attracted great interest in understanding how these properties arise from the sequence of its monomeric precursor, tropoelastin, and how that sequence guides the process of assembly and cross-linking of the monomeric subunits into the architecture of the polymer. Such interest comes both from the basic science point of view, as well as with an aim to generate artificial protein-based biomaterials that not only mimic the properties of native elastin but also can be modified by design to ‘tune’ the properties of the biopolymer.

Unlike collagen and many other structural proteins, elastin is a relative newcomer in evolutionary terms, first appearing in elasmobranchs (e.g. sharks and skates) about 450 million years ago, around the time of the appearance of closed circulatory systems. Recognizable elastin is absent from lampreys and hagfish (agnathans) and from other chordates close to the vertebrate/invertebrate boundary. Although it seems highly unlikely that elastin appeared *de novo* 450 million years ago, there has been no clear identification of possible ancestral proteins in lower chordates and invertebrates^2^.

Although the origin of the unusual elastomeric properties of polymeric elastin has been the subject of debate for many years, there is now a strong consensus that elastin functions as an entropic elastomer, achieving its elastomeric properties as the direct result of the inability of the protein to fold into a fixed, stable structure^3–8^. The consequence of this lack of fixed structure is that elastin, in both its monomeric and polymeric forms, can be best described as an ensemble of energetically similar, rapidly interchanging conformations with a flat energy landscape. In terms of the sequence of the protein, this appears to be achieved by an abundance of well-distributed, short and local sequence motifs rapidly exchanging between beta-turn and polyproline II conformations^6,7,9–13^.

Most investigations of the relationship between the sequence of elastin and its functional elastomeric properties have taken the approach of reproducing the sequences either of full-length tropoelastins, usually human, bovine or mouse, or generating polypeptides mimicking regions of such sequences. Specific alterations in these sequences are then made to assess effects on conformational ensembles, polymeric assembly and functional properties of the resulting biomaterials. Such investigations have led to many useful insights into sequence/structure/function relationships and have indicated that, at least in some cases, even single amino acid substitutions can substantially affect functional properties^13–22^.

An alternative and less-explored approach to understanding the contribution of sequence to functional properties of elastin is to exploit the fact that tropoelastins have been present and evolving in vertebrates for at least 450 million years. As a consequence, identification of characteristics that have been retained over that period of time would give a good indication of fundamental compositional and sequence elements that must be conserved in order to maintain both the conformational flexibility and the mechanisms of polymeric assembly that are common to all elastins.

In an earlier study, using a much smaller dataset of tropoelastin sequences, we identified regions of tropoelastins with remarkable positional sequence conservation across essentially the whole range of elastin evolution^2^. Other regions of the protein showed less direct sequence retention, but were characterized by a sequence ‘style’ consisting of extensive but variable and imperfect tandem repeats of short non-polar motifs. However, because of the fewer available elastin sequences and the wide range of species used, that study could only provide a relatively coarse-grained overview of either compositional or positional sequence characteristics.

Here, to obtain a finer definition of retained sequence characteristics, we have narrowed the range of species investigated to Amniotes (mammals, birds and other reptiles), while still encompassing more than 300 million years of evolution. This narrowing of the phylogenetic range simplifies the analysis because Amniote tropoelastins appear to constitute a clade, with easily identifiable correspondence of domains across species. At the same time, we have increased the number of Amniote species represented to approximately 80, including almost equal numbers of Synapsid (placental and non-placental mammals) and Sauropsid (birds and reptiles) species. This database provides sufficient coverage of evolutionary branches and sub-branches not only to identify universally retained sequence characteristics, but also to distinguish consistent sequence and domain alterations between sub-branches that might reflect evolutionary modulation of properties in response to diverse functional requirements.

## 2. Methodology

### 2.1 Collection and curation of tropoelastin sequences

Supplementary Table S1. contains a complete list of the species from which tropoelastin sequences were collected for this study. Species are grouped by Class: Mammalia (45 species), Aves (19 species), and Reptilia (22 species), for a total of 86 species. These groupings were further sub-divided by Order, Family, and finally individual species. For convenience, Supplementary Table S2 provides an alphabetized list of the three-letter species codes employed.

Figure 1 shows an overall view of the phylogenetic relationships among the species used. Supplementary Figures S1-S4 give more detailed outlines of the phylogenies of sub-groups of species. These can be useful to correlate with modifications in sequences and domain arrangements at the Order and Family level. Phylogenetic relationships are generated from a variety of sources^23–33^, but an excellent general source of such information can be found at the TimeTree5 website^34^ (http://www.timetree.org). Timings of species divergence are approximate. Note that the phylogenetic relationship of testudines (turtles) to other reptilian Orders reflects the more recent molecular genomics data of Field^27^ and Crawford^30^. Previous data, based on morphological characteristics, had suggested an earlier divergence of testudines from all other reptiles.

**Figure 1.**
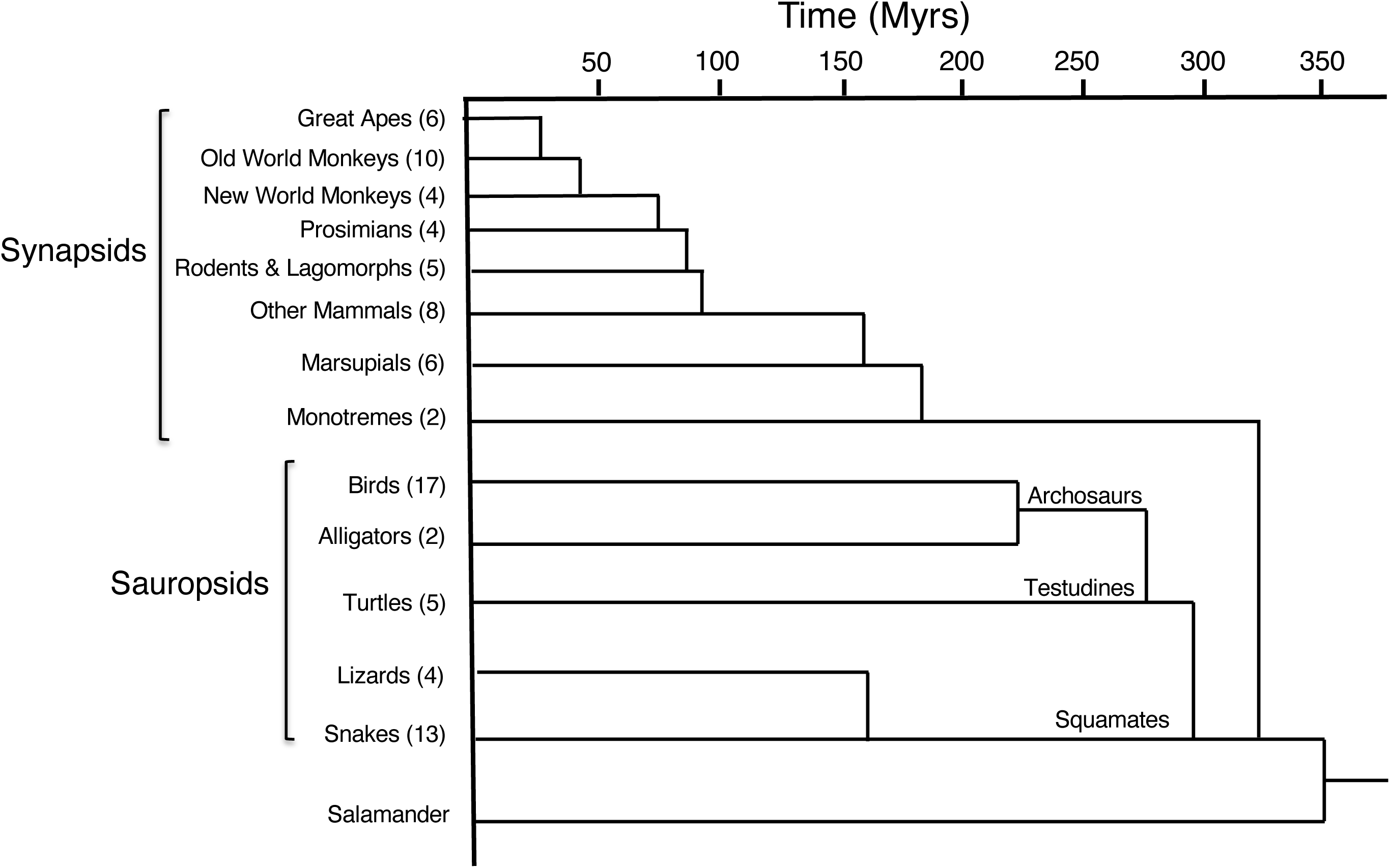
Overall evolutionary relationships among Amniotes, with amphibians (salamander) used as an outgroup. Data was compiled from Kumar et al.^34^. Synapsid and Sauropsid groupings are labelled. Numbers of individual species included in the curated database for each sub-group are indicated in parentheses. Dating of branch points is approximate. Detailed relationships within sub-groups are provided in Supplementary Figures S1-S4.

Most sequences were harvested from the US National Center for Biotechnology Information database (https://www.ncbi.nlm.nih.gov), with a smaller number sourced from the EnsemblGenomes database (http://ensemblgenomes.org). Species were chosen to give broad as possible coverage of Classes and Orders. Selected sequences generally had complete or essentially complete coverage of the coding sequence of tropoelastin, although sequences with more limited coverage were also used for domain-specific analyses.

Designations of genomic source sequences for each species are listed in Supplementary Table S1, together with those of any supporting sequences in the form of RefSeqs, ESTs and transcriptome sequence assemblies (TSAs). However, because of frequent alternate splicing of tropoelastins, such supporting RefSeqs, ESTs or TSAs will not necessarily include all exons present in the genomic sequence. In a few cases no genomic sequence could be found, but apparently complete TSA sequences were available.

In many cases the tropoelastin gene could be found simply by a search using the gene designation ‘eln’. As an alternative, searches using the genomic sequence of exons that were well-conserved across species (e.g. exons 2, 10, 15, 29/29a, 36) were usually successful, particularly for identifying tropoelastin sequences of more closely related species when Order or Family designations could be used to narrow the search parameters. Finally, because of the previously reported consistent conservation of synteny with the Lim kinase-1 gene^2^, unannotated tropoelastin genes could also be found as the next open reading frame upstream of a documented Lim kinase-1 gene, generally with a gap of approximately12,000 bases between these genes. Sequence data were current at the time of final curation. More recent updates or additions, including transcriptome sequence, may not be reflected in the sequence information.

Importantly, each sequence was individually curated, with curation notes summarized in Supplementary Table S1. In general, curation began with the genomic sequence, initially identifying exon boundaries by comparisons with supporting RefSeqs, ESTs and TSAs, as well as with previously identified elastin sequences, especially of species in the same Order or Family. Exon boundaries were also predicted using splice site prediction. The NNSplice v. 9.0 tool (http://www.fruitfly.org/seq_tools/splice.html) was generally successful in identifying and quantifying probabilities of donor and acceptor sites, but was not a good predictor of susceptibility to alternate splicing, at least in human tropoelastin^22^. Exon identification and alignments with genomic sequence using these several criteria were generally consistent, although a few exceptions are noted in Supplementary Table S1. In agreement with previous observations^2^, all splice sites in tropoelastins of all species were phase 1:1, facilitating cassette-like splicing events without altering the reading frame.

Genomic sequences sometimes included sequencing gaps, with nucleotide identifications replaced by Ns. Except in cases where this absent sequence could be recovered through supporting RefSeq, EST or TSA data, this region of sequence was designated as lost in a sequencing gap. In a few cases, an abundance of sequencing gaps, together with the generally repetitive nature of the sequence in some regions of tropoelastin, resulted in mis-assembly of the exons in the source database. Such mis-assemblies could be plausibly corrected using genomic sequence data from closely related species. If expected exons could not be identified from three-frame translations of genomic sequence in spite of the absence of evident sequencing gaps, and these exons were also absent from supporting RefSeq, EST or TSA data, the exon was designated as absent from the genomic sequence. Such absences were strongly supported if the same absence was predicted in a group of closely related species.

Alternate splicing is well-known to be common in tropoelastins, with at least 13 such isoforms identified in human tropoelastin^22^. Therefore, several possibilities had to be considered when sequence mutations at splice sites were identified that would significantly reduce the probability of viable splicing. In general we erred on the side of inclusion. Thus, if the exon sequence was present in at least some supporting RefSeq, EST or TSA data the exon was included. Similarly, if a single base substitution at the mutated site restored a strong probability for splicing, the apparent mutation was considered a sequence reading error, and the exon was included. However, in some cases the same or a similar splice site mutation was identified in several closely related species and the exon sequence appeared to be degenerating, suggesting that the exon was either recently lost from the protein-coding sequence or was in the process of being lost.

In general, all identifiable exons, including those with sequencing gaps or questionable splice sites, were included in the overall sequence tables. Rules governing inclusion or exclusion of such questionable sequences for the purpose of analysis of protein sequence data and motifs are described in detail elsewhere.

## 3. General observations

### 3.1 General characteristics and locations of elastin genes in Amniotes

The genomic data for most tropoelastins used in this database includes large regions both upstream of the translation start site and downstream of the stop codon. Such sequence data could provide a useful resource for identifying conserved sites of importance for the regulation of tropoelastin expression. However, no such analysis has been made here, with the exception of the recognition of significant regions of strong sequence conservation in the proximal 3’utr (see section 5.5). The database might also be a useful resource for investigations of variations in codon usage, both between species groups and between different regions of the sequences. However, no such analysis has been done.

A few general observations should be noted. First, there was no evidence for multiple tropoelastin genes in any Amniote species. This is consistent with earlier observations in which multiple tropoelastin genes could only be identified in teleost fish^2,35,36^, presumably the result of a genomic duplication event that occurred in the lineage leading to teleosts, as well as in some species of frogs^2,37^. Consistent with earlier observations^2^ and in spite of many differences in chromosomal locations among species, all tropoelastin genes remained syntenic with the Lim kinase-1 gene, with eln generally located approximately 12,000 bases upstream of Lim kinase-1.

Overall for Amniotes, tropoelastin protein-coding sequences comprised only a small proportion (approximately 6% on average) of total bases between the start of exon 1 and the stop codon. Intron 1 is consistently by far the largest intron in the gene, including 7,000-17,000 bases, depending on the species. In contrast, some introns in the replicated regions of elastin sequences can be as short as 60-100 bases. As noted above, all intron/exon borders are phase 1:1.

### 3.2 Domain characteristics of Amniote tropoelastins

For clarity, the term ‘exon’ will refer to the DNA sequence and the term ‘domain’ will refer to the protein product of that sequence. A schematic diagram of the domains of Amniote tropoelastins is shown in Figure 2. Each domain consists of the protein product of a single exon.

**Figure 2.**
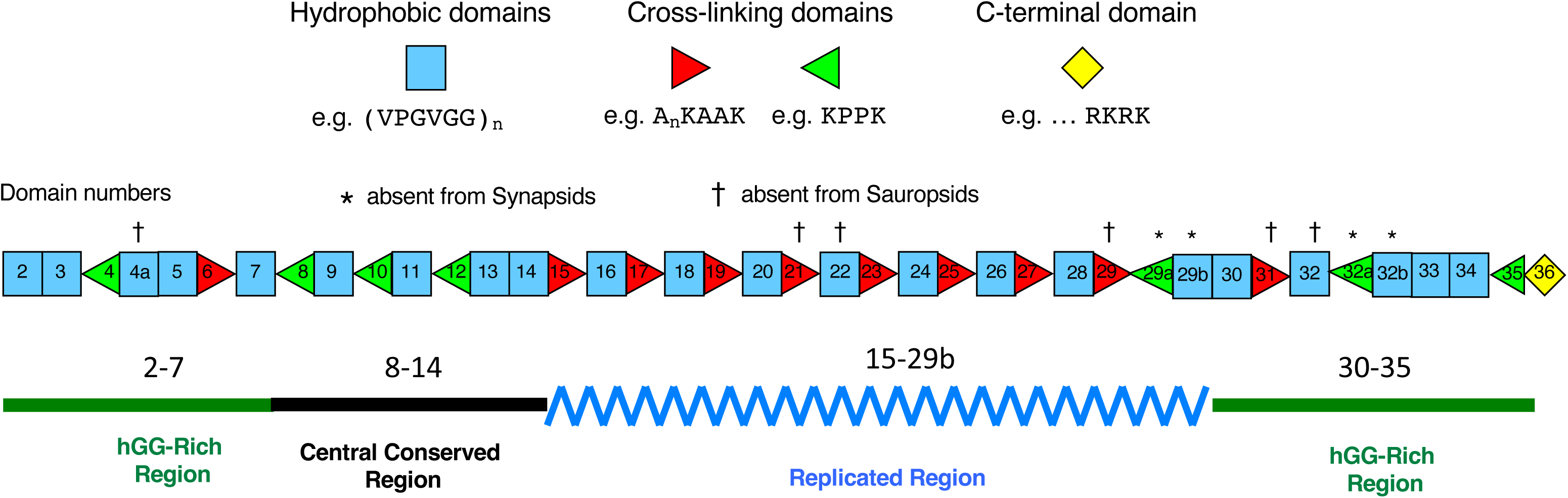
Schematic diagram of ‘domains’ and ‘regions’ of Amniote tropoelastins, highlighting their sequence characteristics. Each domain corresponds to the protein product of a single exon. Domain 1 (exon 1) is excluded because it contains only the signal peptide, which is cleaved from the mature protein sequence. Hydrophobic domains are rich in non-polar amino acids, often present in tandem repeats and pseudo-repeats. Cross-linking domains contain the lysine residues that are subsequently involved in covalent cross-linking of the tropoelastin monomer into polymeric elastin. Cross-linking exons are either KA or KP types, depending on the nature of the amino acid sequences flanking/interspersing the lysine residues. While most domains/exons are common to Synapsid and Sauropsid tropoelastins, exons 4a, 21, 22, 29, 31 and 32 (indicated by †) are consistently absent from the genomic sequences of Sauropsid tropoelastins. Conversely, domains/exons 29a, 29b, 32a and 32b (indicated by *) are absent from the genomic sequences of Synapsid tropoelastins. Domain/exon 4a is present only in a sub-set of Synapsid genomic sequences. The sequence of the Central Conserved Region (domains 8-14) is strongly conserved across all tropoelastins. The sequence of the Replicated Region (domains 15-29b) appears to be generated by the replication of pairs of cross-linking and hydrophobic exons, with subsequent mutations. The flanking hGG-rich Regions are generally rich in hGG motifs, where ‘h’ is a non-polar amino acid, especially V, I, L, A or G. Domain 36, the C-terminal domain, has a well-conserved sequence including the only two cysteine residues in tropoelastin, and consistently terminating with a tetrabasic motif (RKRK).

Domain 1 is not included in Figure 2 because it contains only the signal sequence, except for a single amino acid (glycine) that remains at the N-terminal of the processed protein after signal peptide cleavage. Signal peptide cleavage sites were predicted using SignalP-6.0 (https://biolib.com/DTU/SignalP-6/), using the protein sequence of the first 4-5 domains of tropoelastins, and choosing the Eukarya option.

Domain 36 has a unique, well-conserved sequence that includes the only two cysteine residues in the protein, and consistently terminates with a tetrabasic motif (RKRK).

Other domains in Amniote tropoelastins can be generally categorized as either hydrophobic or cross-linking. Hydrophobic domains are rich in non-polar amino acids (predominantly V, I, L, A, G and P), usually present in tandemly repeated motifs or quasi-motifs. Detailed analyses of these motifs are presented below (see section 6). Cross-linking domains contain the lysine residues that subsequently participate in covalent cross-linking of the tropoelastin monomer into polymeric elastin. Cross-linking domains are either KA or KP types, depending on the nature of the amino acids flanking/interspersing the paired lysine residues.

Domains 6, 13, 14 and 36 provide examples of exceptions to this simple categorization of hydrophobic and cross-linking domains. Domain 6 is classed as a cross-linking domain because it contains a typical lysine pair, although that cross-linking motif is preceded by an unusually extended non-polar sequence. In contrast, domains 13 and 14, consisting of well-conserved, predominantly non-polar sequences, are classed as hydrophobic domains, although each includes a single lysine residue that likely participates in at least some cross-linking reactions. Although domain 36 contains a single lysine residue (in addition to the terminating RKRK motif), there is no current evidence that this lysine singlet participates in crosslinking. These exceptions are consistent features in all Amniote tropoelastin sequences. For more detailed information on polymeric assembly and cross-linking of tropoelastin see Keeley^1^.

### 3.3 Regions common to all Amniote tropoelastins

On the basis of previous sequence information^2^ we had identified five distinct regions (excluding the signal peptide) in the sequence (Figure 2). These regions were also consistently recognizable in all tropoelastins used in this study. The ‘central conserved region’ (domains 8-14) comprises approximately150 amino acids and contains a sequence that is extremely well-conserved across all Amniotes (see section 5.2). Downstream of the central conserved region is the ‘replicated region’ (domains 15-29b), characterized by alternating cross-linking and hydrophobic domains. As the name suggests, the nature of the sequences present in this region strongly suggests that they were generated through multiple replications of a hydrophobic/crosslinking pair of exons, followed by extensive evolutionary sequence diversification through mutation, especially in hydrophobic domains. Upstream of the central conserved region and downstream of the replication region are ‘hGG-rich’ regions (domains 2-7 and 30-35, respectively), so-named because the hydrophobic domains of the region contain a relative abundance of hGG motifs, where ‘h’ is predominantly valine, isoleucine, and leucine (see section 4.4).

## 4. Conservation of bulk or collective properties of Amniote tropoelastins

For most proteins, considerations of conservation across evolutionary time scales focus on retention of sequence regions that are directly associated with basic structural elements, e.g. helices, turns and sheets, as well as other more complex structural motifs. Such proteins fold into specific stable structures that represent states of minimum energy. In contrast, tropoelastin is an intrinsically disordered, low complexity protein, best described as an ensemble of energetically similar, rapidly interchanging conformations. Both self-assembly of tropoelastin into its polymeric state, a process involving liquid-liquid phase separation (coacervation), and the entropic elastomeric properties of polymeric elastin are absolutely dependent on maintenance of a high level of structural/conformational disorder, even in its polymeric state.

Like other low complexity, intrinsically disordered proteins, tropoelastins are characterized both by conservation of positional sequence as well as by conservation of bulk or collective properties^38,39^. Collective properties are derived from amino acid composition but are not strictly dependent on linear positional sequence. Consequently, both modes of conservation must be considered in assessing characteristics of tropoelastins retained over evolutionary time scales as essential for fundamental functional properties of elastin.

### 4.1 Amino acid compositions of Amniote tropoelastins

Amino acid compositions corresponding to the various Amniote tropoelastins were generated from sequence data after removal of the signal peptide (see section 3.2). All predicted cleavage sites were of high probability. Signal peptide sequences were generally well-conserved across all Amniote species (see section 6.4), and in all cases the cleavage site was predicted to be located between the last two amino acids coded for in exon 1, resulting in the addition of a glycine residue at the first amino acid position of the mature protein after cleavage. The addition of this glycine residue to the mature protein sequence was also assumed for species for which no genomic or protein sequence for exon 1 was available.

Amino acid compositions, as well as other general compositional characteristics of the proteins (molecular weight, isoelectric point and grand average of hydropathicity (GRAVY)) were calculated using the EXPASY ProtParam tool (https://web.expasy.org/protparam/). In most cases, amino acid composition data was only calculated for tropoelastin sequences that were judged to be full-length, keeping in mind that exons in some species groups were absent from the genomic sequence. Tropoelastin sequences with incomplete domains because of sequencing gaps were generally not used for amino acid analyses, except in cases where only a single or at most two non-sequential domains were affected, and the sequence of these domains could be plausibly predicted from sequence conservation of the corresponding domains in closely related species. In the absence of transcriptome or direct protein sequence data, compositional calculations for species with unexpected additional exon replications in the genomic sequence (see section 7.7), took into account only a single replicate of these domain pairs. Post-translational conversion of some proline residues to hydroxyproline (see section 4.2) was not taken into account in the compositional calculations.

Species were grouped into data sets in accordance with the evolutionary relationships shown in Figure 1. ‘All Primates’ consisted of great apes, old world monkeys, new world monkeys and prosimians. ‘Other Mammals’ consisted of rodents and lagomorphs, and all other placental and non-placental mammals. ‘Archosaurs’ included birds and crocodiles/alligators. ‘Testudines’ included all turtles. ‘Squamates’ consisted of lizards and snakes.

The proportional distributions of amino acids (residues/100) for these data sets are plotted in Figure 3a. Detailed averaged compositional data are shown in Supplementary Table S3. In general, amino acid proportions are remarkably invariant across all species groups. The relatively limited selection of amino acids, characteristic of low complexity proteins, is evident across all tropoelastins, with a dominance of glycine, alanine, proline and valine. At the same time, there is a consistent paucity of cysteine residues (invariably present only in exon 36) and negatively charged amino acids (aspartic and glutamic acids), as well as a complete absence of histidine and tryptophan. Methionine residues are also absent from the mature protein in all species, with the exception of some, but not all, marsupials, which include a single methionine residue in domain 36 (see opossum in Supplementary Table S11 for an example). The fact that a methionine residue appears independently at this site in more than one marsupial species suggests that it is not the result of a sequencing error.

**Figure 3.**
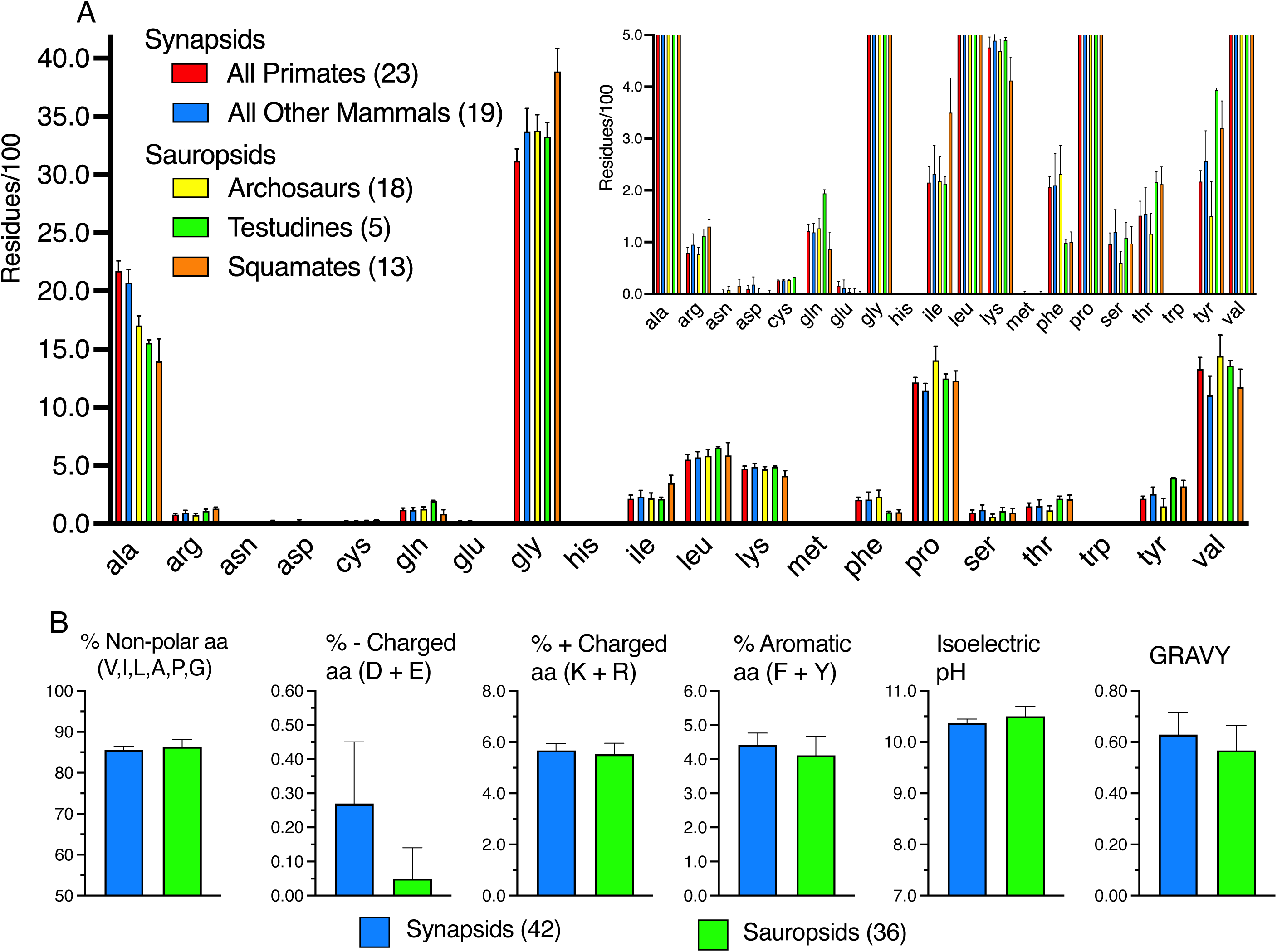
Comparison of proportional amino acid compositions across all Amniote species. **A**: A total of 78 amino acid compositions of tropoelastins across amniote species were plotted as residues/100. Five species groups were used, grouped according to evolutionary relationships (Figure 1). These consisted of All Primates (great apes, old world monkeys, new world monkeys and prosimians), ‘All Other Mammals’ (rodents, lagomorphs, other mammals, marsupials and monotremes), ‘Archosaurs’ (birds and crocodiles/alligators), ‘Testudines’ (turtles) and ‘Squamates’ (lizards and snakes). The inserted plot expands the y-axis to better display low-abundance amino acids. Values are plotted as mean and standard deviation. **B**: Comparison of collective properties of tropoelastins, averaged over Synapsid and Sauropsid sub-groups. Parameters include proportions of non-polar (V, I, L, A, P, G), negatively charged (D, E), positively charged (K, L), and aromatic (F, Y) amino acids, as well as isoelectric pH and GRAVY (Grand Average of Hydropathicity). Data are plotted as mean ± standard deviation. See Supplementary Table S3 for additional details.

It has been previously suggested that evolutionary selection against negatively charged amino acids or generally distributed cysteine residues is likely related to minimization of opportunities for salt bridges and disulphide bond formation, either of which would introduce stable, non-local structural elements^1,2^. Avoidance of methionine residues may be related to the general lack of turnover of polymeric elastin and the fact that this amino acid may be oxidized over time, resulting in the generation of a negatively charged side chain. The reason for the consistent absence of histidine and tryptophan is not clear.

Overall compositional properties are shown in Figure 3b, comparing averaged Synapsid and averaged Sauropsid data. Parameters include total proportions of non-polar amino acids (defined here as glycine, alanine, proline, valine, leucine and isoleucine), proportions of amino acids with negatively charged side-chains (aspartic and glutamic acids), positively charged side-chains (lysine and arginine) or aromatic side-chains (tyrosine and phenylalanine). Other general properties compared were isoelectric pH and grand average of hydropathy (GRAVY), a measure of overall hydropathy of the protein.

These data show a remarkable preservation of collective character over the hundreds of millions of years of evolution separating Synapsids and Sauropsids. Conservation was particularly notable in the proportions of non-polar amino acids, as well as amino acids with positively charged or aromatic sidechains. While the averaged proportion of amino acids with negatively charged sidechains was very small in both Synapsid and Sauropsid species groups (< 0.3 percent), such amino acids were particularly rare in Sauropsid tropoelastins. Indeed, aspartic acid and glutamic acid residues were completely absent from 26 of 36 species of Sauropsids, compared to 4 of 42 species of Synapsids. This was reflected in a small but significant increase in the averaged isoelectric pH of Sauropsid compared to Synapsid tropoelastins (Figure 3b).

The remarkably strong conservation of these compositional parameters between Synapsid and Sauropsids suggests that they are important for the fundamental requirement of tropoelastins for polymeric self-assembly and maintenance, even in the polymeric state, of the degree of conformational disorder necessary for the protein to function as an entropic elastomer. At the same time, small but significant differences in these parameters between species groups may reflect group/species-specific adaptations in functional properties required by environmental or other considerations.

An interesting possible example of such adaptations may be seen in whales and dolphins, whose tropoelastin compositions include unusually high proportions of acidic amino acids (0.7- 0.8 %). As noted above, evolutionary selection against acidic amino acids is likely related to minimizing opportunities for formation of stable, non-local salt bridges that would disfavour the conformationally disordered state of the protein and affect the elastic modulus of the elastin polymer. In this respect, large arteries of deep-diving mammals have been reported to be significantly stiffer than those of other mammals, presumably as an adaptation to substantial increases in transmural pressures in diving^40^.

### 4.2 Proline hydroxlation

As noted above, because compositional properties were calculated from translated genomic sequences, post-translational hydroxylation of proline residues to 4-hydroxproline were not taken into account in compositional data. Hydroxylation of proline residues in elastin preparations was initially thought either to be due to contamination with collagen, or to incidental activity of prolyl hydroxylase. However, recent mass spectrometry data has shown that such hydroxylation is both tissue- and species-specific^41^. Furthermore, there is now evidence that such hydroxylation may have important consequences for the properties of the protein^42^. Using a mass spectrometry approach, Schmeltzer and his colleagues^41^ have demonstrated that hydroxylation of proline residues in tropoelastin takes place in the context of GxPG motifs, where x is V, L, I, A or G, with GIPG and GLPG motifs showing the most consistent incidence of hydroxylation, at least in the several mammalian species they examined.

The percentage of proline residues occurring in the context of such potential hydroxylation motifs, calculated for a total of 82 Amniote tropoelastin sequences, is shown in Supplementary Figure S5. Averaged over all Amniotes, prolines in GxPG contexts accounted for approximately 26% of all prolines, a number which is consistent with the levels of proline hydroxylation determined by Schmelzer^41^. When the ‘x’ residue was restricted to I or L, that proportion decreased to approximately 7%. Data averaged over Synapsid or Sauropsid sub-groups were essentially identical for both general GxPG motifs and for more restricted G(L/I)PG motifs (Supplementary Figure S5A). However, when species sub-groups were compared (Supplementary Figure S5B), it became clear that there were highly significant differences especially among Sauropsid sub-groups, perhaps reflecting increased diversity of functional adaptations in non-mammalian elastins.

### 4.3 Proline residues, proline/glycine ratios and proline-glycine pairs

A characteristic of all Amniote tropoelastins is the presence of a large proportion of proline (P, ∼12.5%) and glycine (G, ∼33.9%) residues (Supplementary Table S3). Consistent with earlier reports using more limited species data^2^, most of these prolines are immediately followed by a glycine residue (Figure 4A). This proportion (∼72 ± 5) is consistent across all Amniotes, although small but sometimes significant differences are apparent among sub-groups of Synapsids and Sauropsids.

**Figure 4.**
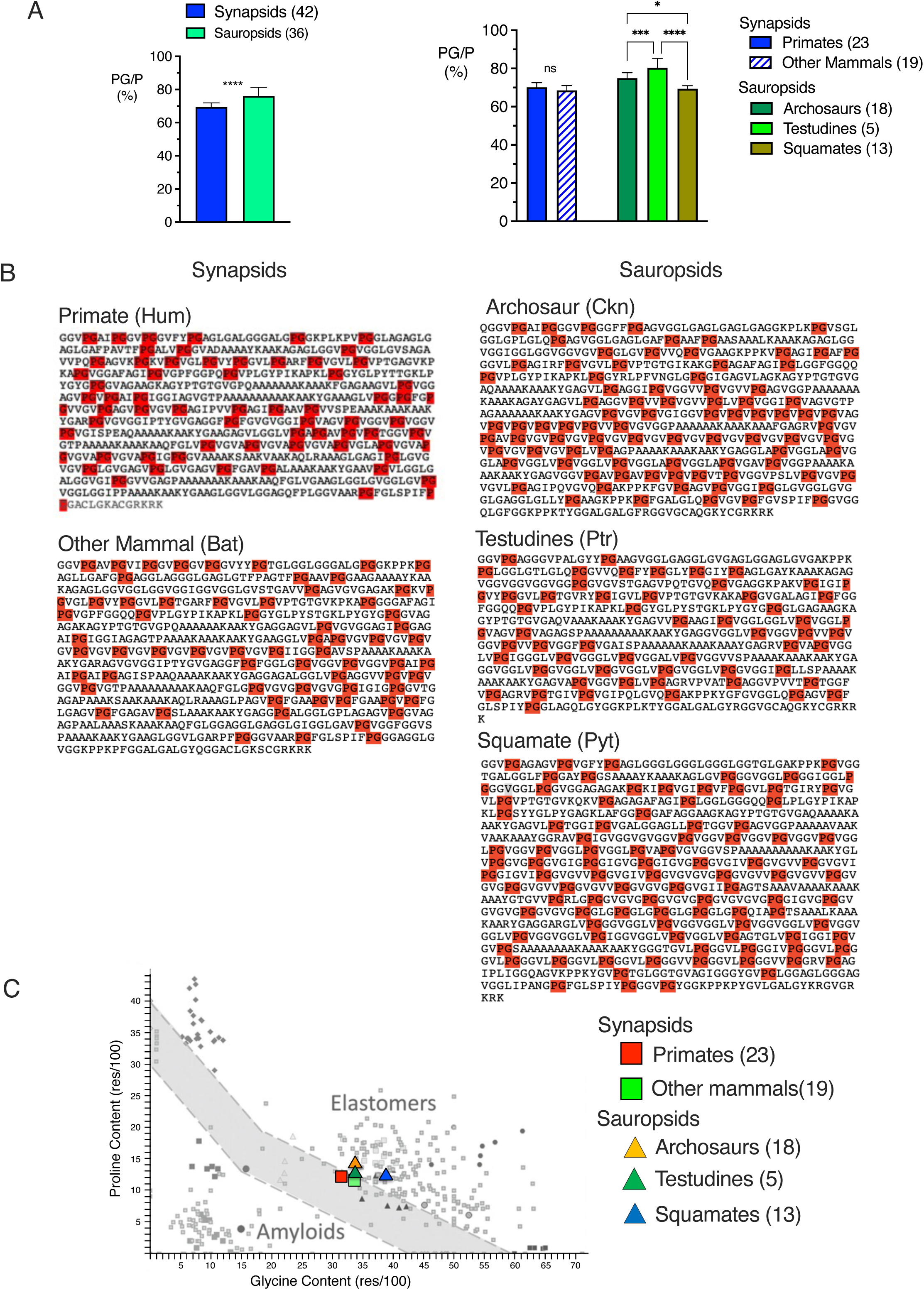
Abundance and distribution of proline-glycine pairs and ratios in Amniote tropoelastins. **A**: Proportions of proline residues in tropoelastin sequences that are immediately followed by a glycine residue (i.e. PG pairs). Data are compared between Synapsids and Sauropsid groupings (left), as well as between major sub-groupings of Synapsids or Sauropsides. Data are plotted as mean ± standard deviation. Statistical analysis of data pairs used an unpaired t-test with Welch’s correction for unequal variances. Multiple data sets were compared using a one-way ANOVA, with a Tukey correction for multiple comparisons. * p<0.5; *** p<0.001; ****p<0.0001. **B**: PG pairs (highlighted in red) are well-distributed across all tropoelastin sequences. Data shown are for representative species from major sub-groupings of Synapsids and Sauropsids. **C**: Proline to glycine relationships in tropoelastin sequences of Amniotes averaged over major sub-groupings of Synapsid and Sauropsid species. Data are superimposed on the original P/G relationship published previously by Rauscher et al.^43^ for polypeptides with amyloid-forming and elastomeric properties. Polypeptides falling within the ‘elastomer’ region of the original Rauscher plot included tropoelastin domains, resilin, mussel byssus, and flagelliform spider silks. Polypeptides falling within the ‘amyloid’ region included amyloids, rigid egg-shell proteins and tubilliform and aciniform spider silks. See Rauscher et al.^43^ for more details. For all species groupings, the averaged data from Amniote tropoelastins were tightly grouped in the elastomer region of the Raucher P/G diagram.

Such PG pairs are not only abundant in all tropoelastins but are also consistently well-distributed over the entire sequence of the protein, with the hydrophobic domains of the ‘replicated region’ of tropoelastins particularly rich in these PG pairs. Distributions of PG pairs for representative species of the major sub-groups of Synapsids and Sauropsids are shown in Figure 4B. Both NMR and molecular dynamics analyses have identified these PG pairs as a major site of the local and transient beta-turns that are important contributors to the conformational instability of tropoelastins^5–7,9–13,43–45^.

Rauscher and her colleagues, using a molecular dynamics approach, reported that, for a number of non-polar structural proteins, proline and glycine compositions were related to peptide backbone hydration, and that a simple threshold of P/G content could distinguish sequences of self-assembling, hydrophobic polypeptides that remained in a hydrated state with conformational flexibility and elastomeric properties from those undergoing hydrophobic collapse to form amyloid-like structures^43^. As shown in Figure 4C, proline/glycine ratios in Amniote tropoelastin sequences, averaged over the major sub-groups of Synapsids and Sauropsids, clustered tightly in the threshold region of the proline vs. glycine plot previously identified as characteristic of elastomers^43^.

### 4.4 Differential distribution of PG and GG pairs across domains/regions of Amniote tropoelastins

As noted above (Figure 4C), PG pairs, representing a site of formation of transient beta-turns, are both abundant and well-distributed across all Amniote tropoelastins. Earlier observations also noted the presence of well-distributed GG residue pairs in tropoelastins. Regions containing high densities of GG pairs have been suggested to result in a greater propensity for formation of short beta-sheets, providing regions of at least partial structural order^46–51^. Indeed, it has been demonstrated, at least *in vitro*, that replacement of PG pairs with GG pairs in elastin-like polypeptide sequences results in increased structural stability, and even in the formation of amyloid-like aggregates^52–55^. Thus, PG and GG densities, and the ratio of PG to GG pairs are likely a factor in achieving the balance between order and disorder that has been suggested to be a necessary characteristic of elastin-like proteins.

Relative densities of PG and GG pairs were calculated for four of the regions of Amniote tropoelastin sequence shown schematically in Figure 2. These included domains 2-7, domains 8- 14 (identified as the ‘central conserved region’), domains 15-29b (identified as the ‘replicated region’) and domains 30-35. Densities were measured as the number of each of these pairs present in the region, divided by the total number of amino acids in that region.

GG/PG density ratios for these four regions are plotted in Figure 5. Figure 5A shows ratios averaged across all Synapsid species, as well as over Primate and All Other Mammals sub-groups. Similarly, Figure 5B shows GG/PG density ratios averaged across all Sauropsid species, as well as over Archosaur, Testudines and Squamate sub-groups. For simplicity, only non-significant comparisons are indicated. Differences among all other groups are statistically significant.

**Figure 5.**
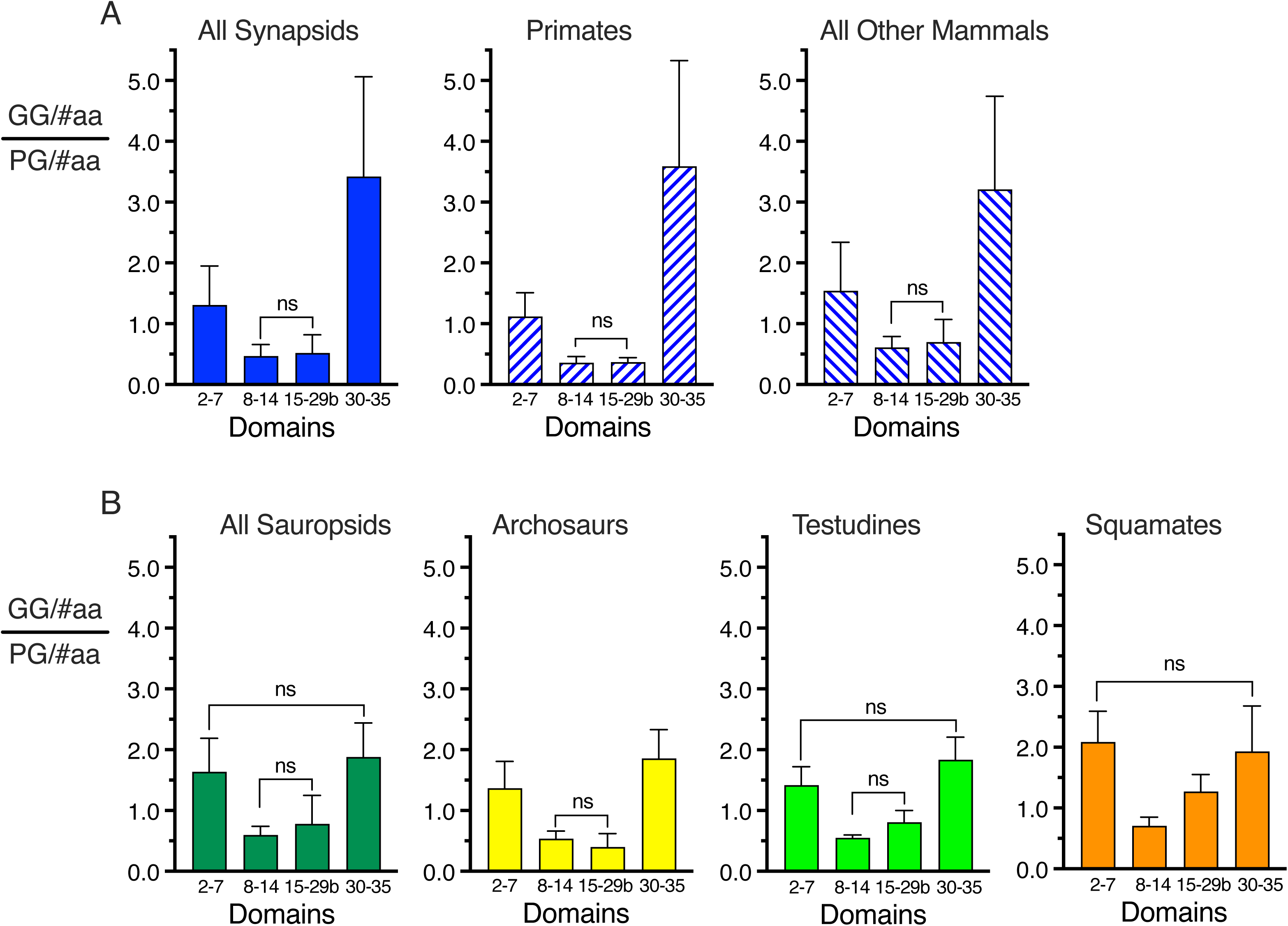
Differential relative densities of GG and PG pairs across sequence regions of Amniote tropoelastins. Densities were measured as the number of each of these pairs present in the region, divided by the total number of amino acids in that region. **A**: Comparison of ratios of GG/PG densities averaged across all Synapsids, as well as across Primates and All Other Mammals sub-groups of Synapsids. **B**: Comparison of ratios of GG/PG densities averaged across all Sauropsids, as well as across Archosaur, Testudines and Squamate sub-groups of Sauropsids. Data are plotted as mean ± standard deviation. See Figure 1 for details of evolutionary relationships among these groups and sub-groups. For simplicity, only non-significant comparisons are indicated. Differences among all other groups are statistically significant. For all Amniotes and sub-groups of Amniotes, PG densities predominated over GG densities (GG/PG ratios <1) in domains 8-14 and domains 15-29b, whereas GG densities predominated over PG densities (GG/PG ratios >1) in domains 2-7 and 30-35.

In all cases PG densities predominated over GG densities (GG/PG ratios were <1) in domains 8-14 and domains 15-29b, whereas GG densities predominated over PG densities (GG/PG ratios were >1) in domains 2-7 and 30-35. These differences were evident across all species and all sub-groups of species. GG/PG densities were notably high and showed greater variability for domains 30-35 as compared to domains 2-7, particularly in Synapsids and Synapsid sub-groups (panels A and B). The high variability in domains 30-35 of Synapsids was due to consistently elevated GG densities in prosimian species included in the Primate sub-group, as well as similarly consistently elevated GG densities in marsupial and monotreme species included in the All Other Mammals sub-group. Note that smaller but significant differences in GG/PG ratios between domains 2-7 and domains 30-35 were seen in the Archosaur sub-group of Sauropsids, but not for Testudines or Squamate sub-groups.

These data would suggest that, for all species, sequences in domains 2-7 and 30-35 may be relatively more susceptible to formation of beta-sheet structures, affecting the balance of conformational order and disorder in different regions of tropoelastins, a feature likely important for the process of oligomeric self-assembly as well as for elastomeric properties of the assembled polymer. Moreover, subtle modifications of this balance, through evolutionary manipulation of GG/PG ratios in these regions, would provide a mechanism for modulating tropoelastin characteristics in response to specific species or species group requirements. Indeed, Savage and Gosline reported decreased GG/PG densities in sequences of elastomeric spider silks as compared to silks with more rigid structural characteristics^56^.

Both PG and GG pairs in tropoelastin sequences are almost always preceded and followed by a non-polar amino acid, i.e. h_1_PGh_2_ or h_1_GGh_2_, where h_1_ and h_2_ are predominantly V, I, or L. More detailed information on preferential evolutionary choices for these non-polar residues is provided in section 6.1. The preponderance of hGG motifs over hPG motifs in domains 2-7 and 30-35 supports designating these regions as ‘hGG-rich’ in Figure 2.

The fact that such differential distribution of these parameters is conserved across all amniotes tropoelastins suggests that they represent important collective properties that must be retained to conserve the functional properties of elastins.

## 5. Regions of high positional sequence conservation in Amniote tropoelastins

While conservation of collective characteristics such as those described above are important for the fundamental properties of tropoelastins, it was also clear from the data that at least some regions of tropoelastins are characterized by significant conservation of linear/positional amino acid sequence. Regions of strong positional sequence conservation across an extended evolutionary period may, for example, represent sites of interactions with other matrix proteins involved in directing assembly of elastin into a polymeric architecture important for functional properties^1,57–65^. In order to identify and quantify regions of positional sequence conservation in tropoelastins it was first necessary to construct a consensus alignment for the sequences. Consensus sequences were constructed both for all Amniotes (∼ 80 species), as well as separately for sub-sets of Synapsids (46 species) and Sauropsids (36 species)

### 5.1 Alignment of tropoelastin sequences to produce consensus sequences

Plausible consensus sequences were generated by domain-by-domain alignment of individual sequences. Domain alignment was done manually, with the underlying assumptions of an evolutionary relationship among all tropoelastin sequences and identifiable domain correspondence across species.

The process began with the alignment of sequences of closely related species, e.g. great apes, followed by the stepwise addition of sequences of other primates, rodents, other mammals, marsupials, monotremes, and finally archosaurs (birds and alligators), testudines (turtles and tortoises) and squamates (lizards and snakes), in accordance with the evolutionary relationships shown in Figure 1. Alignments were adjusted at each step, introducing any gaps required to bring sequences into a plausible overall alignment. For many domains this was a relatively straight-forward process, since the domains were relatively short and the sequence alterations were, for the most part, due to easily recognizable patterns of conservative substitutions, as well as replication or deletion of complete or incomplete sequence motifs.

Alignment of hydrophobic domains, especially those in the ‘replicated region’ (domains 16-28) was more challenging. Such domains appeared to consist largely of semi-repetitive arrays of hPGh or hGGh motifs (where h was predominantly V, I, L, A, or G), with multiple insertions and deletions. These domains were both larger and more variable in size, even between closely related species, with size differences mostly due to variable numbers of repeats and partial/modified repeats. As a consequence, a modified strategy for alignment across such a wide range of species was required.

Alignments of these domains were achieved by using hPG and hGG motifs as ‘anchor points’ in the sequence, introducing gaps as necessary for alignment of these motifs. Table 1 provides simplified examples of representative domain alignments for a small number of selected but evolutionary diverse species. Because of the unusual character of tropoelastin sequences, with their abundance of motif replications, gaps, deletions and insertions, this manual process resulted in a more plausible consensus sequence than that which could be generated using any readily available alignment software.

**Table 1.**
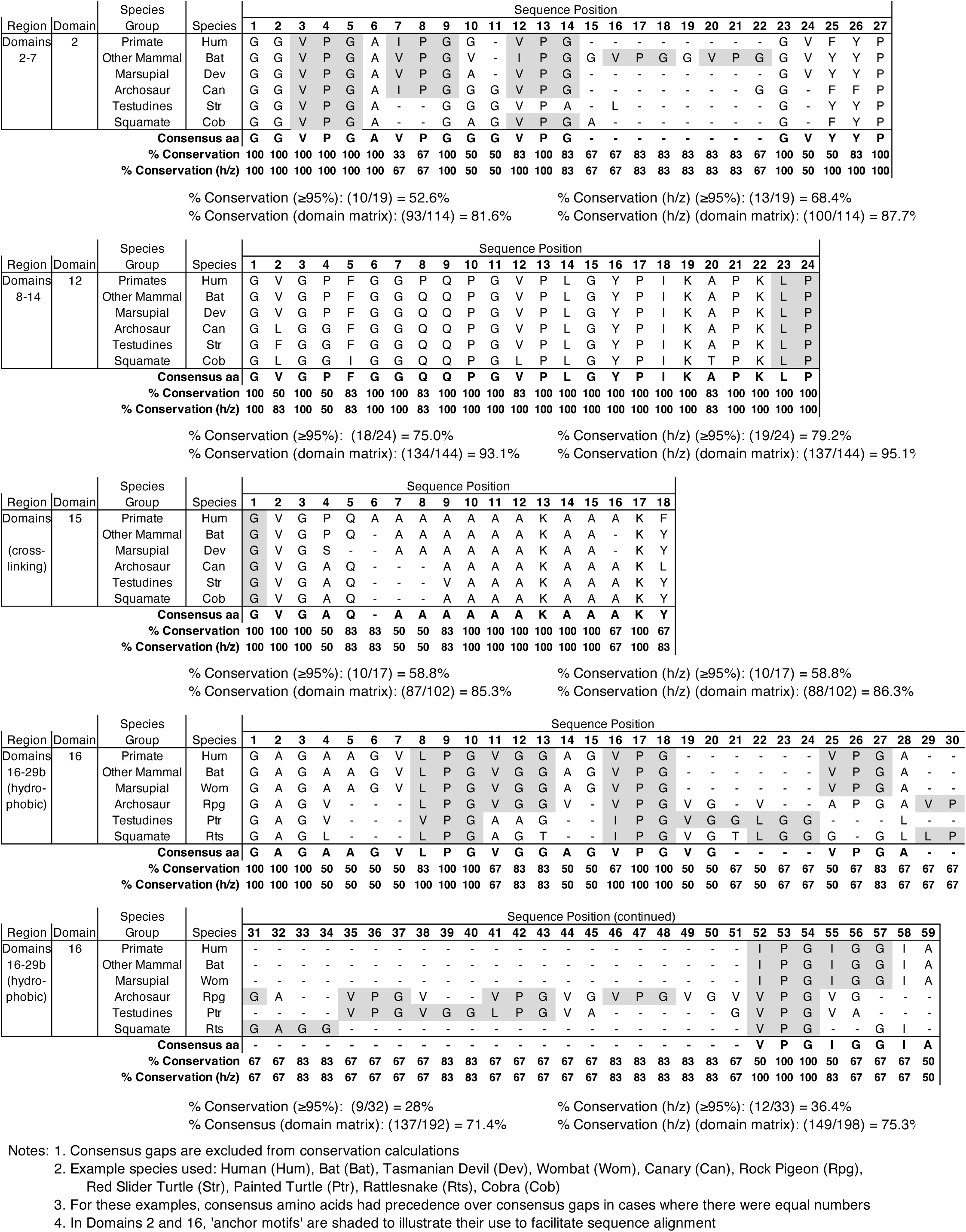
Examples of Domain Alignments and Consensus Calculations.

### 5.2 Calculation of positional conservation parameters from consensus sequence alignments

Several parameters were used to quantify the degree of positional sequence conservation across Amniote tropoelastins. Examples illustrating how these parameters were calculated are also provided in Table 1.

The ‘% Conservation’ parameter was calculated in the following way. The most frequently occurring amino acid at any given position in the aligned domain sequence was taken as the consensus amino acid at that position. The % Conservation for any sequence position was the proportional frequency of the consensus amino acid at that position. Note that, because of extensive sequence insertions and deletions, consensus gaps occurred at several sequence positions, especially in hydrophobic domains. In this way, the % Conservation parameter provided a residue-by-residue measure of positional sequence conservation, not only across domains but also across the entire Amniote consensus sequence.

Because of frequent inter-substitution of valine, isoleucine and leucine residues, especially in hPG and hGG motifs, these amino acids were also counted together as ‘h’. Similarly, phenylalanine and tyrosine residues were together counted as ‘z’, thereby generating a consensus parameter designated ‘% Conservation (h/z)’. This % Conservation (h/z) parameter, plotted residue-by-residue across the entire consensus sequence of Amniote tropoelastins, is shown in Figure 6. For convenience of plotting, consensus gaps were excluded from the plotted sequence, although the number of consensus gaps inserted and their locations in the consensus sequence were indicated on the plots. Sites of consensus hPG motifs and PG pairs, hGG motifs and GG pairs, KP and KA cross-linking motifs and singlet K residues, as well as consensus ‘z’ positions are indicated by color-coding on the plot.

**Figure 6.**
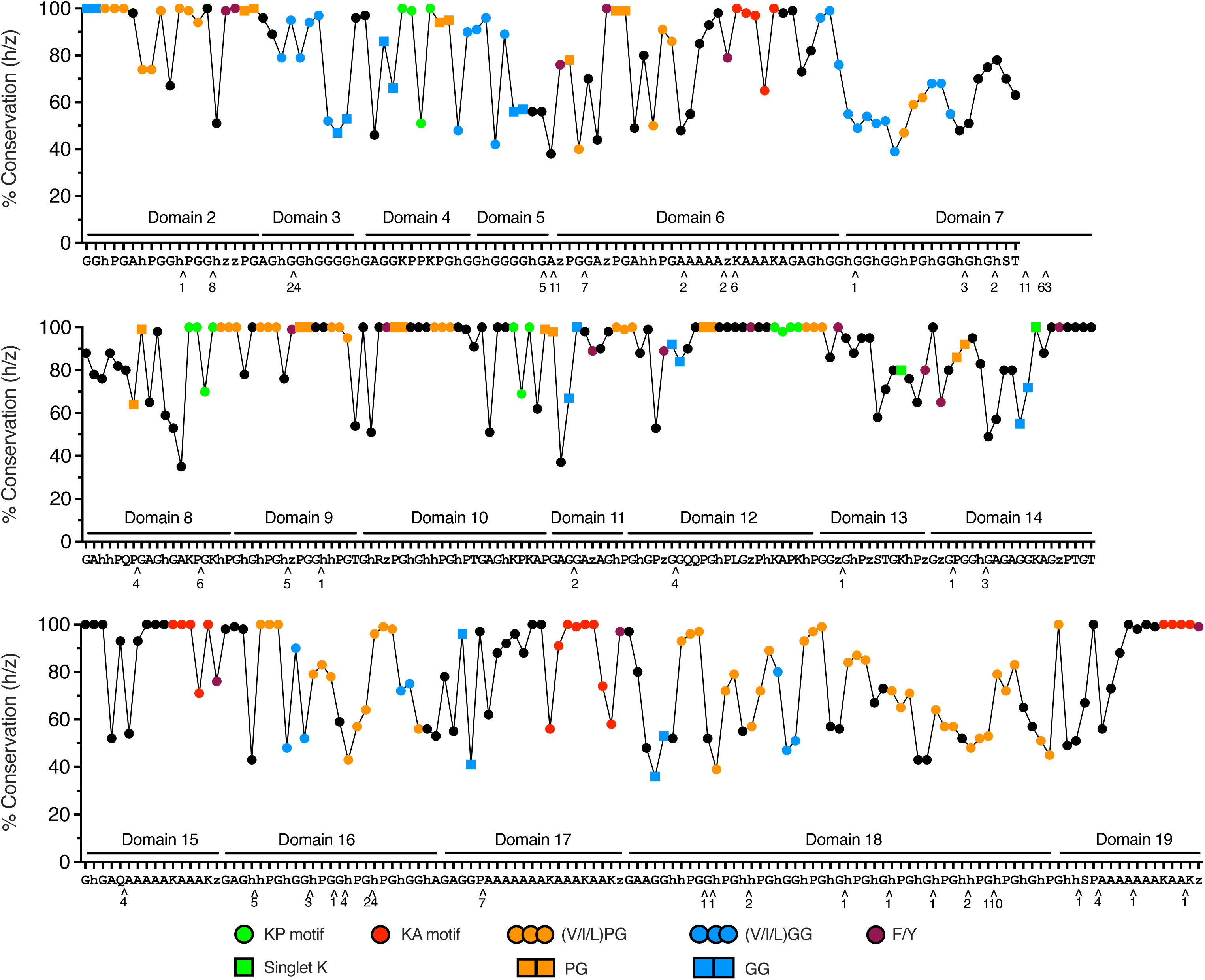
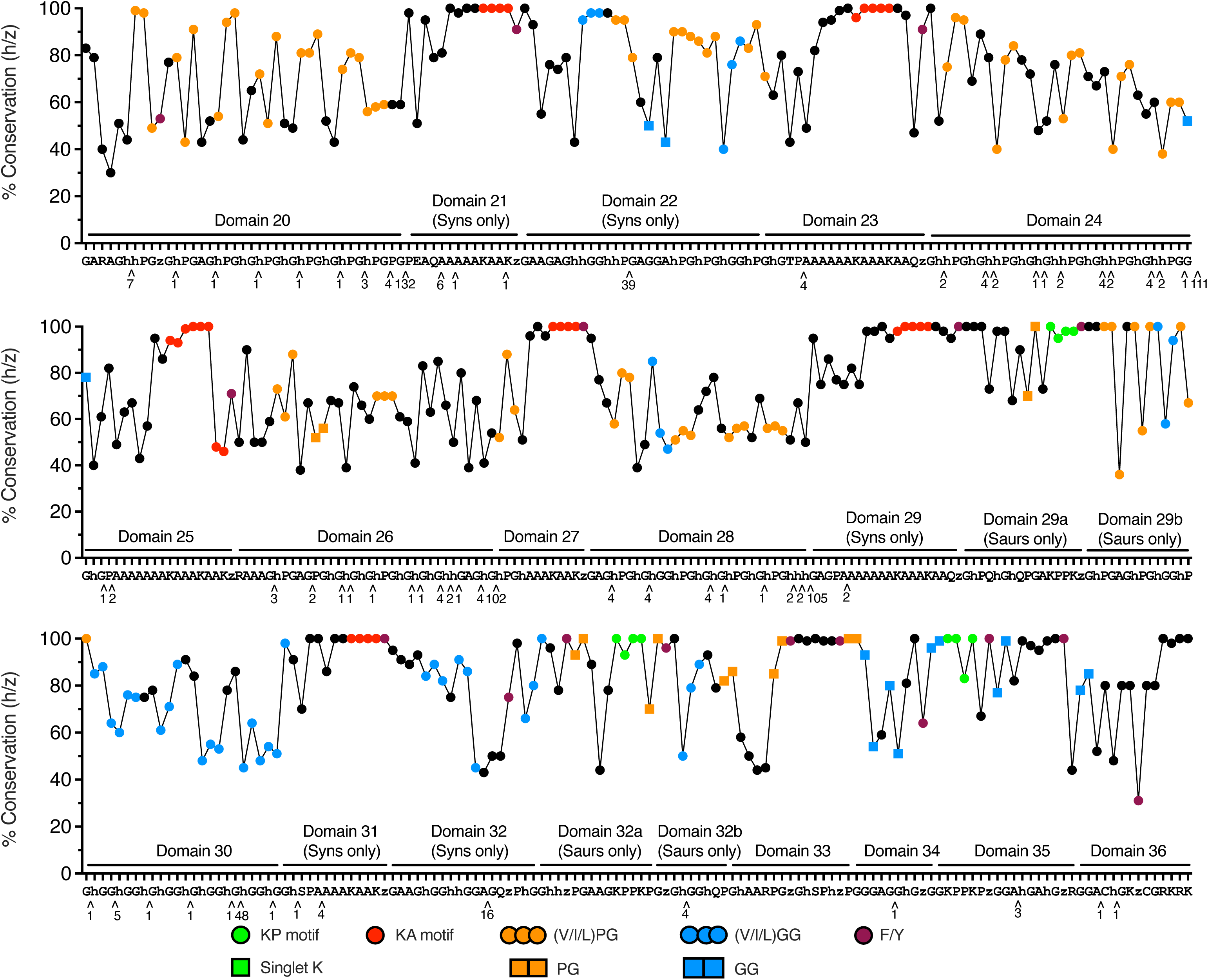
Consensus sequence of Amniote tropoelastins, with residue-by-residue quantification of sequence conservation. **A:** Domains 2 to 19. **B:** Domains 20 to 36. Approximately 80 sequences of Amniote tropoelastins were used to generate this consensus sequence, including approximately equal numbers of Synapsid and Sauropsid species. Methodology for generation of the consensus sequence is described in the text. Details of the calculation of the % Coacervation (h/v) parameter, the proportional frequency of the consensus amino acid at a given sequence position, are shown in Table 1. Consensus gaps, which arise when the majority of aligned sequences contained a gap at that sequence position, were not included in the calculation and are not plotted. However, the numbers of gaps and their locations are indicated below the consensus sequence. Common motifs and other sequence features were color-coded in the plot. These included KP- and KA-type cross-linking motifs, as well as positions of singlet lysine residues, not including lysine residues in domain 36. hPG and hGG motifs (where h is V, I or L) as well as PG and GG pairs and positions of F/Y amino acid residues are also indicated. Domain locations in the consensus sequence are indicated.

The data clearly demonstrate the presence of many areas with very strong positional sequence conservation across all Amniote tropoelastins. As might be expected, both KA- and KP-type cross-linking domains have a very high degree of sequence conservation. Similarly, consistent with previous observations on a smaller data set^2^, domains 8-14 contain the greatest degree of almost 100% positional sequence conservation across all Amniote tropoelastins. Regions of high sequence conservation across species are also evident in domains 2 and 3 and domains 30 to 36. As noted earlier, these areas may represent sites of interaction with other proteins important for the assembly of the elastin polymer or influencing the final architecture of the elastin matrix.

Also evident from the color-coding of the consensus sequence is the differing frequency of occurrence of PG and GG pairs in hydrophobic domains in the replicated region (domains 16- 28) as compared to hydrophobic domains in the flanking regions (domains 3, 5, 7 and 30 to 35), as previously documented in Figure 5.

Two parameters were used to assess levels of positional consensus averaged across domains. Again, examples for their calculation are shown in Table 1. The first of these, designated ‘% Conservation (≥95%)”, was calculated as the number of sequence positions within a domain that had a conservation greater or equal to 95%, divided by the total number of sequence positions in the domain. The level of ≥95% (rather than 100%) was chosen to allow for a small number of raw sequence data misreads at that position over the more than 80 species used.

The second parameter, designated ‘% Conservation (domain matrix)’, was used as a measure of overall domain sequence conservation, and is calculated from the matrix of sequence positions vs. specific amino acids. To derive this parameter, the sum of sites in the domain matrix that were in agreement with the consensus amino acid at each particular sequence position was divided by the total number of sites in the domain matrix. See Table 1 for example calculations.

Plots of averaged domain and region conservation are shown in Figure 7. In both cases, h/z sequence data was plotted, and consensus gaps were excluded from the calculations, as described above. Data from Synapsid and Sauropsid consensus alignments were plotted separately, recognizing differences with respect to the occurrence of some exons/domains between the genomic sequences of these major sub-groups of Amniotes (see Figure 2, and section 7.1).

**Figure 7.**
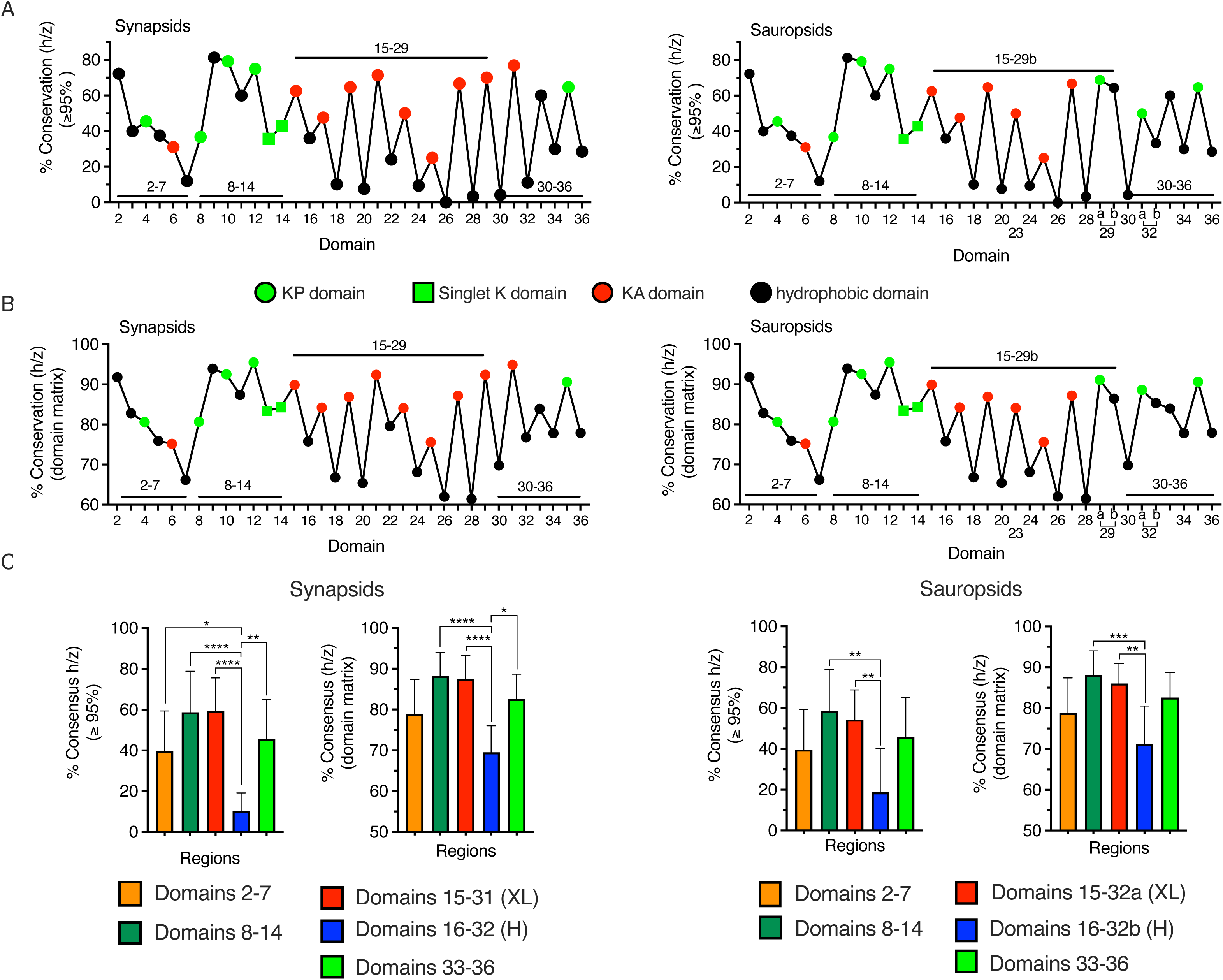
Domain-by-domain and regional conservation across Amniote species. Synapsid and Sauropsid consensus sequences were plotted separately, reflecting the absence of exons/domains 21, 22, 29, 31, 32 from Sauropsids, and their replacement with domains 29a, 29b, 32a and 32b. As in Figure 6, the h/z consensus sequence was used, and consensus gaps were not included in the calculations. **A:** The % Conservation (h/z) (≥95%) parameter represents the proportion of sequence positions within a domain that had a conservation greater or equal to 95% (see text and Table 2 for details). Cross-linking domains and hydrophobic domains are color-coded on the plots. **B:** The % Conservation (h/z) (domain matrix), an alternate method to assess overall domain conservation, is calculated as the proportion of the sites in the overall sequence position vs. amino acid matrix that were in agreement with their respective consensus amino acid (see text and Table 2 for details). Cross-linking domains and hydrophobic domains are color-coded on the plots. **C:** Domain conservation parameters from Panels A and B averaged across the specified regions of the tropoelastin consensus sequence. for the domain 15-32b region, cross-linking domains (XL) are averaged separately from hydrophobic domains (H). Statistical comparisons used ANOVA followed by a Tukey correction for multiple comparisons. P-values are as follows: ****, P<0.0001; ***, P<0.001; **, P<0.01; *, P<0.05.

**Table 2.**
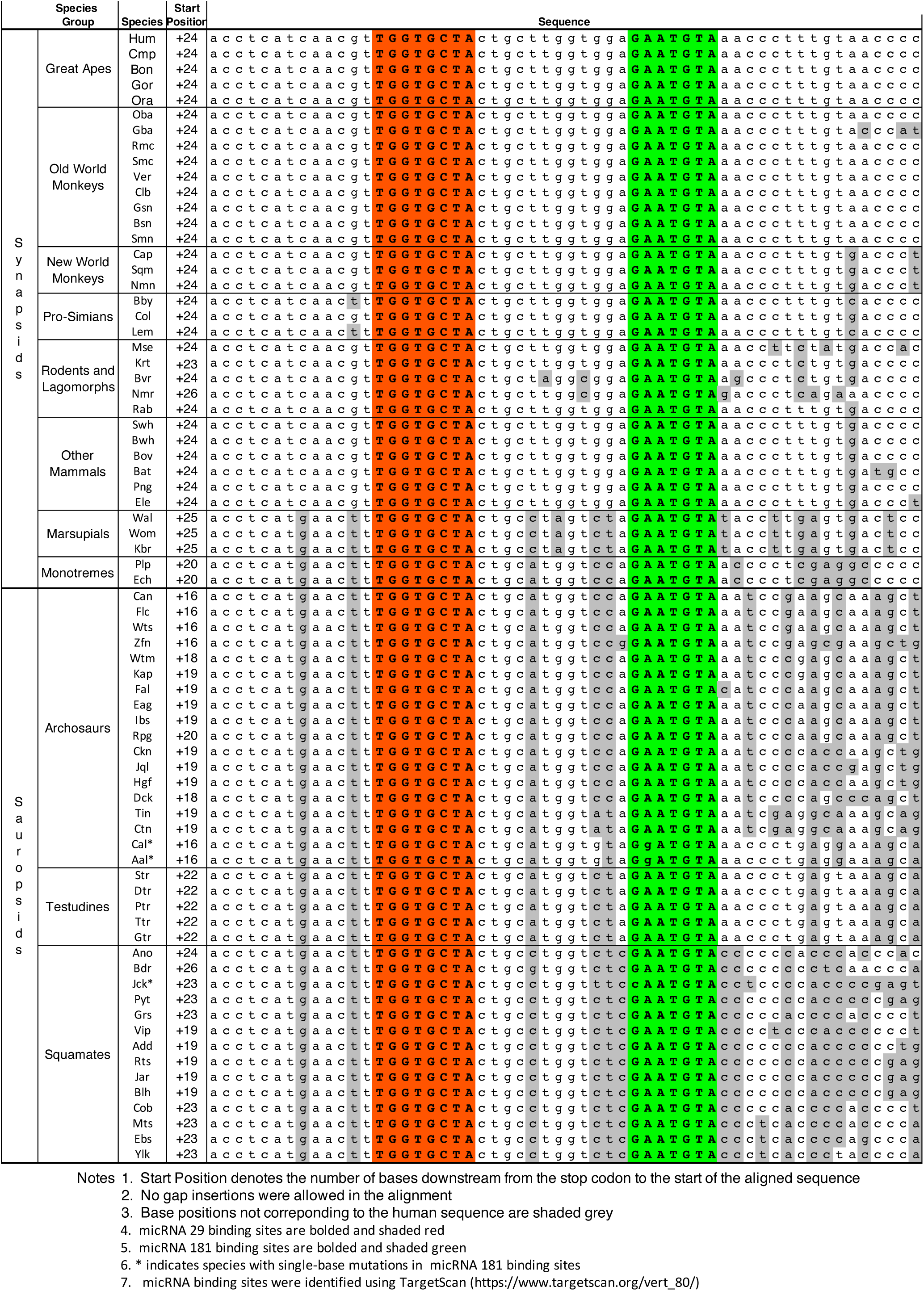
Conservation of Proximal 3’utr Sequence in Amniote Tropoelastins.

Whether plotted as % Conservation (h/z) (≥ 95%) (Figure 7A) or % Conservation (h/z) (domain matrix) (Figure 7B), the patterns of averaged domain conservation were essentially identical for Synapsids and Sauropsids, in spite of their evolutionary separation of approximately 300 million years and the genomic differences in occurrence of some exons/domains between these species groups. In all cases, the results emphasize the consistently strong positional sequence conservation of domains 8-14, the generally high positional sequence conservation of both KA and KP cross-linking domains, and the generally low positional sequence conservation of hydrophobic domains, particularly in the region of the sequence including domains 16-32.

Domain conservation data averaged across regions of the Synapsid and Sauropsid consensus sequences are plotted in Figure 7C. Note that, for the domain 15-32b region, cross-linking domains (XL, i.e. domains 15, 17, 19, 21, 23, 25, 27, 29, 29a, 31, 32a) are averaged separately from hydrophobic domains (H, i.e. domains 16, 18, 20, 22, 24, 26, 28, 29b, 30, 32b). The data for both Synapsid and Sauropsid groups emphasize the consistently low conservation of positional sequence of hydrophobic domains of the ‘replicated region’. Although generally lacking positional sequence conservation, a subsequent analysis of these hydrophobic domains showed a notable conservation of ‘style’ or character of sequence (see section 6).

Parameters used to quantify positional sequence conservation may, in fact, underestimate such conservation. For example, while data from particular domains that were absent from genomic sequence of a species or lost due to sequencing gaps were not used in the calculations, the deletion of the otherwise well-conserved central portion of domain 36 in all squamates would negatively influence the averaged consensus calculations for those positions in Sauropsids. In other cases, the incidence of amino acids at a given sequence position might be almost evenly divided between two closely related amino acids (e.g. serine and threonine), resulting in a misleadingly low % Conservation of approximately 50%. However, for the sake of simplicity such cases were not taken into account in the calculations.

### 5.3 Validating the representative nature of the consensus sequences

As discussed earlier, sequences for tropoelastins have an unusual character, with shared low-complexity amino acid distributions together with an apparent evolutionary history of domain replications followed by abundant modifications within domains involving further motif replications, deletions and insertions. The nature of these sequences therefore necessitated an unusual approach to establishing plausible consensus sequences for Amniotes species in general, and for Synapsid and Sauropsid sub-groups of species.

In order to assess how well the consensus sequences represented the contributing tropoelastin sequences, the collective or non-positional properties of the consensus sequences determined for Synapsids and Sauropsid sub-groups were compared to these collective properties averaged over these species groups. (Supplementary Figure S6). Parameters compared included patterns of amino acid composition and other general characteristics such as isoelectric pH and hydropathicity (Supp Figure S6A), the proportion of prolines present in PG pairs and the overall ratio of prolines to glycines (Supp Figure S6B), and the varied distribution of densities of GG/PG sequence pairs across regions of the sequences (Supp Figure S6C). For all such parameters, the characteristics of the consensus sequences were entirely consistent with the averaged values for Synapsid and Sauropsid species sub-groups (Figures 3, 4 and 5), confirming that the derived consensus sequences were indeed representative of the collective characteristics of the contributing tropoelastin sequences.

### 5.4 Conservation of sequences of the signal peptides (domain 1) of Amniote tropoelastins

As for most proteins destined for transport into the extracellular matrix, all tropoelastins included a signal peptide sequence, present entirely in domain 1 of the protein. An alignment of domain 1 representing more than 80 species of Amniote tropoelastins is shown in Supplementary Table S4

Signal peptide sequences have been characterized as having 4 general regions. These consist of an N-region, usually beginning with a methionine residue and containing proximal positively charged amino acids, followed by an alpha-helical H-region particularly rich in leucine residues, followed by beta-sheet C/Pro-regions, providing a recognition sequence for the membrane-bound signal peptidase responsible for cleavage of the signal peptide from the mature protein once it has been directed into the endoplasmic reticulum of the cell. See Owji et al.^66^ for a detailed general review of sequence and structural characteristics of signal peptides.

Domain 1 of all tropoelastins conform to this general structure. However, for most Synapsid species the N-region is not particularly rich in positively charged amino acids. Indeed, Synapsids have a consistent N-terminal leading sequence of MAGL(T/S)A, with the first and only R or K residue approximately 10 residues downstream of the initiator methionine. This lead sequence transitions to MA(S/T)RTA in non-placental mammals (marsupials and monotremes), to MARQ(A/L/P) in archosaurs and testudines and then to MARLR in squamates.

Signal peptides of all tropoelastins include a central, leucine-rich H-domain, the general length and character of which is well-conserved between Synapsid and Sauropsid species. The exception to this is tropoelastin from naked mole rat, which has an extended H-region. Unusually, this extension includes a proline residue, interrupting the normal alpha-helical structure of the region. Such helix-breaker residues inserted into the H-region have been suggested to favor certain secretory pathways^67^. On the other hand, the presence of this unusual proline residue might simply be attributed to a single-base misread of P for L in the raw sequence data.

The final C-region/Pro-region leading to the cleavage site for tropoelastins is also reasonably consistent across species. In primates, this sequence is conserved as PSRP-GGVG. The predicted cleavage site (using SignalP-6.0, https://biolib.com/DTU/SignalP-6/) is indicated by a dash. The sequence of the region transitions to PSQG-GGVG in most other mammals, to TRQG-GGVG in marsupials, (T/S)QQG-GGVG in most archosaurs, and S(W/R)QG-GGVG in testudines and squamates. As noted previously, in all cases the signal peptide is entirely contained within domain 1, and the final amino acid of domain 1 (glycine), becomes the first amino acid of the sequence of mature tropoelastin. All mature tropoelastins begin with this glycine residue.

Whether these particular characteristics of signal peptides in Amniote tropoelastins have any role to play in determining the rate or efficiency of passage of the protein into the endoplasmic reticulum, direct the protein into specific secretory pathways, or simply represent permissible mutation events over evolution is not clear from these data. However, it may be of significance to note that the highly non-polar nature of tropoelastins renders the protein susceptible to aggregation. There have been several reports of association of tropoelastin with ‘companion’ proteins in the secretion pathway that both prevent premature, intra-cellular oligomerization and may also be important for regulating polymeric assembly in the extracellular matrix^67–70^.

### 5.5 Conservation of sequence motifs in the 3’utr of Amniote tropoelastins

Conservation of proximal 3’utr sequence of tropoelastins had previously been identified in a much more limited subset of species^2^. Indeed, this region provided a reliable and specific search sequence for locating tropoelastins in genomic sequence databases. Table 2 shows an alignment of 56 bases downstream but proximal to the stop codon for 36 Synapsid and 37 Sauropsid species. Although the start point of the aligned sequence region with respect to the stop codon has been shifted by a few bases between species (from 16-25 base positions), no gaps were introduced in the alignment downstream of that start point. Motifs shaded red and bolded are complete binding motifs for the micRNA29 family. Motifs shaded in green and bolded are complete binding motifs for micRNA181. Bases that differ from the base at that position in the human sequence are shaded grey. All unshaded bases are identical to the base in the human sequence at that position.

Tropoelastin 3’utrs contain multiple polyadenylation signal motifs (polyA, e.g aataaa), with the second of these thought to be used for termination of transcription, at least in the case of human tropoelastin. The region of sequences aligned in Table 2 is, in all cases, upstream of the first polyA motif in the 3’utr.

One or more additional copies of complete or partial micRNA29 (but not micRN181) target sites are also found downstream of the sequences shown in Table 2, as well as a single target site for micRNA101. MicRNA101 target sites are consistently present in all Synapsids (placental and non-placental mammals), but only present in some Sauroposid sub-groups (testudines and non-avian archosaurs). In most cases these additional micRNA sites are also upstream of the first polyA motif. Archosaur sequences are exceptions to this arrangement. For these species the first polyA motif follows the initial micRNA29/micRNA181 pair, and additional micRNA29 binding sites are located between the first and second polyA motifs. In general, the first polyA motif appears much earlier in Archosaurs compared to all other species, i.e. ∼200 bases rather than ∼1000 bases downstream of the stop codon. See Supplementary Table S5 for detailed information on the relative arrangements of micRNA target sequences with respect to polyadenylation sites in Amniote tropoelastins. Such microRNA target sequences, located between the translation stop codon and functional polyadenylation sites and consistently conserved over such evolutionary significant periods of time, strongly suggests a role in regulation of tropoelastin expression.

The stability and lack of turnover of elastin polymers, once laid down in many if not most elastic tissues, implies that, after a relatively brief developmentary period of rapid production, further synthesis and accumulation of polymeric elastin must cease. Although mechanisms for regulation of production and assembly of tropoelastin are not well-understood, control is likely exerted at several levels, including transcription, mRNA stability and translational efficiency. Indeed, there have been suggestions that destabilization of tropoelastin mRNA, perhaps related to elements of secondary structure in the 3’utr, may be one important mechanism for post-transcriptional down-regulation of tropoelastin synthesis^71–76^. The micRNA29 family of non-coding RNAs has been generally linked to the regulation of several extracellular matrix proteins, and up-regulation of this micRNA has been associated with suppression of expression of tropoelastin and other ECM genes^77–80^. In contrast micRNA181 has primarily been associated with cell cycle regulation in inflammation, immunity and cancer^81–84^. Similarly, micRNA101 has primarily been associated with regulation of cell proliferation and fibrosis^85^.

With this in mind, it is of particular interest that modeling of the structure of the first 80 bases of the 3’utr of all Amniote tropoelastins (RNAstructure, version 6.4, https://rna.urmc.rochester.edu/RNAstructureWeb/Servers/Predict1/Predict1.html), predicts a stable stem-loop structure in this region, with the micRNA29 and micRNA181 target sequences forming the two sides of the stem (Supplementary Figure S7). Upregulation of either or both of these non-coding RNAs would be expected to destabilize this structure, with potential consequences for the stability of the mRNA and the expression of tropoelastin.

## 6. Regions of low positional sequence conservation in Amniote tropoelastins

### 6.1 Investigation of the sequence characteristics of highly non-polar domains

Non-polar domains of tropoelastin sequences generally showed very low levels of positional sequence conservation with respect to the derived Amniote consensus sequence (Figures 6 and 7). This was especially the case for the hydrophobic domains of the replicated region (domains 16, 18, 20, 22, 24, 26, and 28), but was also true for several hydrophobic domains in the GG-rich flanking regions either preceding the central conserved region or downstream of the replicated region (domains 3, 5, 7, 30, 32).

In spite of their low scores for positional sequence conservation, all of these domains maintained a pattern of sequence characterized by PG, GG and VG pairs, abundant but variable in number and present in motifs as tandem or partial repeats. Attempts to align these motifs to highlight overall positional sequence conservation (see section 5) suggested a possible underlying sequence repeat pattern of [hPGhGG]_n_, where h represents non-polar amino acids (V, I, L, A, G, F, Y, Q). Replication of this sequence pattern, followed by extensive and variable mutations, especially insertions and deletions, might account for the characteristic ‘style’ of these sequences.

To investigate further, H-domain sequences, defined as domains 3, 5, 7, 16, 18, 20, 22, 24, 26, 28, 30 and 32) were searched for all occurrences of xPG, PGx, xVG, VGx, xGG, GGx, GxG and GxP motifs, where ‘x’ included all amino acids with the exception of histidine, tryptophan and methionine, for a total of 136 possible motif types. To avoid duplicate counting, PGG was counted as PGx, VGG and GGG were counted as xGG, GGP was counted as GGx, GVG was counted as GxG, and GPG was counted as xPG.

A total of 20,991 such ‘x-motifs’ were identified in H-domains of collected Amniote sequences, distributed approximately equally between Synapsid and Sauropsid sub-groups. H-domain sequences were also sub-divided into hydrophobic domains from the ‘replicated region’ of tropoelastins (designated domains 16H-28H) and those from flanking hydrophobic domains (domains 3, 5, 7, 30, 32). In all groups, >90% of the x-motifs could be classified as h-motifs, where x was restricted to V, I, L, A. G. F, Y, or Q (Table 3).

**Table 3.** Motif Analysis in Hydrophobic (H)-Domains) with Low Positional Sequence Conservation.

|  | All Amniotes |  |  | Synapsids |  |  | Sauropsids |  |  |
| --- | --- | --- | --- | --- | --- | --- | --- | --- | --- |
|  | Total x<br>Motifs | Total h<br>Motifs | %<br>h Motifs | Total x<br>Motifs | Total h<br>Motifs | %<br>'h' Motifs | Total x<br>Motifs | Total h<br>Motifs | %<br>'h' Motifs |
| All H-Domains | 20991 | 20270 | 96.6 | 10949 | 10744 | 98.1 | 10558 | 9632 | 91.2 |
| Domains 3,5,7,30,32 | 5601 | 5435 | 97.0 | 3090 | 3060 | 99.0 | 2812 | 2576 | 91.6 |
| Domains 16H-28H | 15,390 | 14835 | 96.4 | 7859 | 7684 | 97.8 | 7746 | 7056 | 91.1 |

Motifs include xPG, PGx, xVG, VGx, xGG, GGx, GxG and GxP
x = V, I, L, A, G, F, Y, Q, K, R, N, S, T, E, D, C, P
h = V, I, L, A, G, F, Y, Q
Domains 16H-28H include domains 16, 18, 20, 22, 24, 26 and 28

Motifs were ranked according to frequency of occurrence in H-domains of collected sequence sets of Amniotes, Synapsids or Sauropsids (Table 4). For each group, the top ≥95% of these ranked motifs were analyzed for motif type. These most frequent motifs, consisting almost entirely of h-motifs (hPG, PGh, GhG, hGG, GGh and GhP), were individually counted and color-coded (Table 5). Less frequent but recurrent motifs were collectively designated as ‘other motifs’ (e.g. VGV, VGA, GTG, GGT, PGT, VGI, VGP). Most of these ‘other motifs’ could plausibly arise through single-base mutations in the codons of the color-coded motifs.

**Table 4.**
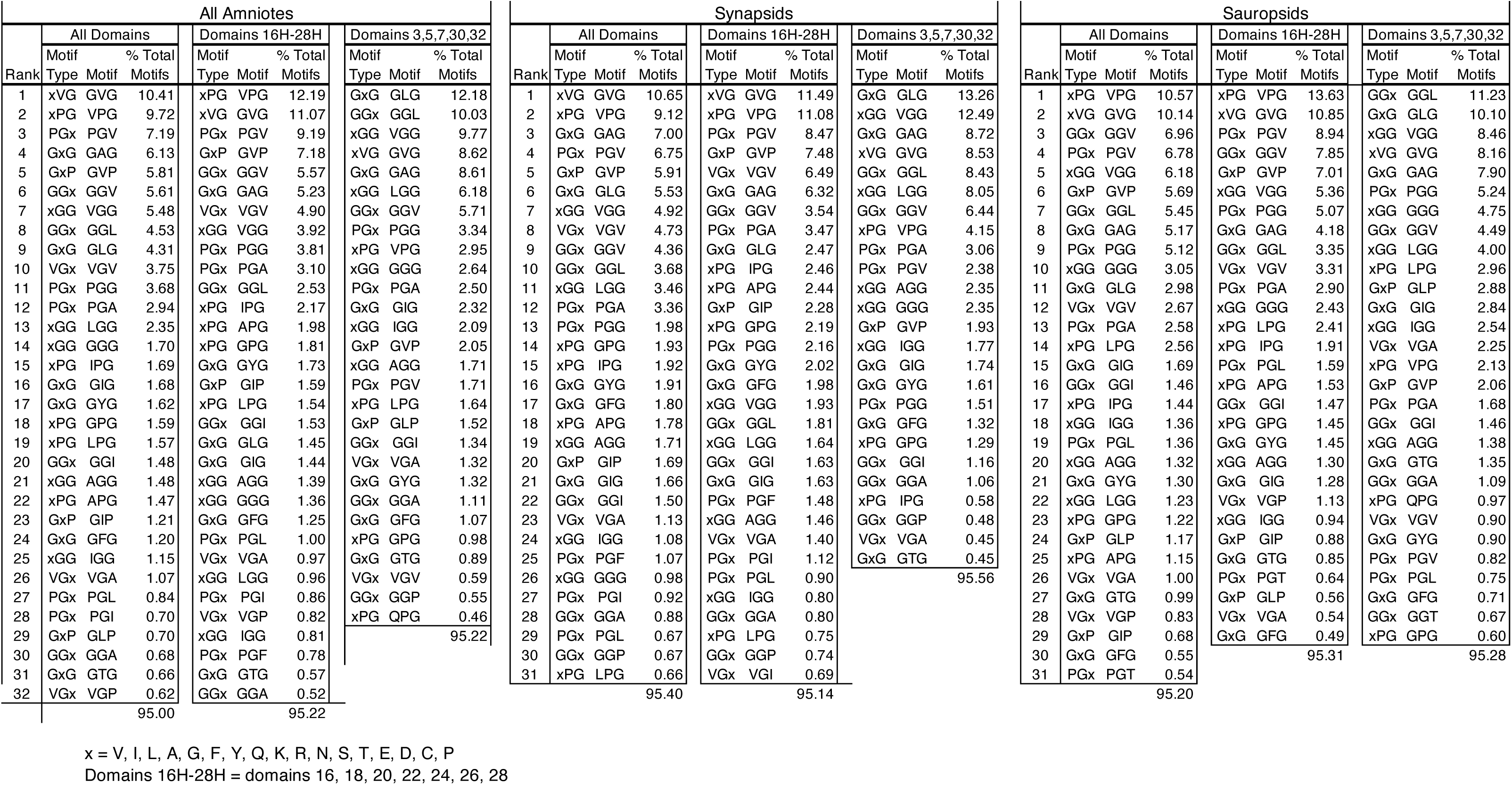
Motif Frequency Ranking (top 95% of motifs)

**Table 5.** Motif Frequencies in H-Domains of Amniotes Accounting for ≥95% of All Motifs.

| A | All Amniotes (% Frequency, ≥95% of 20,991 total motifs) |  |  |  |  |  |  |  |  |  |  |  |  |
| --- | --- | --- | --- | --- | --- | --- | --- | --- | --- | --- | --- | --- | --- |
|  | h-Motifs |  |  |  |  |  | Other Motifs |  |  |  |  |  |  |
|  | hPG | PGh | GhG | hGG | GGh | GhP | VGV | VGA | GTG | GGT | PGT | VGI | VGP |
| All H-Domains | 16.04 | 15.35 | 25.35 | 12.17 | 12.30 | 7.72 | 3.75 | 1.07 | 0.66 |  |  |  | 0.62 |
|  | Total h-Motifs |  |  |  |  | 88.9 |  |  |  |  |  |  |  |
| Total h-Motifs (inc. single-base mutations) |  |  |  |  |  | 94.4 |  |  |  |  |  |  |  |
| Domains<br>16,18,20,22,24,26,28 | 19.69 | 18.73 | 22.16 | 8.45 | 10.15 | 8.77 | 4.90 | 0.97 |  | 0.57 |  |  | 0.82 |
|  | Total h-Motifs |  |  |  |  | 88.0 |  |  |  |  |  |  |  |
| Total h-Motifs (inc. single-base mutations) |  |  |  |  |  | 94.4 |  |  |  |  |  |  |  |
| Domains 3,5,7,30,32 | 6.41 | 7.95 | 34.12 | 22.39 | 18.19 | 4.12 | 0.59 | 1.32 | 0.89 |  |  |  |  |
|  | Total h-Motifs |  |  |  |  | 93.2 |  |  |  |  |  |  |  |
| Total h-Motifs (inc. single-base mutations) |  |  |  |  |  | 96.0 |  |  |  |  |  |  |  |

| B | Synapsids (% Frequency, ≥95% of 10,744 total motifs) |  |  |  |  |  |  |  |  |  |  |  |  |
| --- | --- | --- | --- | --- | --- | --- | --- | --- | --- | --- | --- | --- | --- |
|  | h-Motifs |  |  |  |  |  | Other Motifs |  |  |  |  |  |  |
|  | hPG | PGh | GhG | hGG | GGh | GhP | VGV | VGA | GTG | GGT | PGT | VGI | VGP |
| All H-Domains | 15.41 | 14.74 | 28.54 | 12.15 | 10.41 | 8.26 | 4.73 | 1.13 |  |  |  |  |  |
|  | Total h-Motifs |  |  |  |  | 89.5 |  |  |  |  |  |  |  |
| Total h-Motifs (inc. single-base mutations) |  |  |  |  |  | 95.4 |  |  |  |  |  |  |  |
| Domains<br>16,18,20,22,24,26,28 | 18.92 | 17.61 | 25.92 | 5.84 | 7.77 | 10.50 | 6.49 | 1.40 |  |  |  | 0.69 |  |
|  | Total h-Motifs |  |  |  |  | 86.6 |  |  |  |  |  |  |  |
| Total h-Motifs (inc. single-base mutations) |  |  |  |  |  | 94.5 |  |  |  |  |  |  |  |
| Domains 3,5,7,30,32 | 6.02 | 6.95 | 35.18 | 27.00 | 17.57 | 1.93 |  | 0.45 | 0.45 |  |  |  |  |
|  | Total h-Motifs |  |  |  |  | 94.7 |  |  |  |  |  |  |  |
| Total h-Motifs (inc. single-base mutations) |  |  |  |  |  | 95.6 |  |  |  |  |  |  |  |

| C | Sauropsids( % Frequency, ≥95% of 10,558 total motifs) |  |  |  |  |  |  |  |  |  |  |  |  |
| --- | --- | --- | --- | --- | --- | --- | --- | --- | --- | --- | --- | --- | --- |
|  | h-Motifs |  |  |  |  |  | Other Motifs |  |  |  |  |  |  |
|  | hPG | PGh | GhG | hGG | GGh | GhP | VGV | VGA | GTG | GGT | PGT | VGI | VGP |
| All Domains | 16.96 | 15.84 | 21.82 | 13.15 | 13.87 | 7.55 | 2.67 | 1.00 | 0.99 |  | 0.54 |  | 0.83 |
|  | Total h-Motifs |  |  |  |  | 89.2 |  |  |  |  |  |  |  |
| Total h-Motifs (inc. single-base mutations) |  |  |  |  |  | 94.4 |  |  |  |  |  |  |  |
| Domains<br>16,18,20,22,24,26,28 | 20.94 | 18.50 | 18.24 | 10.02 | 12.67 | 8.45 | 3.31 | 0.54 | 0.85 |  | 0.64 |  | 1.13 |
|  | Total h-Motifs |  |  |  |  | 88.8 |  |  |  |  |  |  |  |
| Total h-Motifs (inc. single-base mutations) |  |  |  |  |  | 94.2 |  |  |  |  |  |  |  |
| Domains 3,5,7,30,32 | 6.66 | 8.50 | 30.61 | 21.15 | 18.26 | 4.94 | 0.90 | 2.25 | 1.35 | 0.67 |  |  |  |
|  | Total h-Motifs |  |  |  |  | 90.1 |  |  |  |  |  |  |  |
| Total h-Motifs (inc. single-base mutations) |  |  |  |  |  | 95.3 |  |  |  |  |  |  |  |

'h' in h-motifs refers to amino acids V, I, L, A, G, F, Y, or Q
H-Domains include domains 3,5,7,16,18,20,22,24,26,28,30 and 32
VGV, VGA, GTG, GGT, PGT motifs (grey, hatched) may be generated by a single-base mutation in an h-moti
VGV, VGA, VGI motifs may also be generated by a single amino acid deletion from an hPGhGG repeat
VGP motifs may be generated by a double amino acid deletion from an hPGhGG repeat

Color-coded motifs were mapped onto the putative sequence repeat pattern of [hPGhGG]_n_ (Figure 8A). These motifs provided complete coverage of this sequence pattern, including regions of repeat overlap, and together accounted for almost 90% of all x-motifs. The results were consistent with the postulation that the character of these H-domain sequences was derived from this underlying sequence repeat pattern, subsequently modified by numerous mutations, insertions and deletions, but preserving a recognizable ‘style’ of sequence. This pattern was seen not only for sequences generated from all Amniotes, but also for sequences of Synapsids and Sauropsid sub-groups, indicating that, in spite of low levels of positional sequence conservation, this sequence ‘style’ was retained over an evolutionary separation of 300 myrs.

**Figure 8.**
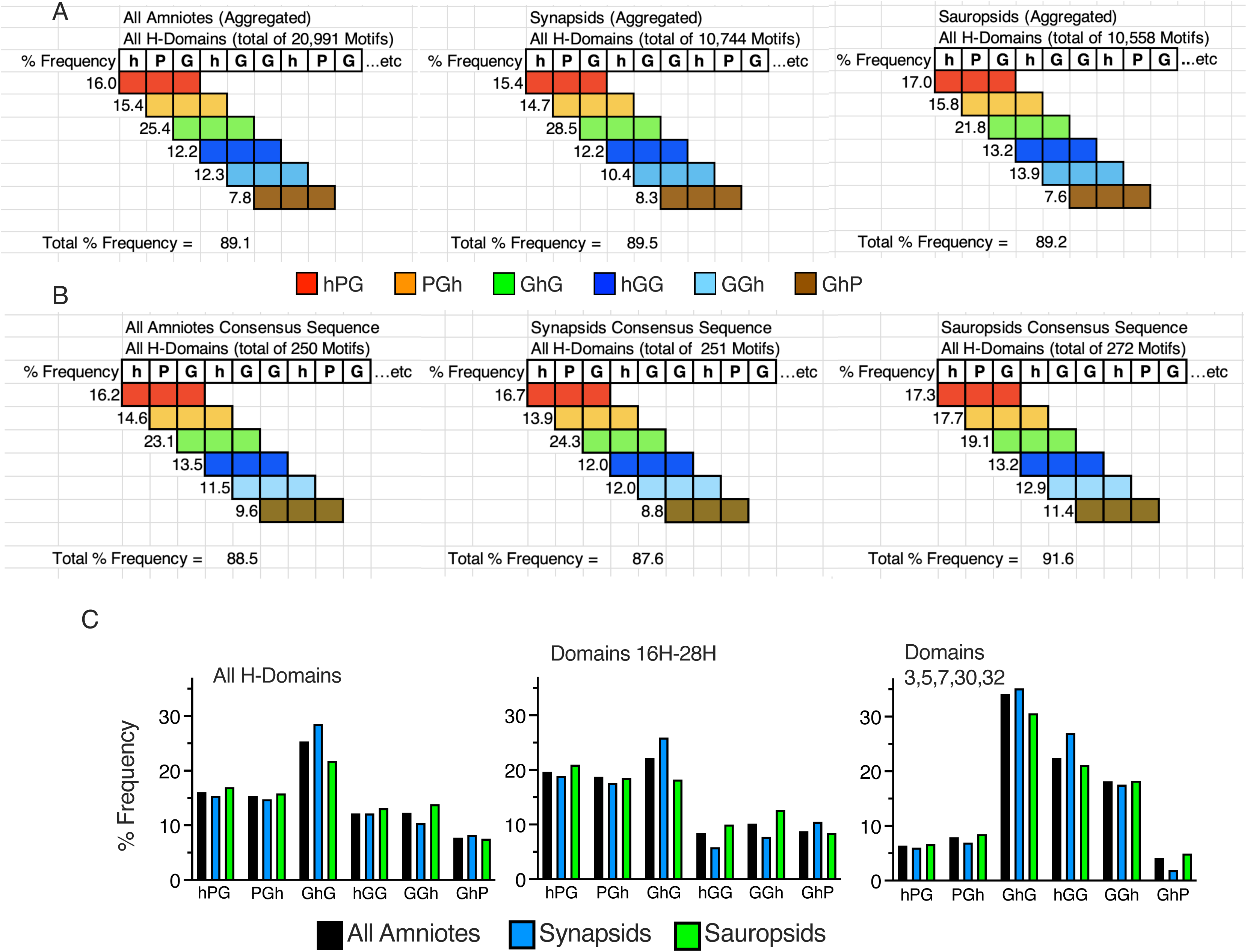
Analysis of motif types and motif frequencies in hydrophobic domains (H-domains) of Amniotes. H-domains include domains 3, 5, 7, 16, 18, 20, 22, 24, 26, 28, 30 and 32, and can be sub-divided into domains that are present in the ‘replicated region’ of tropoelastin (domains 16, 18, 20, 22, 24, 26, 28, designated ‘Domains 16H-28H’) and domains that flank the ‘replicated region’ on either side (designated ‘Domains 3, 5, 7, 30, 32’). Motifs counted included xPG, PGx, xVG, VGx, xGG, GGx, GxG and GxP, where x is any amino acid except for histidine, tryptophan or methionine, for a total of 129 possible motif types. ‘h’-motifs are a sub-class of x- motifs, where x is V, I, L, A, G, F, Y or Q. Motifs were ranked according to frequency, and motifs accounting for the top ≥95% of all motifs were analysed. **A:** Frequencies of motif types present in aggregated sequences of all H-domains of all Amniotes, and of Synapsid and Sauropsid sub-groupings are mapped against a putative fundamental tandemly repeated sequence of hPGhGG. Total numbers of motifs are indicated for each group. h-motif types are color-coded. The frequency of any given h-motif is indicated beside the motif, with the sum of these frequencies appearing below the map. For all Amniotes, as well as for Synapsid and Sauropsid subgroups, almost 90% of all motifs map directly onto the hPGhGG repeat sequence, i.e. hPG, PGh, GhG, hGG, GGh and GhP. **B:** Similar mapping of frequencies of motif types present in the H-Domains of the consensus sequences (Figure 6). The distribution of motif types in the consensus sequence is essentially identical to that in the aggregated H-Domain data in panel A. **C:** Frequency distributions of h-motif types, where h = V, I, L, A, G, F, Y and Q. Frequencies are plotted for all Amniotes, and for Synapsid and Sauropsid sub-groupings. Separate plots are provided for all H-domain sequences and for sub-groupings of domains 16H-28H and domains 3,5,7,30,32.

A similar motif frequency analysis done on H-domain consensus sequence data for all Amniotes as well as for Synapsid and Sauropsid sub-groups showed motif frequencies very similar to those seen in the aggregated data (Figure 8B), providing further confirmation that these derived consensus sequences were representative of the aggregated sequence data, even with respect to these domains with low positional sequence conservation.

We had demonstrated above (Figure 5) that the relative densities of PG and GG sequence pairs varied between the domains making up the ‘replicated region’ (domains 15-29b), and the hGG-rich flanking regions (domains 2-7 and 30-35) of tropoelastins. This was also evident in the residue-by-residue consensus sequence data (Figures 6 and 7). We therefore compared the distribution of h-motifs in domains 16H-28H to that in domains 3, 5, 7, 30, 32 for all Amniotes as well as between Synapsid and Sauropsid sub-groups (Figure 8C). For each sub-set of H-domains the % frequency of each major motif type was close to that for all Amniotes, and consistent between Synapsids and Sauropsids sub-groups. However, patterns of motif frequency were clearly different between the two sub-sets of H-domains. Differences included relatively increased frequencies of hGG and GGh motifs and correspondingly lower frequencies of hPG and PGh motifs in the domain 3, 5, 7, 30, 32 sub-set. These data were consistent with our earlier observations in Figures 6 and 7, and suggested that, if they are indeed derived from the same fundamental repeat pattern of [hPGhGG]_n_, the subsequent evolutionary process of mutation, insertion and deletion applied to this repeated pattern was consistently different between these sub-sets of H-domains.

Finally, for each of these major motifs we determined the frequency of appearance of individual amino acids in the x-position of a given motif. Data shown in Figure 9 are for all H-domain sequences, and compare amino acid frequencies between Synapsid and Sauropsid sub-groups. Amino acids of the h-group (V, I, L, A, G, F, Y, Q) are individually color-coded and plotted. Other amino acids (K, R, N, S, T, E, D, C. P) are collectively shown in black. Below each color wheel is the total number of motif types counted. Above the wheel is the proportion of the motifs that contained an h-group amino acid. This proportion was generally greater than 90%.

**Figure 9.**
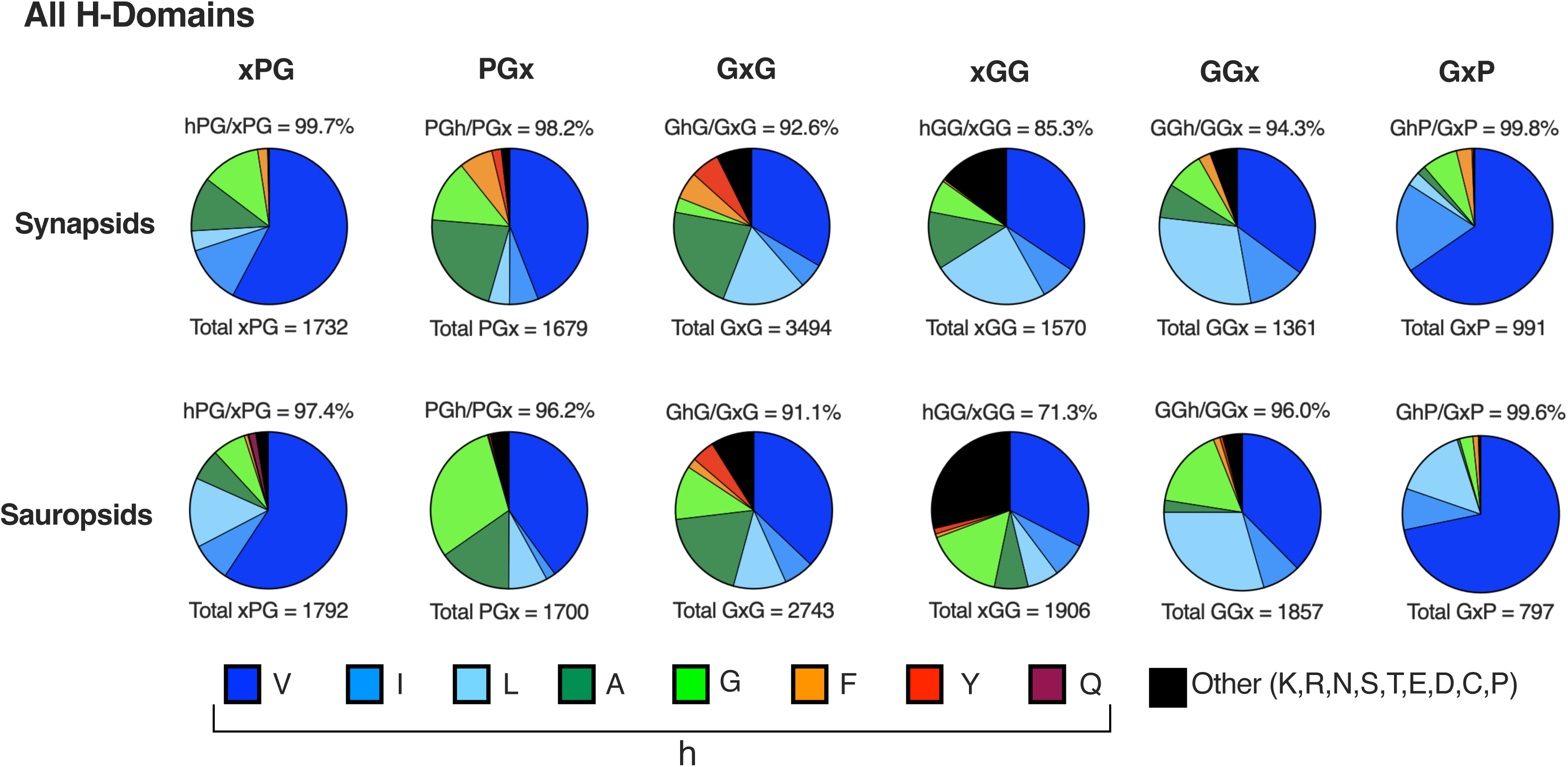
Frequency distributions of amino acids in the x position of xPG, PGx, GxG, xGG, GGx and GxP motifs. ‘h’-amino acids (V, I, L, A, G, F, Y or Q) are individually color-coded as indicated. Other amino acids in the x-position (K, R, N, S, T, E, D, C or P) are collectively designated as ‘other’ and color-coded as black. The proportion of h-amino acids compared to all amino acids present in each motif type is shown above the color wheel. The total count of each motif type is shown below the color wheel. Frequency distributions are shown for all H-domains, with Synapsid and Sauropsid data compared.

Clearly the evolutionary selection of amino acids present in these x-motifs is not random. Valine dominates the selection, and valine, isoleucine and leucine together make up at least 50% of all amino acids appearing in any x-motif. Aromatic and other amino acids occur only very infrequently. In the case of GxG and xGG motifs, which included more substantial proportions of ‘other’ amino acids, the major contributor to the ‘other’ group was P, as in GPG or PGG. Furthermore, the general selection of these amino acids is remarkably consistent between Synapsid and Sauropsid sequence groups. Possible reasons for this strong bias towards V, I, and L in the evolutionary selection of these amino acids will be discussed below.

### 6.2 Collective conservation of total h-motifs and h-motif distributions in H-Domains of Amniote tropoelastins

In spite of substantial variability in size of H-Domains across individual species, measured as the number of amino acids (Figure 10A), the collective sum of h-motifs over all H-Domains was strongly conserved between Synapsid and Sauropsid tropoelastins (Figure 10 B). Similarly conserved was the ratio of total h-motifs to total x-motifs (approximately 90%) in these hydrophobic domains (Figure 10C). Furthermore, patterns of distribution of individual h-motif types between Synapsid and Sauropsid species groups were remarkably consistent (Figure 10D), in spite of the prolonged evolutionary separation of these species groups.

**Figure 10.**
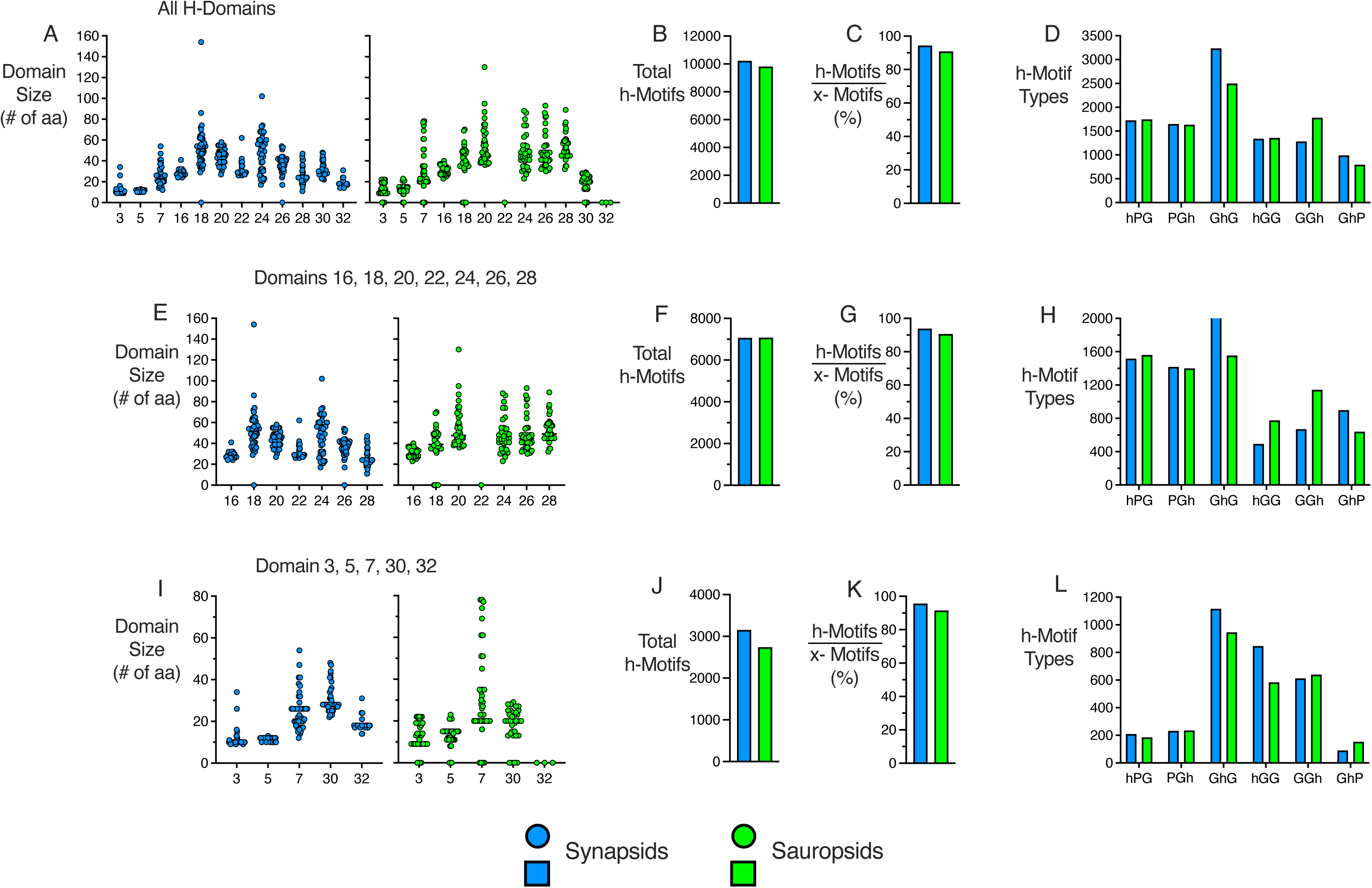
Conservation of total h-motifs and h-motif types in aggregated H-domain sequences. In spite of substantial variability in size of H-domains across individual species of Amniotes, measured as the number of amino acids in the domain (**A**), the collective sum of all h-motifs (h is V, I, L, A, G, F, Y or Q) (**B**) and the ratio of h-motifs to x-motifs (x includes all amino acids with the exception of histidine, tryptophan and methionine) (**C**) were strongly conserved between Synapsid and Sauropsid species. Patterns of distribution of individual h-motif types were also well-conserved (**D**). Similarly strong conservation of these properties were evident when H-domains were separated into those in the ‘replication region’ of tropoelastin (domains 16, 18, 20, 22, 24, 26, and 28) (**E-H**) and those from flanking regions (domains 3, 5, 7, 30, 32) (**I-L**). Previously noted differences in distribution of motif types between these two regions of H-domains were also evident, comparing **H** to **L**.

Strong conservation of these properties was also evident when the H-domains were divided into sub-sets of domains within the replication region (domains 16,18, 20, 22, 24, 26, and 28) (Figures 9E to H), and domains in the flanking regions of tropoelastins (domains 3, 5, 7, 30, and 32) (Figures 10I to L), respectively. Differences in distribution of h-motif types between these sub-sets of H-Domains, previously noted in % motif frequency data (Figure 8C), were also represented in the collective totals of motif types (comparing Figure 10H and L). These data suggest that the specific domain location of these motifs is less important in providing the required properties of tropoelastins than the collective number of the motifs summed over these hydrophobic domains.

## 7. Variance in occurrence and character of exons/domains among Amniote tropoelastins

As noted above, Amniote species were chosen for this analysis because they appear to represent a clade, with easily discerned correspondence of exons/domains across all species. Nevertheless, several differences with respect to exon occurrence and/or character were noted across sub-groups of species. Details of these are provided in Figure 11. This map designates exons that are present (white squares) or absent (black squares) in the genomic sequences of individual species. Criteria for deeming exons absent from the genomic sequence are discussed above (see section 2.1).

**Figure 11.**
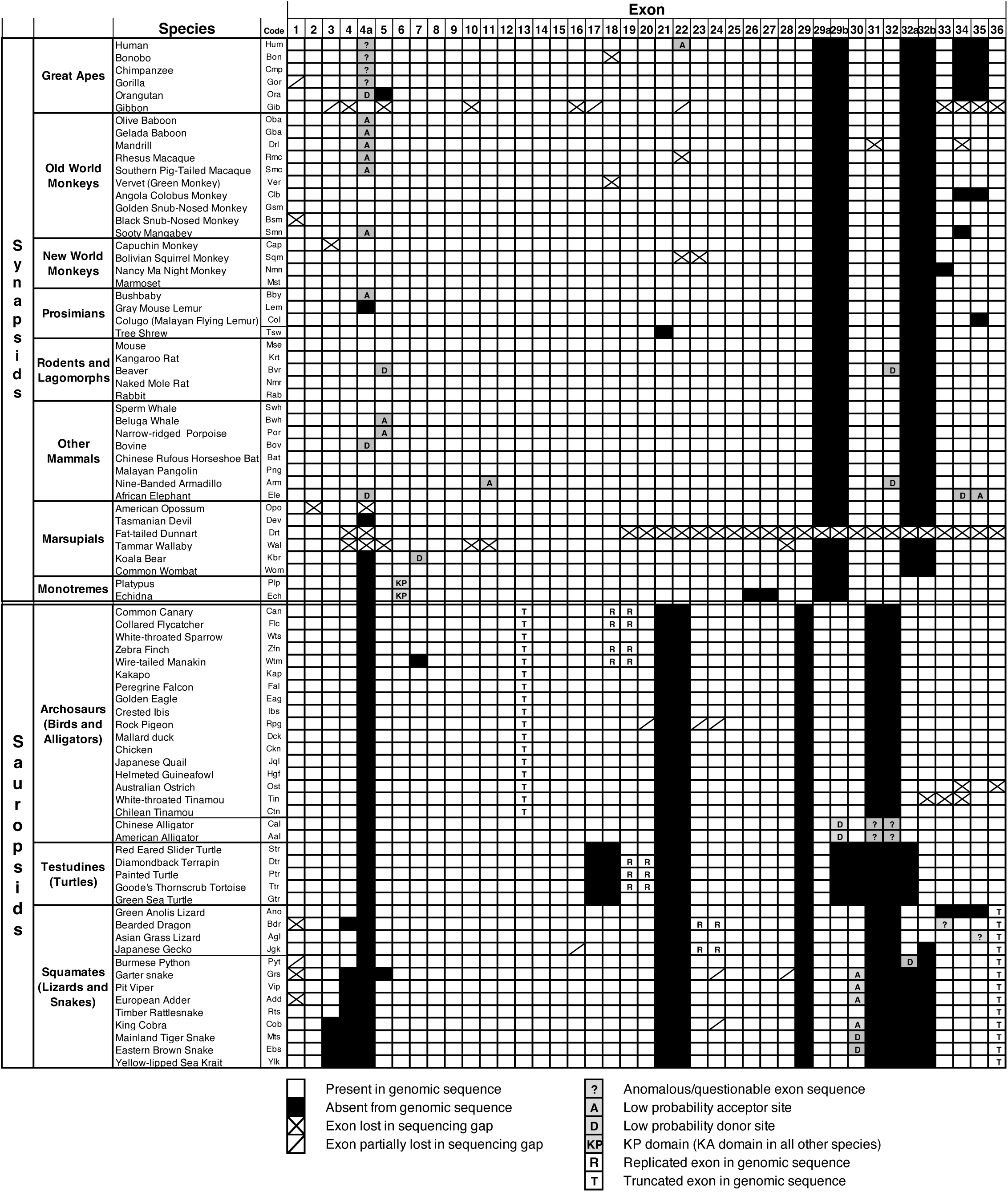
Mapping of variability in occurrence and characteristics of exons in Amniote tropoelastins. Filled squares indicate exons that are present (white) or absent (black) from the genomic sequence. Hatches across squares designate exons with sequence that was partially (single hatch) or completely (double hatch) lost due to sequencing gaps. Exons with anomalous or unexpected sequence relative to the corresponding exons in closely related species are marked ‘?’. Exons with very low probability or otherwise questionable acceptor or donor splice sites are marked ‘A’ or ‘D’ respectively. Exons with truncated sequence are marked ‘T’. Exon pairs that are replicated in the genomic sequence are marked ‘R’. Note that unusual exons that are identified in only a single or a few species (e.g. exon 5a, see text) are not represented in this map.

Exons with sequences partially lost in sequencing gaps (and not retrievable with RefSeq or TSA sequence) are indicated with a single cross-hatch. Exons with sequences completely lost in sequencing gaps are indicated with a double cross-hatch. Exons that are truncated with respect to their expected sequence are marked ‘T’. Exons with unexpected or anomalous sequence relative to corresponding exons in closely related species are indicated ‘?’. Exons recognizable by sequence but with questionable acceptor or donor splice sites are indicated ‘A’ or ‘D’ respectively. Exon pairs that are replicated in the genomic sequence of some species are indicated ‘R’.

### 7.1 Variations in exon occurrence between Synapsids and Sauropsids

The transition between Synapsids (placental and non-placental mammals) and Sauropsids (birds/reptiles) provides the most striking example of modification in exon occurrence in Amniote tropoelastins. Exons 21, 22, 29, 31 and 32 are present in the genomic sequences of all Synapsids. However, in Sauropsid species these exons are all absent, resulting in the disappearance of three KA-type cross-linking domains (21, 29 and 31) and two hydrophobic domains (22 and 32) from the tropoelastin sequences. However, coincident with this transition, four new exons appear in the genomic sequence of Sauropsids (exons 29a, 29b, 32a and 32b), corresponding to two KP-type cross-linking domains (29a and 32a) and two short hydrophobic domains (29b and 32b). These four Sauropsid exons are absent from genomic sequences of all Synapsid tropoelastins, including marsupials. The only exception to this is found in monotremes.

Monotreme species (platypus and echidna) are situated at the interface between Synapsids and Sauropsids (Figure 1). This state appears to be reflected in a transitional pattern of occurrence of these sets of exons. While the expected Synapsid exons, 21, 22, 29, 31, 32 are all present in monotreme tropoelastins, exons 32a and 32b, normally found only in Sauropsid species, are also present in monotremes (Figure 11). Divergences between Synapsids and Sauropsids species (approximately 310 myrs ago), and between marsupials and monotremes (approximately 180 myrs ago) presumably set a timeframe over which this transition in exon usage has taken place.

We also noted that domain sequences of monotremes, particularly in the replicated region, were generally more difficult to identify with corresponding domains of either Sauropsid or Synapsid tropoelastins. Furthermore, unlike all other species, Synapsid or Sauropsid, domain 6 of monotreme tropoelastins contains a KP rather than KA cross-linking motif. This may suggest a more complex evolutionary relationship of monotremes to other Amniote species than that indicated in Figure 1. Unfortunately, available sequence is limited by the small number of extant monotreme species.

### 7.2 Loss of exons 34 and 35 in primates

The loss of exons 34 and 35 from the genomic sequence of human tropoelastin was first noted several years ago, and attributed to an abundance of alu repeats in the region between exons 33 and 36 in human tropoelastin^86,87^. Subsequently Szabo et al.^88^, predominantly using a PCR approach, investigated the presence or absence of these two exons more broadly in genomic sequences of tropoelastins from a variety of human sub-populations, as well as in a limited number of sequences of other primates. They concluded that both exons 34 and 35 were absent from the genomic sequence of all human population sub-groups tested. Their data also suggested that some other primate sequences contained exon 34 but not exon 35, implying that the evolutionary loss of these exons from genomic sequences occurred in a stepwise manner.

In contrast to these earlier findings, our observations (Supplementary Table S6) using a large primate sequence dataset including 5 great apes (hominidae), 10 old world monkeys, 4 new world monkeys and 4 prosimians, clearly show that both exons 34 and 35 are consistently absent from the genomic sequences of tropoelastins of all great apes for which sequence was available (human, bonobo, chimpanzee, gorilla and orangutan). Gibbon tropoelastin sequence for these exons was lost in a sequencing gap.

However, exons 34 and 35 were present with high probability acceptor and donor splice sites in all other primates for which sequence was available. In many of these cases the presence of these exons was also predicted from available supporting data, including RefSeqs and TSA sequences. Exceptions were the Angolan colobus monkey, in which both exons were absent, and the colugo/Malayan flying lemur, in which exon 35 was absent. Exons 34 and 35 were present and predicted in all other Synapsids and Sauropsids, with the exception of the anolis lizard. On the basis of these data, and in contrast to the reports of Szabo et al.^88^ it appears that, with the exceptions noted above, exons 34 and 35 are absent from all great apes but present in the genomic sequences of essentially all other Amniote species.

### 7.3 Expression of Exon 22 in Synapsids

Exon 22 is unusual in that, although present in the genomic sequence, it appears to be spliced out of the majority of the common isoforms of human tropoelastin and rarely, if ever, appears as a protein product in elastic tissues. In the case of human tropoelastin this is probably due, at least in part, to a mutation that abolishes a viable acceptor site for exon 22, resulting in exon skipping. It should be noted that a rare polymorphism in human tropoelastin (rs782509254) is predicted to significantly increase the probability of the acceptor site for exon 22, increasing the likelihood of domain 22, a large hydrophobic domain, being inserted between domains 21 and 23, two cross-linking domains in human tropoelastin. In vitro investigations have suggested that such inclusion of domain 22 may significantly affect functional properties of the protein^18,20^.

The availability of our sequence database provided the opportunity to assess whether consistent splicing out of exon 22 might be exclusive to human tropoelastin or was also the case in other primates (Supplementary Table S7). With the exception of human tropoelastin, high splice site probabilities for exon 22 were predicted not only for the remainder of the great ape sub-group, but also for all 17 other primates for which sequence was available in the database. However, the great ape sub-group is unusual in that exon 22 was spliced out of 36 of the 39 available isoform RefSeqs, compared to only 13 of the 77 available isoform RefSeqs for other subgroups of primates. These data indicate that, in spite of high probability acceptor sites, splicing out of exon 22 may be much more frequent for tropoelastins of great apes compared to other primates, suggesting that trans-acting factors^89^ may be particularly important for regulating the splicing of this exon.

### 7.4 Presence of exon 4a in rodents and other mammals

Exon 4a was first identified in rodents^2^. This exon, situated between exons 4 and 5 in the genomic sequence, codes for a short, hydrophobic domain (e.g. GAGLLGTFGA in mouse). Exon sequences coding for very similar domains can readily be found in the intronic sequence between exons 4 and 5 in essentially all placental mammals (Supplementary Table S8). In many cases, including great apes and old world monkeys, these sequences have high probability splice sites, but are never reported in supporting sequences such as RefSeqs or TSA/EST sequences. Indeed, in great apes the domain sequences show signs of degeneration, as might be expected for an unexpressed exon. In contrast, in new world monkeys and prosimians, the sequence of domain 4a has been predicted in several supporting sequences. Domain 4a expression is also clearly supported in all rodents and lagomorphs, as well as in many (but not all) other non-primate mammals.

A fuller understanding of the distribution and expression of exon 4a in tropoelastins will await more supporting data, particularly transcriptome sequence. However, the currently available data make it clear that the presence and expression of this exon is not restricted to rodents, as previously thought, but includes many other placental mammals. As indicated above, recognizable exon 4a sequence is absent from all marsupials and monotremes, as well as all Sauropsid species.

### 7.5 Rare exons

Additional possible exons have been identified in tropoelastin genomic sequences of some Amniotes. However, because these are generally limited to a single or at most a few species. they have not been included in Figure 11. As an example, exon 5a is situated between exons 5 and 6 and codes for a short, hydrophobic sequence (typically GPGGLA) (Supplementary Table S9). Exon 5a is found only in one sub-group of marsupials, consisting of Tasmanian devil, fat-tailed dunnart and koala bear. Expression of exon 5a in these species is supported by high probability splice sites, as well as by RefSeqs or TSA sequence data. The exon is absent from opossum and wombat tropoelastins, and from monotreme species. Although rare, and limited to only some marsupial species, the independent recognition of this exon in three relatively closely related species suggests that its presence is not the result of sequencing errors or mis-assembly.

### 7.6 Exon truncations

Modifications to tropoelastin exons across Amniote species might be expected, particularly between Synapsid and Sauropsid groups that are separated by more than 300 myrs of independent evolution. The two most striking of these modifications occur within the Sauropsid sub-group and consisted of truncations of a significant portion of otherwise well-conserved domain sequences. These truncations are limited to specific evolutionary lineages but are consistently present within those species groups.

The first example of these is a deletion of 5 amino acids from the C-terminal portion of domain 13, a truncation which is found in all avian members of the archosaur sub-group (Figure 1), but not in alligators/crocodiles (Supplementary Table S10). This truncation can be attributed to a single base polymorphism generating a new, high probability donor site located within the normal sequence of exon 13. This mutation results in the loss of the singlet lysine residue normally present in this domain, with possible consequences for the distribution of covalent cross-links stabilizing the elastin polymer.

A second example of such a truncation is found in exon 36 of squamates (lizards and snakes) and consists of an in-frame deletion of 24 bases resulting in the loss of 8 amino acids from the central portion of this domain. The sequence of domain 36 is otherwise strongly conserved, not only across all other Amniotes (Supplementary Table S11) but also across non-Amniote tropoelastins^2^. Although the C-terminal tetrabasic sequence (RKRK) is retained, the deletion removes both a singlet lysine residue as well as the only two cysteine residues normally found in tropoelastins. This truncation is particularly notable because exon 36 has been thought to have a critical role in assembly of tropoelastin into the elastin polymer, suggesting a unique and significant modification of this process in Squamates.

### 7.7 Additional replications of exon pairs

As indicated in Figure 2, exons/domains 15-29b has been designated as the ‘replicated region’ of tropoelastins because of previous observations that the region appeared to be generated by replication of cross-linking/hydrophobic exon pairs, followed by sequence diversification through mutations, particularly in hydrophobic exons^2^. It was therefore of interest to observe the presence of several clear examples of additional and likely recent replications of exon pairs within the replicated regions of some Amniote tropoelastins. Exons pairs demonstrating such recent replication are labelled ‘R’ in Figure 11, and examples of such replication events are given in Supplementary Table S12.

Multiple replicates of the 18/19 exon pair were detected in the genomic sequences of birds belonging to the passeriformes Order, including species of canary, flycatcher, zebra finch and manakin. No such replications were observed in species belonging to any other avian Orders (Supplementary Figure S3). Replications consisted of 4-fold to 8-fold repeats of the exon pair, and were not only observed in genomic sequences but also supported by predicted RefSeqs. With few exceptions, the replicated sequences were highly similar or even identical to each other at both protein and DNA levels (Supplementary Table S12a), suggesting that the replication event(s) took place relatively recently on an evolutionary timescale. The consistency of this additional replication together with its exclusivity to passeriformes birds argues against any suggestion that such replications may be artifacts of sequence mis-assembly.

Similar multiple replications, in this case of the 19/20 exon pair, were observed in several species of testudines, including diamondback terrapin, painted turtle, thornscrub tortoise, loggerhead turtle and Mexican gopher tortoise, with examples shown in Supplementary Table S12b. Again, genomic sequence replications were supported by RefSeqs, and replicated sequences were highly similar at both the DNA and protein levels. Unfortunately, transcriptome data was not available for any of these species.

The clear evidence of such additional cross-linking/hydrophobic exon pairs within the region covering exons 15 to 29b supports its earlier designation as the ‘replicated region’ of Amniote tropoelastins (Figure 2) and suggests that the types of replication events that generated this region may be on-going in at least some species, perhaps as part of an active remodeling process of tropoelastin sequences.

## 8. General Discussion

Recent advances in understanding how the unusual properties of polymeric elastin arise from the sequence of the monomeric precursor, tropoelastin, have primarily depended on molecular biological and biophysical approaches, modeling sequences or sequence fragments derived from, at most, a few mammalian species. In this study we have taken an alternate approach, analyzing a database of tropoelastin sequences compiled from a wide range of Amniote species, including approximately equal numbers of Synapsids and Sauropsids, sub-groups of Amniotes that have been separated by approximately 300 million years of evolution. The careful curation and comprehensive nature of this database has allowed the identification of characteristics of tropoelastins that have been retained over this extended period of evolution, presumably as a requirement for the fundamental properties of elastic matrices.

Essential properties of elastin include the ability to retain, even in its polymeric form, the high degree of conformational flexibility/structural disorder required to function as a robust entropic elastomer, capable of literally billions of energy-efficient cycles of extension and recoil without mechanical failure. At the same time the sequence of the monomer must also permit a sufficient degree of ordering interactions between monomeric chains to allow for a self-assembly process involving liquid-liquid phase separation (i.e. coacervation) of monomers to form hydrogel-like oligomeric assemblies that will mature into covalently cross-linked fibrillar structures. This ability to maintain a delicate balance between order and disorder would be expected to be a powerful constraining factor on evolutionary permissible sequence characteristics of tropoelastins.

Structural disorder and liquid-liquid phase separation have more recently been identified as properties of a number of intracellular proteins, and the ability of this process to selectively cluster proteins and other macromolecules into transient functional units is now thought to be essential to many biological processes^90–93^. While there are similarities at the biophysical level between retention of conformational flexibility/phase separation in such intra-cellular proteins and tropoelastins, there appears to be less similarity in the sequences associated with these characteristics. This difference may be related to the fact that formation of protein clusters by most intracellular proteins must be completely reversible. Indeed, mutations that limit such reversibility have been associated with a number of pathological consequences^94–96^. In contrast, evolutionary pressures on sequences of tropoelastins have been to form protein clusters which, while initially reversible, normally mature into more stable oligomeric assemblies that can eventually be covalently cross-linked into highly stable polymers.

### 8.1 Conservation of collective properties

Consistent with observations on other low-complexity proteins^38,39^, this study demonstrates that conserved characteristics of tropoelastins include not only preservation of regions of positional sequence, but also conservation of collective or compositional characteristics that describe regions of the protein as a whole. These bulk properties are, of course, derived from the overall sequence of the protein, but are not strictly dependent on conserved linear or positional sequence.

In spite of the extended evolutionary divergence between Synapsid and Sauropsid species groups, coarse-grained properties such as overall amino acid composition, predominance of high proportions of non-polar amino acids (e.g. V, I, L, A, P, G), a paucity of amino acids with negatively charged side chains, and a consistent absence of histidine, tryptophan and internal methionine residues were essentially unchanged between these species groups. More subtle characteristics, including P/G ratios, high proportions of P residues in PG pairs and abundant and well-distributed hPG and hGG motifs, were also conserved. For all species, valine residues were strongly dominant in the h-position of these motifs. Similarly retained was the differential relative density of hGG and hPG motifs in hydrophobic domains of the N-and C-terminal flanking regions (domains 3, 5, 7, 30, 32), as compared to such hydrophobic domains in the replicated region (domains 16, 18, 20, 22, 24, 26, 28), a feature which may be a determinant of the balance of order and disorder in the structure of tropoelastins.

Perhaps the most unexpected example of such collective conservation was the observation that, in spite of large variability across species in the size of individual hydrophobic domains in both replicated and flanking regions of the tropoelastin sequence, the collective totals and patterns of these non-polar motifs were remarkably conserved. This observation indicated that specific domain locations of such h-motifs was less important than the overall sum of h-motifs across these hydrophobic domains

Such collective properties are likely important for the maintenance of structural flexibility in the protein, and for the general ability to undergo liquid-liquid phase separation into condensed oligomeric assemblies, an important early step in the formation of the extracellular elastic matrix.

### 8.2 Conservation of positional sequence

Regions of relatively high levels of cross-species conservation between tropoelastin sequences had previously been noted^2^. However, no attempt was made at that time either to quantify overall conservation across species or to measure differences in degrees of conservation between regions within the protein. Such quantification would first require the construction of a consensus sequence for the protein from a plausible alignment of a large number of tropoelastin sequences. Generation of such an alignment presented a significant challenge, particularly for hydrophobic domains that, in spite of retaining a recognizable ‘style’ of sequence, are characterized by semi-repetitive motifs involving variable replications and many insertions, gaps and deletions. Such sequences were not amenable to analysis by standard alignment software and required manual, domain-by-domain alignment.

The strategy for generating a plausible consensus sequence has been described in detail in section 6. Importantly, this process involved assumptions of both domain correspondence across species and an evolutionary relationship between sequences, recognizing not if, but how these sequences were related. ‘Anchor’ motifs, consisting of hPG and hGG triplets, were particularly useful for sequence alignment of hydrophobic domains.

For consensus sequences constructed from tropoelastin sequences of all Amniotes, position-by-position conservation levels were particularly high in domains 8-14 and in the KA cross-linking domains, as well as in some shorter regions, including domain 33. Indeed, for these sites, conservation approached 100% when V/I/ L and Y/F were treated as equivalent residues. Comparing consensus sequences separately derived from Synapsid and Sauropsid sub-sets of tropoelastin sequences, position-by-position conservation patterns were essentially identical.

The functional significance of such remarkable positional sequence conservation in domains 8-14 of tropoelastin is not clear but may reflect retention of specific sequences serving as sites of interaction with other matrix proteins that take part in in vivo assembly and influence the final architecture of the elastic matrix^1,57–65^. Similarly, conservation of sequences in the regions of KA cross-link formation may be important for subsequent interactions with lysyl oxidase during the cross-linking process. That having been said, the polyalanine sequence context of the lysines involved in KA cross-linking domains is not present in equally well-conserved KP cross-linking domains, suggesting that cross-link formation involving KP domains may require other types of interactions with lysyl oxidase.

When positional sequence conservation was averaged over regions of tropoelastin, domains 8-14 and KA cross-linking domains of the replicated regions consistently had the highest levels of conservation, while hydrophobic domains, particularly those in the replicated region (domains 16, 18, 20, 22, 24, 26, 28) were relatively poorly conserved. This pattern of regional sequence conservation was also consistent between Synapsid and Sauropsid sub-groups. For all three data sets (all Amniotes, Synapsids only, Sauropsids only), analyses of the collective properties of the consensus sequences were in very good agreement with averaged data from the individual sequences contributing to the consensus sequence, confirming that the consensus sequences provided good collective representations of the individual sequences.

Finally, it is important to note the extended region of unusual sequence conservation in the proximal 3’utr of Amniote tropoelastins. This sequence region consistently included target sites for at least three micro-RNAs, including multiple copies of micRNA29 binding sites. The fact that the micRNA29 family has previously been reported to affect expression of extracellular matrix proteins^77–80^, including tropoelastin, and its potential involvement in destabilizing structural elements in the 3’utr of tropoelastin strongly suggests a role in post-transcriptional regulation of elastin production in tissues.

### 8.3 Motif patterns in hydrophobic domains

As indicated above, a relatively low levels of linear positional sequence conservation was a common feature of hydrophobic domains, both from flanking regions of tropoelastins (domains 3, 5, 7, 30 and 32) and from replicated regions (domains 16, 18, 20, 22, 24, 26, 28). Nevertheless, such domains, collectively referred to as H-domains, clearly retained a certain ‘style’ that could be described as generated from tandem repeats and quasi-repeats of motifs rich in PG and GG. Indeed, achieving plausible positional sequence alignments in such domains was facilitated by the use of hPG and hGG ‘anchor’ sequences, where ‘h’ consisted of a non-polar amino acid. This suggested that hPGhGG might describe the fundamental repeated unit, subsequently modified by extensive substitutions, insertions and deletions, accounting for the conservation of sequence ‘style’ in H-domains of all Amniote tropoelastins.

Consistent with this suggestion, for all possible triplets comprising a repeated unit of the putative hPGhGG sequence (xPG, PGx, GxG, xGG, GGx and GxP), non-polar amino acids (V, I, L, A, G, F, Y, Q) were dominant in the ‘x’ position, accounting for >90% of these triplets in H-domain sequences. Moreover, such non-polar triplets provided good coverage of all possible triplets of an hPGhGG repeat sequence. Similar extents and patterns of coverage were also seen when Amniote H-domain sequences were divided into sub-sets of Synapsid and Sauropsid H-domain sequences. For all of these triplets, V, I and L were the favoured amino acids (>75%) in the ‘h’ position, with V particularly dominant.

Earlier studies suggested that the conformational flexibility of elastin sequences was due to a rapid exchange between local beta-turn and polyproline II structural elements in the hydrophobic domains of the protein^6,7,9–13^. This was subsequently confirmed by NMR and molecular dynamics analysis, identifying PG and VG as principal sites of these transient beta-turns^6–8,11–13,43^. The strong conservation of valine residues both proceeding and following such PG pairs may reflect an evolutionary constraint related to preserving the transient nature of these beta turns.

When the H-domain analyses were sub-divided into those in flanking regions (domains 3, 5, 7, 30, 32) and those in the replicated region (domains 16, 18, 20, 22, 24, 26, 28), again frequencies and patterns of triplets were in good agreement between Synapsid and Sauropsid sub-groups. However, the pattern of triplet coverage was different between flanking region and replicated region triplets, reflecting the altered hGG/hPG ratios between these two regions that had been previously observed. A similar analysis of H domains in consensus sequences for all Amniotes, and for Synapsid and Sauropsid sub-groups produced essentially identical results, a further indication that the derived consensus sequences were a good representation of individual tropoelastin sequences. All of these data were consistent with the hypothesis that hPGhGG (where ‘h’ is a non-polar amino acid, predominantly valine, isoleucine and leucine) may represent the fundamental repetitive unit of hydrophobic domains of tropoelastin in all Amniotes.

### 8.4 Exon usage and permitted inter-species evolutionary diversity

While our database identified strongly retained characteristics of tropoelastins, this approach has also allowed recognition of evolutionarily permissible modifications that may tailor the properties of the resulting elastic matrix to specific requirements of species groups. In this respect, it is of interest to point out that, unlike the many collagen genes that provide for a wide array of variations in tissue- and site-specific physical properties, Amniotes have only a single gene for tropoelastin, such that other means must be available for modulation of elastic properties to meet the specific requirements of different tissues and species.

Variations in exon usage in the final protein product clearly provide a mechanism for modification of elastin sequences, with subsequent modulation of both the assembly process and the final properties of the elastic matrix. One way of achieving these variations is through alternate splicing. Extensive alternate splicing was clearly evident in all tropoelastins for which there was sufficient sequence data, and is facilitated by the consistent phase of all intron/exon borders, a feature that is conserved for all tropoelastin genes. This, together with the predominance of alternating hydrophobic and cross-linking exons in the sequence, permits cassette-like removal of one or more of these exons without affecting downstream sequence. In-frame extensions or truncations of exons/domains using alternate donor sites are less common but have also been noted.

The probability of alternate splicing is affected both by cis-sequences at the intron/exon borders and within the intron, as well as by trans-acting factors interacting with the sequence^89^. This allows for a wide variety of permutations and combinations of sequence, all of which conserve the general character of tropoelastin. Moreover, since alternate splicing may not be all-or-none, the possibility of mixed polymers in tissues^22^ provides an additional mechanism for modulation of the final properties of the matrix.

As an example, exon 22, while present in the genomic sequence of all Synapsid tropoelastins is commonly spliced out of tropoelastins of all great ape species, removing a significant hydrophobic spacer between two cross-linking domains. In the case of human tropoelastin, this is likely the result of a mutation leading to loss of a viable acceptor site for this exon. All other primates, including other great apes, have high probability acceptor sites for exon 22. Nevertheless, splicing out of exon 22 was much more frequent in great apes compared to all other primate species, suggesting that regulatory trans-acting factors may also be involved. In vitro modelling of restoration of exon 22 has been shown to have subtle but measurable effects on assembly and elastomeric properties of human tropoelastin^18,20^.

A second process for generation of sequence diversity while retaining essential properties of tropoelastin is through variability in inclusion/exclusion of exons in the genomic sequence itself. Several examples of this were seen in the database. Exons 34 and 35 were absent not only from the human tropoelastin gene, but also from the genomic sequences of all other great apes. In contrast, these exons are present and expressed in essentially all other Synapsid and Sauropsid species. Exons can also be gained between species groups. For example, exon 4a is a short, hydrophobic exon previously identified only in rodent species^2^ but demonstrated in our database to also be present and expressed in several other mammalian species.

The most striking example of generalized exon gain/loss is provided at the interface between Synapsids and Sauropsid species. This transition is marked by the disappearance of exons 21, 22, 29, 31 and 32, generally present in genomic sequences of all Synapsids, and their replacement by exons 29a, 29b, 32a, 32b, generally present in all Sauropsid species. Interestingly, monotremes, a Synapsid species group evolutionarily positioned at the interface between Synapsids and Sauropsids, provide a transitional genomic sequence, including exons 32a and 32b without loss of any Synapsids exons. The evolutionary pathway leading to this significant shift in exon usage between Sauropsid and Synapsid species, and its consequences for the properties of tropoelastins is not clear.

The database has also provided examples of variations between species/species groups as a result of truncations or extensions of sequences within exons, perhaps the most striking of which was the in-frame deletion of 24 bases (8 amino acids) from a central region of exon/domain 36 in all Squamate species, removing both a lysine residue and the only two cysteine residues in this otherwise well-conserved domain.

Sequence variations also included extensions of the ‘replicated region’ of tropoelastin sequences by the introduction of additional replications of a cross-linking/hydrophobic exon pair. Such extensions were relatively rare. However, they were consistent within the passeriformes sub-set of bird species, as well as in some species of testudines. As noted earlier, a general susceptibility to such replication events in this specific region of tropoelastin genes may have an important role in both the history as well as the on-going evolution of tropoelastins. However, specific sequence elements in tropoelastin genes that lead to this susceptibility to internal replication have not been identified.

Removal, inclusion or other modifications to domains by any of these processes will have consequences for positioning of potential cross-linking sites in the protein as well as for spacing between cross-links, both of which can affect modulus and other material properties of polymeric elastin. Such changes may also affect the availability of sites for interactions with other matrix-associated proteins that are thought to play a role in determining the architecture of the elastic matrix in different tissues^1,57–65^.

The database also identifies several other possible opportunities for more subtle sequence modulations with potential effects on the conformational disorder/order balance in tropoelastins. These include mutations affecting hGG/hPG ratios, locations and number of negatively charged residues or even the proportions of I, L, or V residues proceeding and following PG turns in the protein. Such mutations may well have been exploited by species groups to fine-tune properties of the elastic matrix.

Together these examples suggest a rich source of evolutionarily permitted opportunities for subtle variations in the characteristics of elastic matrices, tailoring properties to specific requirements of individual species groups while maintaining the fundamental sequence requirements for assembly, stability and elastomeric properties of elastic matrices. They also provide insights into the evolutionary history of tropoelastin, and suggest possible mechanisms for on-going, dynamic evolution of the protein.

## Acknowledgements

The author wishes to acknowledge the contribution of the many former members of his laboratory who together contributed to enriching our understanding of this unusual and remarkable protein, particularly recognizing Richard Stahl, who played an important initial role in harvesting and curation of tropoelastin sequences and began the long journey of assembly and analysis of the database of sequences.

## References

1. Keeley FW (2021) Structural Proteins: The Biochemistry of Elastin. Encyclopedia of Biological Chemistry III (Third Edition). Joseph Jez, ed. Elsevier, pp 668–689.

2. Keeley FW (2013) Evolution of Elastin in Evolution of the Extracellular Matrix. R. P Mecham and F. W. Keeley, eds., Springer-Verlag Press, pp 73–119.

3. Pometun MS, Chekmenev EY, Wittebort RJ (2004) Quantitative observation of backbone disorder in native elastin. J. Biol. Chem. 279, 7982–7987.

4. Kumashiro KK, Ohgo K, Elliott DW, Kagawa TF, Niemczura WP (2012) Backbone motion in elastin’s hydrophobic domains as detected by 2H NMR spectroscopy. Biopolymers 97, 882– 888.

5. Rauscher S, Pomès R (2012) Structural disorder and protein elasticity. Adv Exp Med Biol 725, 159–183.

6. Rauscher S, Pomès R (2017) The liquid structure of elastin. Elife 6:e26526.

7. Reichheld SE, Muiznieks LD, Keeley FW, Sharpe S (2017) Direct observation of structure and dynamics during phase separation of an elastomeric protein. Proc Natl Acad Sci USA 114, E4408–E4415.

8. Depenveiller C, Baud S, Belloy N, Bochicchio B, Dandurand J, Dauchez M, Pepe A, Pomès R, Samouillan V, Debelle L (2024) Structural and physical basis for the elasticity of elastin. Q Rev Biophys. 2024 Mar 19;57:e3.

9. Tamburro AM, Bochicchio B, Pepe A (2003) Dissection of human tropoelastin: Exon-by-exon chemical synthesis and related conformational studies. Biochemistry 42, 13347–13362.

10. Bochicchio B, Pepe A, Tamburro AM (2008) Investigating by CD the molecular mechanism of elasticity of elastomeric proteins. Chirality 20, 985–994.

11. Ohgo K, Dabalos CL, Kumashiro KK (2018) Solid-state NMR spectroscopy and isotopic labeling target abundant dipeptide sequences in elastin’s hydrophobic domains. Macromolecules 51, 2145–2156.

12. Dabalos CL, Ohgo K, Kumashiro KK (2019) Detection of labile conformations of elastin’s prolines by solid-state nuclear magnetic resonance and Fourier transform infrared techniques. Biochemistry 58, 3848–3860.

13. Reichheld SE, Muiznieks LD, Huynh Q, Wang N, Ing C, Miao M, et al. (2021) The evolutionary background and functional consequences of the rs2071307 polymorphism in human tropoelastin. Biopolymers Feb;112(2):e23414.

14. He D, Miao M, Sitarz EE, Muiznieks LD, Reichheld S, Stahl RJ, et al. (2012) Polymorphisms in the human tropoelastin gene modify in vitro self-assembly and mechanical properties of elastin-like polypeptides. PLoS ONE. 7(9):e46130.

15. Yeo GC, Baldock C, Tuukkanen A, Roessle M, Dyksterhuis LB, Wise SG, et al. (2012) Tropoelastin bridge region positions the cell-interactive C terminus and contributes to elastic fiber assembly. Proc Natl Acad Sci USA 109, 2878–2883.

16. Miao M, Sitarz E, Bellingham CM, Won E, Muiznieks LD, Keeley FW (2013) Sequence and domain arrangements influence mechanical properties of elastin-like polymeric elastomers. Biopolymers 99, 392–407.

17. Yeo GC, Baldock C, Wise SG, Weiss AS (2014) A negatively charged residue stabilizes the tropoelastin N-terminal region for elastic fiber assembly. J Biol Chem 289, 34815–34826.

18. Yeo GC, Tarakanova A, Baldock C, Wise SG, Buehler MJ, Weiss AS (2016) Subtle balance of tropoelastin molecular shape and flexibility regulates dynamics and hierarchical assembly. Sci Adv. 2:e1501145.

19. Yeo GC, Baldock C, Wise SG, Weiss AS (2017) Targeted modulation of tropoelastin structure and assembly. ACS Biomater Sci Eng 3, 2832–2844.

20. Miao M, Reichheld SE, Muiznieks LD, Sitarz EE, Sharpe S, Keeley FW (2017) Single nucleotide polymorphisms and domain/splice variants modulate assembly and elastomeric properties of human elastin. Implications for tissue specificity and durability of elastic tissue. Biopolymers 107:e23007.

21. Tarakanova A, Yeo GC, Baldock C, Weiss AS, Buehler MJ (2018) Molecular model of human tropoelastin and implications of associated mutations. Proc Natl Acad Sci USA 115, 7338–7343.

22. Reichheld SE, Muiznieks LD, Lu R, Sharpe S, Keeley FW (2019) Sequence variants of human tropoelastin affecting assembly, structural characteristics and functional properties of polymeric elastin in health and disease. Matrix Biol. 84, 68–80.

23. Murphy WJ, Pringle TH, Crider TA, Springer MS, Miller W (2007) Using genomic data to unravel the root of the placental mammal phylogeny Genome Res 17, 413–421.

24. Janecka JE, Miller W, Pringle TH, Wiens F, Zitzmann A, Helgen KM, et al. (2007) Molecular and genomic data identify the closest living relative of primates. Science 318, 792–794.

25. Deakin JE, Graves JAM, Rens W (2012) The evolution of marsupial and monotreme chromosomes. Cytogenet Genome Res 137, 113–129.

26. Pyron RA, Burbrink FT, Wiens JJ (2013) A phylogeny and revised classification of Squamata, including 4161 species of lizards and snakes. BMC Evol Biol. 13, 93–145.

27. Field DJ, Gauthier JA, King BL, Pisani D, Lyson TR, Peterson KJ (2014) Toward consilience in reptile phylogeny: miRNAs support an archosaur, not lepidosaur, affinity for turtles. Evolution & Development 16, 189–196.

28. Pozzi L, Hodgson JA, Burrell AS, Sterner KN, Raaum RL, Disotell TR (2014) Primate phylogenetic relationships and divergence dates inferred from complete mitochondrial genomes. Mol Phylogenet Evol 75, 165–183.

29. Jarvis ED, Mirarab S, Aberer AJ, Li B, Houde P, Li C, et al. (2014) Whole genome analyses resolve early branches in the tree of life of modern birds. Science 346, 1320–1331.

30. Crawford NG, Parham JF, Sellas AB, Faircloth BC, Glenn TC, Papenfuss TJ, et al. (2015) A phylogenomic analysis of turtles. Mol Phylogenet Evol 83, 250–257.

31. Brusatte SL, O’Connor JK, Jarvis ED (2015) The origin and diversification of birds. Curr Biol 25, R888–R898.

32. Dawkins R, Wong Y (2016) The Ancestor’s Tale: A Pilgrimage to the Dawn of Life, 2nd Edition, Houghton Mifflin.

33. Foley NM, Springer MS, Teeling EC (2016) Mammal madness: Is the mammal tree of life not yet resolved? Philos Trans R Soc Lond B Biol Sci. Jul 19;371(1699):20150140.

34. Kumar S, Suleski M, Craig JM, Kasprowicz AE, Sanderford M, Li M, et al. (2022) TimeTree 5: An expanded resource for species divergence times. Mol Biol Evol 39, msac174

35. Chung MIS, Miao M, Stahl RJ, Chan E, Parkinson J, Keeley FW (2006) Sequences and domain structures of mammalian, avian, amphibian and teleost tropoelastins: Clues to the evolutionary history of elastins. Matrix Biol 25, 492–504.

36. Miao M, Bruce AEE, Bhanji T, Davis EC, Keeley FW (2007) Differential expression of two tropoelastin genes in zebrafish. Matrix Biol 26, 115–124.

37. Miao M, Stahl RJ, Petersen LF, Reintsch WE, Davis EC, Keeley FW (2009) Characterization of an unusual tropoelastin with truncated C-terminus in the frog. Matrix Biol 28, 432–441.

38. Mier P, Paladin L, Tamana S, Petrosian S, Hajdu-Soltész B, Urbanek A, et al. (2020) Disentangling the complexity of low complexity proteins. Brief Bioinform. 21, 458–472.

39. Lee B, Jaberi-Lashkari N, Calo E (2022) A unified view of low complexity regions (LCRs) across species. eLife 11:e77058.

40. Lillie MA, Piscitelli MA, Vogl AW, Gosline JM, Shadwick RE (2013) Cardiovascular design in fin whales: high-stiffness arteries protect against adverse pressure gradients at depth. J Exp Biol 216 (Pt 14), 2548–2563.

41. Schmelzer CEH, Nagel MBM, Dziomba S, Merkher Y, Sivan SS, Heinz A (2016) Prolyl hydroxylation in elastin is not random. Biochim Biophys Acta 1860, 2169–2217.

42. Bochicchio B, Laurita A, Heinz A, Schmelzer CEH, Pepe A (2013) Investigating the role of (2S,4R)294-hydroxyproline in elastin model peptides. Biomacromolecules 14, 4278−4288.

43. Rauscher S, Baud S, Miao M, Keeley FW, Pomès R (2006) Proline and glycine control protein self-organization into elastomeric or amyloid fibrils. Structure 14, 1667–1676.

44. Muiznieks LD, Weiss AS, Keeley FW (2010) Structural disorder and dynamics of elastin. Biochem Cell Biol 88, 239–250.

45. Roberts S, Dzuricky M, Chilkoti A (2015) Elastin-like polypeptides as models of intrinsically disordered proteins. FEBS Letters 589, 2477–2486.

46. Kozel BA, Wachi H, Davis EC, Mecham RP (2003) Domains in tropoelastin that mediate elastin deposition in vitro and in vivo. J Biol Chem 278, 18491–18498.

47. Muiznieks LD, Keeley FW (2010) Proline periodicity modulates the self-assembly properties of elastin-like polypeptides. J Biol Chem 285, 39779–39789.

48. 48. Muiznieks LD, Cirulis JT, van der Horst A, et al. (2014) Modulated growth, stability and interactions of liquid-like coacervate assemblies of elastin. Matrix Biol 36, 39–50.

49. Muiznieks LD, Reichheld SE, Sitarz EE, Miao M, Keeley FW (2015) Proline-poor hydrophobic domains modulate the assembly and material properties of polymeric elastin. Biopolymers 103, 563–573.

50. Muiznieks LD, Miao M, Sitarz EE, Keeley FW (2016) Contribution of domain 30 of tropoelastin to elastic fiber formation and material elasticity. Biopolymers 105, 267–275.

51. Muiznieks LD, Keeley FW (2017) Biomechanical design of elastic protein biomaterials: A balance of protein structure and conformational disorder. ACS Biomater Sci Eng 3, 661–679.

52. Miao M, Bellingham CM, Stahl RJ, Sitarz EE, Lane CJ, Keeley FW (2003) Sequence and structure determinants for the self-aggregation of recombinant polypeptides modeled after human elastin. J Biol Chem 278, 48553–48562.

53. Tamburro AM, Pepe A, Bochicchio B, Quaglino D, Ronchetti IP (2005) Supramolecular amyloid-like assembly of the polypeptide sequence coded by exon 30 of human tropoelastin. J Biol Chem 280, 2682–2690.

54. Bochicchio B, Pepe A, Delaunay F, Lorusso M, Baud S, Dauchez M (2013) Amyloidogenesis of proteolytic fragments of human elastin. RSC Adv 3, 13273–13285.

55. Reichheld SE, Muiznieks LD, Stahl R, Simonetti K, Sharpe S, Keeley FW (2014) Conformational transitions of the cross-linking domains of elastin during self-assembly. J Biol Chem 289, 10057–10068.

56. Savage KN, Gosline JM (2008) The role of proline in the elastic mechanism of hydrated spider silks. J Exp Biol 211(Pt 12), 1948–1957.

57. Nakamura T, Lozano PR, Ikeda Y, Iwanaga Y, Hinek A, Minamisawa S, et al. (2002) Fibulin-5/DANCE is essential for elastogenesis in vivo. Nature 415, 171–175.

58. Broekelmann TJ, Kozel BA, Ishibashi H, Werneck CC, Keeley FW, Zhang L, et al. (2005) Tropoelastin interacts with cell-surface glycosaminoglycans via its COOH-terminal domain. J Biol Chem 280, 40939–40947.

59. Yanagisawa H, Davis EC (2010) Unraveling the mechanism of elastic fiber assembly: The roles of short fibulins. Int J Biochem Cell Biol 42, 1084–1093.

60. Noda K, Dabovic B, Takagi K, Inoue T, Horiguchi M, Hirai M, et al. (2013) Latent TGF-beta binding protein 4 promotes elastic fiber assembly by interacting with fibulin-5. Proc Natl Acad Sci USA 110, 2852–2857.

61. Pilecki B, Holm AT, Schlosser A, Moeller JB, Wohl AP, Zuk AV, et al. (2015) Characterization of microfibrillar-associated protein 4 (MFAP4) as a tropoelastin- and fibrillin-binding protein involved in elastic fiber formation. J Biol Chem 291, 1103–1114.

62. Kozel BA, Mecham RP (2019). Elastic fiber ultrastructure and assembly. Matrix Biol 84, 31– 40.

63. Shin SJ, Yanagisawa H (2019) Recent updates on the molecular network of elastic fiber formation. Essays Biochem 63, 365–376.

64. Lockhart-Cairns MP, Newandee H, Thomson J, Weiss AS, Baldock C, Tarakanova A (2020) Transglutaminase-mediated cross-linking of tropoelastin to fibrillin stabilises the elastin precursor prior to elastic fibre assembly. J Mol Biol 432, 5736–5751.

65. Schmelzer CEH, Hedtke T, Heinz A (2020) Unique molecular networks: Formation and role of elastin cross-links. IUBMB Life 72, 842–854.

66. Owji H, Nezafat N, Negahdaripour M, Hajiebrahimi A, Ghasemi Y (2018) A comprehensive review of signal peptides: Structure, roles, and applications. Eur J Cell Biol 97, 422–441.

67. Hinek A, Rabinovitch M (1994) 67-kD elastin-binding protein is a protective “companion” of extracellular insoluble elastin and intracellular tropoelastin. J. Cell Biol 126, 563–574.

68. Hinek A, Keeley FW, Callahan J (1995) Recycling of the 67-kDa elastin binding protein in arterial myocytes is imperative for secretion of tropoelastin. Exp Cell Res 220, 312–324.

69. Davis EC, Broekelmann TJ, Ozawa Y, Mecham RP (1998) Identification of tropoelastin as a ligand for the 65-kD FK506-binding protein, FKBP65, in the secretory pathway. J Cell Biol 140, 295–303.

70. Miao M, Reichheld SE, Muiznieks LD, Huang Y, Keeley FW (2013) Elastin binding protein and FKBP65 modulate in vitro self-assembly of human tropoelastin. Biochemistry 52, 7731– 7741.

71. Swee MH, Parks WC, Pierce RA (1995) Developmental regulation of elastin production. Expression of tropoelastin pre-mRNA persists after down-regulation of steady-state mRNA levels. J Biol Chem 270, 14899–14906.

72. Johnson DJ, Robson P, Hew Y, Keeley FW (1995) Decreased elastin synthesis in normal development and in long-term aortic organ and cell cultures is related to rapid and selective destabilization of mRNA for elastin. Circ Res 77, 1107–1113.

73. Hew Y, Grzelczak Z, Lau C, Keeley FW (1999) Identification of a large region of secondary structure in the 3’-untranslated region of chicken elastin mRNA with implications for the regulation of mRNA stability. J Biol Chem 274, 14415–14421.

74. Hew Y, Lau C, Grzelczak Z, Keeley FW (2000) Identification of a GA-rich sequence as a protein-binding site in the 3’-untranslated region of chicken elastin mRNA with a potential role in the developmental regulation of elastin mRNA stability. J Biol Chem 275, 24857– 24864.

75. Hagmeister U, Reuschlein K, März A, Wenck H, Gallinat S, Lucius R, et al. (2012) Poly(A) tail shortening correlates with mRNA repression in tropoelastin regulation. J Dermatol Science 67, 44–50.

76. Procknow SS, Kozel BA (2022) Emerging mechanisms of elastin transcriptional regulation. Am J Physiol Cell Physiol. 323, C666–C677

77. Ott CE, Grünhagen J, Jäger M, Horbelt D, Schwill S, Kallenbach K, et al. (2011) MicroRNAs differentially expressed in postnatal aortic development downregulate elastin via 3′ UTR and coding-sequence binding sites. PLoS ONE 6, e16250.

78. Zhang P, Huang A, Ferruzzi J, Mecham RP, Starcher BC, Tellides G, et al. (2012) Inhibition of microRNA-29 enhances elastin levels in cells haploinsufficient for elastin and in bioengineered vessels. Arterioscler Thromb Vasc Biol 32, 756–759.

79. Jin M, Wu Y, Wang J, Ye W, Wang L, Yin P, et al. (2016) MicroRNA-29 facilitates transplantation of bone marrow-derived mesenchymal stem cells to alleviate pelvic floor dysfunction by repressing elastin. Stem Cell Res Ther 7, 167.

80. Rothuizen TC, Kemp R, Duijs JM, de Boer HC, Bijkerk R, et al. (2016) Promoting Tropoelastin Expression in Arterial and Venous Vascular Smooth Muscle Cells and Fibroblasts for Vascular Tissue Engineering. Tissue Eng Part C Methods 22, 923–931.

81. Di Gregoli K, Mohamad Anuar NN, Bianco R, White SJ, Newby AC, George SJ, et al. (2017) MicroRNA-181b controls atherosclerosis and aneurysms through regulation of TIMP- 3 and elastin. Circ Res 120, 49–65.

82. Autenshlyus AI, Perepechaeva ML, Studenikina AA, Grishanova AY, Lyakhovich VV (2023) Serum miR-181а and miR-25 Levels in Patients with Breast Cancer or Benign Breast Disease. Dokl Biochem Biophys 512, 279–283.

83. Chen J, Liu K, Vadas MA, Gamble JR, McCaughan GW (2024) The Role of the MiR-181 Family in Hepatocellular Carcinoma. Cells 13, 1289.

84. Mills TW, Wu M, Alonso J, Puente H, Charles J, Chen Z, Yoo SH, Mayes MD, Assassi S (2024) Unraveling the role of MiR-181 in skin fibrosis pathogenesis by targeting NUDT21. FASEB J. Sept;38(17):e70022.

85. Zhao J, Wang X, Wu Y, Zhao C (2024) Krüppel-like factor 4 modulates the miR- 101/COL10A1 axis to inhibit renal fibrosis after AKI by regulating epithelial-mesenchymal transition. Ren Fail Dec;46(1):2316259.

86. Indik Z, Yeh H, Ornstein-Goldstein N et al. (1987) Alternative splicing of human elastin mRNA indicated by sequence analysis of cloned genomic and complementary DNA. Proc Natl Acad Sci USA 84, 5680–5684.

87. Bashir MM, Indik Z, Yeh H, Ornstein-Goldstein N, Rosenbloom JC, Abrams W, et al. (1989 Characterization of the complete human elastin gene. Delineation of unusual features in the 5’-flanking region. J Biol Chem 264, 8887–8891.

88. Szabó Z, Levi-Minzi SA, Christiano AM, Struminger C, Stoneking M, Batzer MA, et al. (1999) Sequential loss of two neighboring exons of the tropoelastin gene during primate evolution. J Mol Evol 49, 664–671.

89. Lee Y, Rio DC (2015) Mechanisms and regulation of alternative pre-mRNA splicing. Annu Rev Biochem 84, 291–323.

90. Chong PA, Forman-Kay JD (2016) Liquid-liquid phase separation in cellular signaling systems. Curr Opin Struct Biol 41, 180–186.

91. Banani SF, Lee HO, Hyman AA, Rosen MK (2017) Biomolecular condensates: organizers of cellular biochemistry, Nat Rev Mol Cell Biol 18, 285–298.

92. Murray DT, Kato M, Lin Y, Thurber KR, Hung I, McKnight SL, et al. (2017) Structure of FUS protein fibrils and its relevance to self-assembly and phase separation of low-complexity domains. Cell 171, 615–627.

93. Muiznieks LD, Sharpe S, Pomès R, Keeley FW (2018) Role of liquid-liquid phase separation in assembly of elastin and other extracellular matrix proteins. J Mol Biol 430, 4741–4753.

94. Molliex A, Temirov J, Lee J, Coughlin M, Kanagaraj AP, Kim HJ, et al. (2015) Phase separation by low complexity domains promotes stress granule assembly and drives pathological fibrillization. Cell 163, 123–133.

95. Shin Y, Brangwynne CP (2017) Liquid phase condensation in cell physiology and disease. Science Sep 22;357(6357):eaaf4382.

96. Mathieu C, Pappu RV, Taylor JP (2020) Beyond aggregation: Pathological phase transitions in neurodegenerative disease. Science 370, 56–60.

