## Supplemental material for "Evolutionary Constraints on Positional Sequence, Collective Properties and Sequence Style of Tropoelastin Dictated by Fundamental Requirements for Formation and Function of the Extracellular Elastic Matrix"

Fred W. Keeley

**Supplementary Material**

#### Supplementary Table S1 Sequence Source Information and Curation Notes

| Species | Code[1] | Taxonomy | Sequence Group[2] | Genomic Sequence (strand) | Position of First Exon (exon #)[3] | Supporting Sequences[4] | Curation Annotations[5] |
|---|---|---|---|---|---|---|---|
| Human (*Homo sapiens sapiens*) | Hum | Class: Mammalia<br>Order: Primates<br>Family: Hominidae | Great Apes | NG_009261.1<br>(+) | 5092 (1) | NM_000501.3; NM_001278939.1; P15502.3 | Full sequence coverage. Exon 22 present in genomic sequence but has a low probability acceptor site. |
| Bonobo (*Pan paniscus*) | Bon | Class: Mammalia<br>Order: Primates<br>Family: Hominidae | Great Apes | NC_27875.1<br>(+) | 81178099 (1) | XM_024929386.1 | Exon 18 lost in sequencing gap |
| Chimpanzee (*Pan troglodytes*) | Cmp | Class: Mammalia<br>Order: Primates<br>Family: Hominidae | Great Apes | NW_019932819.1<br>(+) | 3487327 (1) | GABF01011800.1; GABF01011799.1; GABF01011797.1; GABE01015702.1 | Full sequence coverage |
| Gorilla (*Gorilla gorilla gorilla)* | Gor | Class: Mammalia<br>Order: Primates<br>Family: Hominidae | Great Apes | NC_018431.2<br>(+) | 72670482 (1 ) | XM_019030426.1 | Exon 1 partially lost in sequencing gap |
| Orangutan *(Pongo abelii*) | Ora | Class: Mammalia<br>Order: Primates<br>Family: Hominidae | Great Apes | NC_036910.1<br>(+) | 8009725 (1) | XM_024250278.1; XM_024250277.1; CR630179.1 | Full sequence coverage |
| Gibbon (*Nomascus leucogenys*) | Gib | Class: Mammalia<br>Order: Primates<br>Family: Hominidae | Great Apes | NW_004087903.1<br>(-) NW_003502016.1<br>(-) | NW_004087903.1: 39799594 (1)<br>NW_003502016.1: 73453 (17) | XM_012504028.1 | Exons 3, 4, 5, 10, 16, 17, 22, 32-36 partially or fully lost in sequencing gaps |
| Olive Baboon *(Papio anubis)* | Oba | Class: Mammalia<br>Order: Primates<br>Family: Cercopithecidae | Old World Monkeys | NC_018154.2<br>(+) | 45305189 (1) | AC091092.3 (exons 20-23) XM_017956649.1 *Papio hamadryas*: GAAH01004259.1; GAAH01004260.1; GAAH01004261.1; GAAH01004262.1; GAAH01000562.1 | Full sequence coverage |
| Gelada Baboon (*Theropithecus gelada*) | Gba | Class: Mammalia<br>Order: Primates<br>Family: Cercopithecidae | Old World Monkeys | NC_037670.1<br>(-) | 8966016 (1) | XM_025380695.1 | Full sequence coverage |
| Mandrill *(Mandrillus leucophaeus*) | Drl | Class: Mammalia<br>Order: Primates<br>Family: Cercopithecidae | Old World Monkeys | NW_012101632.1<br>(+) | 1959253 (1) | XR_001005635.1 | Exons 31, 34 lost in sequencing gaps |
| Rhesus Macaque (*Macaca mulatta*) | Rmc | Class: Mammalia<br>Order: Primates<br>Family: Cercopithecidae | Old World Monkeys | NC_027895.1<br>(-) | 43411431 (1) | | |
| Southern Pig-Tailed Macaque (*Macaca* | Smc | Class: Mammalia<br>Order: Primates<br>Family: Cercopithecidae | Old World Monkeys | NW_012013022.1<br>(-) | 9925899 (1) | XM_011715349.1 | Full sequence coverage |
| Vervet [Green Monkey] (*Chlorocebus* | Ver | Class: Mammalia<br>Order: Primates<br>Family: Cercopithecidae | Old World Monkeys | NC_023669.1<br>(-) | 9048207 (1) | XM_008018944.1 | Full sequence coverage. Some mis-ordered exons. Exon 18 absent from genomic sequence |
| Angola Colobus Monkey (*Colobus angolensis palliatus*) | Clb | Class: Mammalia<br>Order: Primates<br>Family: Cercopithecidae | Old World Monkeys | NW_012113142.1<br>(-) | 3234039 (1) | XM_011930905.1 | Exons 34, 35 lost in sequencing gap |

| | | | | | | | |
|---|---|---|---|---|---|---|---|
| Golden Snub-Nosed Monkey (*Rhinopithecus roxellana*) | Gsm | Class: Mammalia<br>Order: Primates<br>Family: Cercopithecidae | Old World Monkeys | NW_010828282.1 (+) | 3258858 ( 1) | XM_010387470.1 | Full sequence coverage |
| Black Snub-Nosed Monkey (*Rhinopithecus bieti*) | Bsm | Class: Mammalia<br>Order: Primates<br>Family: Cercopithecidae | Old World Monkeys | NW_016804227.1 (+) | 231922 (2) | | Exon 1 lost in sequencing gap |
| Sooty Mangabey (*Cercocebus atys*) | Smn | Class: Mammalia<br>Order: Primates<br>Family: Cercopithecidae | Old World Monkeys | NW_012001517.1 (+) | 8317143 (1) | XM_012076413.1 | Exon 34 absent from genomc sequence |
| Capuchin mA20:H23onkey (Cebus capucinus | Cap | Class: Mammalia<br>Order: Primates<br>Family: Cebidae | New World Monkeys | NW_016107397.1 (-) | 197269 (1) | XM_017533409.1 | Exon 3 lost in sequencing gap |
| Bolivian Squirrel Monkey (*Saimiri boliviensis* | Sqm | Class: Mammalia<br>Order: Primates<br>Family: Cebidae | New World Monkeys | NW_003943666.1 (-) | 6184372 (1) | XM_010344835.1 | Exons 22, 23 lost in sequencing gap |
| Nancy Ma's Night Monkey (*Aotus nancymaae* ) | Nmn | Class: Mammalia<br>Order: Primates<br>Family: Aotidae | New World Monkeys | NW_018487315.1 (+) | 3108180 (1) | XM_012449205.2; XR_001106924.1 | Exon 33 absent from genomc sequence |
| Marmoset (*Callithrix jacchus* ) | Mst | Class: Mammalia<br>Order: Primates<br>Family: Callitrichidae | New World Monkeys | NTIC01011112.1 (+)<br>NW_003185370.1 (-) | NTIC01011112.1: 191663849 (1)<br>NW_003185370.1: 141785 (30) | XM_008990251.2; XM_008981486.2; GAMT01012626.1; GAMT01024455.1; GAMT01017644.1 | Full sequence coverage |
| Bushbaby (*Otolemur garnettii*) | Bby | Class: Mammalia<br>Order: Primates<br>Family: Galagidae | Prosimians | NW_003852472.1 (+) | 2836354 (1) | XM_023515738.1 | Full sequence coverage |
| Gray Mouse Lemur (*Microcebus murinus*) | Lem | Class: Mammalia<br>Order: Primates<br>Family: Cheirogaleidae | Prosimians | NC_033676.1 (+) | 33362407 (1) | XM_012742255.2 | Full sequence coverage |
| Colugo (Malayan Flying Lemur) (*Galeopterus variegatus*) | Col | Class: Mammalia<br>Order: Dermoptera<br>Family: Cynocephalidae | Prosimians | Class: Mammalia<br>Order: Rodentia<br>Family: Muridae | 193062 (1) | JMZW01096893.1; XM_008581617.1 | Exons mis-ordered but recoverable. Exon 35 absent from genomic sequence |
| Tree Shrew *(Tupaia chinensis)* | Tsw | *Class: Mammalia*<br>*Order: Scandentia*<br>*Family: Tupalidae* | Prosimians | NW_006159934.1 (+) | 912463 (1) | XM_006155027.2; XM_006155028.2 | Exon 21 absent from genomic sequence |
| Mouse *(Mus musculus)* | Mse | Class: Mammalia<br>Order: Rodentia<br>Family: Muridae | Rodents and Lagomorphs | NC_000071.6 (-) | 134747242 (1) | XM_006504360.3; GGAD01003544.1: GGAD01003543.1; GGAD01026245.1; GEAB01124321.1; CJ185511; BQ951050; BQ265900; CF738958; BQ919729; CK387220; BI698060; CF724314 | Full sequence coverage |
| Kangaroo Rat *(Dipodomys ordii)* | Krt | Class: Mammalia<br>Order: Rodentia<br>Family: Heteromyidae | Rodents and Lagomorphs | NW_012267380.1 (+) | 257995 (1) | XM_013033168 | Full sequence coverage |
| Beaver *(Castor canadensis)* | Bvr | Class: Mammalia<br>Order: Rodentia<br>Family: Castoridae | Rodents and Lagomorphs | NW_017869557.1 (-) | 786099 (1) | XM_020164796.1; XM_020164826.1; XM_020164834.1 | Full sequence coverage |

| | | | | | | | |
|---|---|---|---|---|---|---|---|
| *Naked Mole Rat (Heterocephalus glaber)* | Nmr | Class: Mammalia<br>Order: Rodentia<br>Family: Heterocephalidae | Rodents and Lagomorphs | NW_004624740.1 (-) | 13886201 (1) | XM_013070497.1; XM_013070498.1; XM_013070499.1; XM_013070500.1; XM_013070501.1;, XM_013070502.1 | Full sequence coverage |
| Rabbit *(Oryctolagus cuniculus )* | Rab | Class: Mammalia<br>Order: Lagomorpha<br>Family: Leporidae | Rodents and Lagomorphs | NW-003159391.1 (-) | 681354 (1) | XM_008249104.2; XM_017337950.1; XM_012927495.1; GBCN01145537.1; GBCT01118148.1; GBCT01143583.1; GBCJ01181743.1; GBCT01143583.1; GBCM01049244.1 | Full sequence coverage |
| Sperm Whale *(Physeter catodon)* | Swh | Class: Mammalia<br>Order: Artiodactyla<br>Family: Physeteridae | Other Mammals | NW_019873548.1 (-) | 142559775 (1) | RNA RefSeqs: XM_024125848.1 XM_028485383.1 | Full sequence coverage |
| Beluga Whale *(Delphinapterus leucas)* | Bwh | Class: Mammalia<br>Order: Artiodactyla<br>Family: Monodontidae | Other Mammals | NW_019160893.1 (+) | 5249515 (1) | XM_022588081.1; GGBT01050051.1; GGBT01042559.1; GGBT01038646.1; GGBT01061683.1; GGBT01012384.1, | Exon 5 acceptor site mutation |
| Narrow ridged Finless Porpoise *(Neophocaena [illegible]* | Por | Class: Mammalia<br>Order: Artiodactyla<br>Family: Phocoenidae | Other Mammals | NW_020173405.1 (-) | 1201101 (1) | XM_024747173.1; GBYP01021118.1; GBYL01027692.1 | Exon 5 acceptor site mutation |
| Bovine *(Bos taurus)* | Bov | Class: Mammalia<br>Order: Artiodactyla<br>Family: Bovidae | Other Mammals | NC_037352.1 (-) | 33310399 (1) | XM_005225254.4; GGVG01114211.1; GGVI01113361.1; EV632554.1; EH172866.1; EE358551.1; EV626801.1; EE250041.1. | Full sequence coverage |
| Chinese rufous horseshoe bat *(Rhinolophus sinicus)* | Bat | Class: Mammalia<br>Order: Chiroptera<br>Family: Rhinolophidae | Other Mammals | NW_017739330.1 (-) | 756949 (1) | XM_019755001.1; GAVY01036303.1 | Full sequence coverage |
| Malayan Pangoln *(Manis javanica)* | Png | Class: Mammalia<br>Order: Philodota<br>Family: Manidae | Other Mammals | NW_016527738.1 (-) | 86151 (1) | XM_017662454.1; GEUV01071946.1 | Full sequence coverage |
| Nine-Banded Armadillo *(Dasypus novemcinctus)* | Arm | Class: Mammalia<br>Order: Cingulata<br>Family: Dasypodidae | Other Mammals | NW_004480668.1 (-) | 322259 (1) | XM_012526232.2 | Full sequence coverage |
| African Elephant *(Loxodonta africana)* | Ele | Class: Mammalia<br>Order: Proboscidea<br>Family: Elephantidae | Other Mammals | NW_003573465.1 (-) | 17172181 (1) | XM_010596331.2. | Full sequence coverage |
| American Opossum *(monodelphis domestica)* | Opo | Class: Mammalia<br>Order: Didelphimorphia<br>Family: Didelphidae | Marsupials | NC_008802.1 (+) | 530804482 (1) | ENSMODT00000042404.1 | Exon, 2, 4a lost in sequencing gaps |
| Tasmanian Devil *(Sarcophilus harrisii)* | Dev | Class: Mammalia<br>Order: Dasyuromorphia<br>Family: Dasyuridae | Marsupials | GL952809.1 (-)<br>NW_003839709.1 (-) | GL952809.1: 72258 ( 1) | GEDN01068181.1; GEDN01068183.1; GEDN01066991.1; GEDN01066990.1 | Full sequence coverage |

| | | | | | | | |
|---|---|---|---|---|---|---|---|
| Fat-tailed Dunnart *(Sminthopsis crassicaudata)* | Drt | Class: Mammalia<br>Order: Dasyuromorphia<br>Family: Dasyuridae | Marsupials | | | GFCN01213778.1; GFCN01213782.1; GFCN01213779.1 | Partial coverage from TSA sequences |
| Tammar Wallaby *(Macropus eugenii)* | Wal | Class: Mammalia<br>Order: Diprotodontia<br>Family: Macropodidae | Marsupials | Assembled from: ABQO020549863.1 ABQO020216569.1, ABQO020049922.1, ABQO021105351.1 ABQO020607756.1, ABQO020663305.1, ABQO020774403.1, ABQO020829952.1, ABQO020885501.1, ABQO020941050.1, | ABQO020549863.1: 329 (1) | FY622604.1; EX201067.1; EX202095.1 | Genomic sequence manually assembled. Exons 4, 4a, 5, 10, 11, 28 lost in sequencing gaps |
| Koala Bear *(Phascolarctos cinereus)* | Kbr | Class: Mammalia<br>Order: Diprotodontia<br>Family: Phascolarctidae | Marsupials | NW_018344061.1 (+) | 8184449 (1) | XM_021005681.1 | Full sequence coverage |
| Common Wombat *(Vombatus ursinus)* | Wom | Class: Mammalia<br>Order: Diprotodontia<br>Family: Vombatidae | Marsupials | NW_020954546.1 (+) | 13394064 (1) | XM_027861968.1 | Full sequence coverage |
| Platypus *(Ornithorhynchus anatinus)* | Plp | Class: Mammalia<br>Order: Monotremata<br>Family: Ornithorhynchidae | Monotremes | NC_041744.1 (-) | 3009822 (1) | PTTO01000885.1; XM_029082364.1; XM_007669854.1 | Full sequence coverage. Domain 6 is a KP cross-linking domain rather than the KA cross-linking domain present in all other species. Unexpected presence of exons 32a, 32b |
| Echidna *(Tachyglossus aculeatus)* | Ech | Class: Mammalia<br>Order: Monotremata<br>Family: Tachyglossidae | Monotremes | NC_052082.1 (+) JADRJE010000005.1 (+) | JADRJE010000005.1 (1) | XM_038759310.1; XM_038759311.1 ; XM_038759313.1; XM_038759313.1 XM_038759311.1 | Full sequence coverage. Domains 26 and 27 absent from genomic sequence. Domain 6 is a KP cross-linking domain rather than the KA cross-linking domain present in all other species. Unexpected presence of exons 32a, 32b. |
| Common Canary *(Serinus canaria)* | Can | Class: Aves<br>Order: Passeriformes<br>Family: Fringillidae | Birds | NW_007931150.1 (+) | 739887 (1) | XM_018919278.1 | Full sequence coverage. First instance of absence of exons 21, 22, 29, 31, 32. First appearance of exons 29a, 29b. Exons 32a and 32b present. Genomic sequence includes 4 replicates of exons 18/19. |
| Collared Flycatcher *(Ficedula albicollis)* | Flc | Class: Aves<br>Order: Passeriformes<br>Family: Muscicapidae | Birds | NC_021690.1 (+) | 743958 (1) | XM_016303078.1. SRR1021712.10424662.1 (golden flycatcher) | Full sequence coverage. Exon 35 recovered from golden flycatcher. Genomic sequence includes 6 replicates of exons 18/19. |
| White-throated Sparrow *(Zonotrichia albicollis)* | Wts | Class: Aves<br>Order: Passeriformes<br>Family: Passerellidae | Birds | NW_005081737.1 (-) | 631502 (2) | GBBC01076568.1; GBBB01044180.1; GBBC01048991.1;GBBC01048312.1; | Full sequence coverage. No apparent replication of exons 18/19 in genomic sequence. |

| | | | | | | | |
|---|---|---|---|---|---|---|---|
| Zebra Finch *(Taeniopygia guttata)* | Zfn | Class: Aves<br>Order: Passeriformes<br>Family: Estrildidae | Birds | MUGN01000955.1 (+) | 726767 (1) | NC_011483.1; GGLD01153872.1; GGLD01072803.1; GGLD01074195.1; GGLD01074196.1 | Full sequence coverage. Genomic sequence includes 5 replicates of exons 18/19. |
| Wire-tailed Manakin *(Pipra filicauda)* | Wtm | Class: Aves<br>Order: Passeriformes<br>Family: Pipridae | Birds | NW_020895081.1 (+) | 478935 ( 1) | XM_027729277.1 | Exon 7 absent from genomic sequence. Genomic sequence includes 6 replicates of exons 18/19. |
| Kakapo *(Strigops habroptilus)* | Kap | Class: Aves<br>Order: Psittaciformes<br>Family: Strigopidae | Birds | RXXE01000054.1 (-) | 150519 (1) | | Full sequence coverage. |
| Peregrine Falcon *(Falco peregrinus)* | Fal | Class: Aves<br>Order: Falciniformes<br>Family: Falconidae | Birds | NW_004930114.1 (-) | 81529 (1) | XM_027784749.1. | Full sequence coverage |
| Golden Eagle *(Aquila chrysaetos canadensis)* | Eag | Class: Aves<br>Order: Accipitriformes<br>Family: Accipitridae | Birds | NW_011950985.1 (-) | 2316538 (1) | XM_011594512.1. | Full sequence coverage |
| Crested Ibis (Nipponia nippon) | Ibs | Class: Aves<br>Order: Pelicaniformes<br>Family: | Birds | NW_009000314.1 (-) | 5488826 (1) | XM_009464831.1. | Full sequence coverage |
| Rock Pigeon *(Columba livia)* | Rpg | Class: Aves<br>Order: Columbiformes<br>Family: Columbidae | Birds | NW_004974336.1 (+) | 247421 (1) | XM_021282733.1 | Partial loss of exons 20, 23, 24 in sequencing gaps. |
| Chicken *(Gallus gallus)* | Ckn | Class: Aves<br>Order: Galliformes<br>Family: Phasianidae | Birds | NC_006106.5 (+) | 943342 (1) | XM_025141904.1; DR425139.1; BI389853.1; BI389865.1; DR428148.1; DR428148.1; AM066490.1; BM485600.1 | Full sequence coverage |
| Japanese Quail *(Coturnix japonica)* | Jql | Class: Aves<br>Order: Galliformes<br>Family: Phasianidae | Birds | NW_015438850.1 (-) | 6709289 (1) | XM_015880998.1 | Full sequence coverage |
| Helmeted Guineafowl *(Numida meleagris)* | Hgf | Class: Aves<br>Order: Galliformes<br>Family: Numididae | Birds | NC_034426.1 (+) | 768588 (1) | XM_021415950.1; GBYG01022455.1; GBYG01022456.1 | Full sequence coverage |
| Mallard duck *(Anas platyrhynchos)* | Dck | Class: Aves<br>Order: Anseriformes<br>Family: Anitidae | Birds | NOIJ01000248.1 (-) | 678014 (1) | XM_013096136.1; GGZN01057217.1 (Muscovey Duck) | Full sequence coverage |
| Australian Ostrich *(Struthio camelus australis)* | Ost | Class: Aves<br>Order: Struthioniformes | Birds | NW_009271464.1 (-) | 4576446 (1) | XM_009678162.1; XR_694588.1 | Exons 34 and 36 lost in sequencing gaps. |
| White throated Tinamou *(Tinamus guttatus)* | Tin | Class: Aves<br>Order: Tinamiformes<br>Family: Tinamidae | Birds | NW_010576439.1 (-) | 51689 (1) | | Exons 32b, 33, 34, lost in sequencing gaps. |

| | | | | | | | |
|---|---|---|---|---|---|---|---|
| Chilean Tinamou *(Nothoprocta perdicaria)* | Ctn | Class: Aves<br>Order: Tinamiformes<br>Family: Tinamidae | Birds | NW_020455932.1 (-) | 2983394 (1) | XM_026050155.1 . | Full sequence coverage |
| Chinese Alligator *(Alligator sinensis)* | Cal | Class: Reptilia<br>Order: Crocodilia<br>Family: Alligatoridae | Alligators | NW_005842661.1 (+) | 556561 (1) | RNA RefSeq: XM_006033788.3 | Full sequence coverage. |
| American Alligator *(Alligator mississippiensis)* | Aal | Class: Reptilia<br>Order: Crocodilia<br>Family: Alligatoridae | Alligators | NW_017709491.1 (-) | 683450 (1) | RNA RefSeq: XM_019486312. | Full sequence coverage. |
| Red Eared Slider Turtle *(Trachemys scripta elegans)* | Str | Class: Reptilia<br>Order: Testudines<br>Family: Emydidae | Turtles | Only transcriptome sequence available | | JW458827.1; JW458828.1; JW458829.1; JW458830.1 | Full sequence coverage. Exons 17, 18, 29b, 30, 32a absent from genomic sequence (characteristic of all turtles). |
| Diamondback Terrapin *(Malaclemys terrapin terrapin)* | Dtr | Class: Reptilia<br>Order: Testudines<br>Family: Emydidae | Turtles | MDXI01019100.1 (-) | 122856 (1) | GEXU01006778.1; GEXU01006774.1; GEXS01000297.1; GEXU01006776.1 | Full sequence coverage. Genomic sequence includes 6 replicates of exons 19/20. |
| Painted Turtle *(Chrysemys picta bellii)* | Ptr | Class: Reptilia<br>Order: Testudines<br>Family: Emydidae | Turtles | NW_007281577.1 (-) | 1134120 (1) | XM_024111351.1; XM_024111352.1; XM_024111353.1 | Full sequence coverage. Genomic sequence includes 5 replicates of exons 19/20. |
| Goode's Thornscrub Tortoise *(Gopherus evgoodei)* | Ttr | Class: Reptilia<br>Order: Testudines<br>Family:Testudinidae | Turtles | NC_044338.1 (+) | 23775472 (1) | RNA RefSeq: XM_030536861.1 | Full sequence coverage. Genomic sequence includes 5 replicates of exons 19/20. |
| Green Sea Turtle *(Chelonia mydas)* | Gtr | Class: Reptilia<br>Order: Testudines<br>Family: Chelonidae | Turtles | NW_006660285.1, (+ ) and NW_006592343.1, (+) | NW_006660285.1 :528106 (1) NW_006592343.1 : 2895 (20) | XM_027829142.1; XR_003565464.1; GGMX01028276.1; GGMX01047413.1 | Full sequence coverage. |
| Green Anolis Lizard *(Anolis carolinus)* | Ano | Class: Reptilia<br>Order: Squamata<br>Family: Polychrodidae | Lizards and Snakes | NW_003339591.1 (+) | 2008 ( 2) | XM_016998779.1; GBDW01030679.1; GADN01012596.1; GAEC01007362.1; GBDW01011564.1 | Regions of exons 33-35 have unusual sequence in both Ano and Bdr, but not in other squamates. Exon 35 is anaomalous KA cross-linking exon in Ano, whereas Bdr has both expected KP cross-linking exon as well as the anomalous KA exon. These exons are confirmed by supporting seqeunces data. Central region of exon 36 is deleted (characteristic of all lizards and [illegible] |

| | | | | | | | |
|---|---|---|---|---|---|---|---|
| Bearded Dragon *(Pogona vitticeps)* | Bdr | Class: Reptilia Order: Squamata Family: Agamidae | Lizards and Snakes | NW_018150994.1 (-) | 49279 (2) | XM_020801655.1 | Exon 1 lost in sequencing gap. Exon 4 absent from genomic sequence. Genomic sequence includes 4 replicates of exons 23/24. Both KA and KP versions of exon 35 present in Bdr (see Ano notes), and sequences |
| Asian Grass Lizard *(Takydromus sexlineatus)* | Agl | Class: Reptilia Order: Squamata Family: Lacertidae | Lizards and Snakes | | | GEMF01016334.1; GEMF01016335.1 | Full sequence coverage. Central region of exon 36 deleted, as in other squamates. |
| Japanese Gecko *(Gekko japonicus)* | Jgk | Class: Reptilia Order: Squamata Family: Gekkonidae | Lizards and Snakes | NW_015167538.1 (+) | 557680 (1) | | Exon 16 partially lost in sequencing gap. Exon 8 truncated (charactistics of all snakes). Genomic sequence includes 2 replicates of exons 23/24 as well a a replicate of exon 27. Exon 32b absent from genomic sequence (also characteristic of all snakes) |
| Burmese Python *(Python bivittatus)* | Pyt | Class: Reptilia Order: Squamata Family: Pythonidae | Lizards and Snakes | NW_006533746.1 (-) and NW_006535511.1 (-) | NW_006533746.1 : 45887 (1) NW_006535511.1 : 134939 (7) | XR_456222.1; XR_001559950.1; XM_015889598.1; XM_015889599.1; XM_015889600.1; XM_025172369.1 | Full sequence coverage. Exons 32a and 32b absent from genomic sequence (characteristic of all snakes). |
| Garter snake *(Thamnophis sirtalis*) | Grs | Class: Reptilia Order: Squamata Family: Colubridae | Lizards and Snakes | NW_013658800.1 (+) and LFLD01067291.1 (-) | NW_013658800.1 : 143304 (2) LFLD01067291.1: 3975 (33) | LFLD01107771.1; LFLD01107777.1; XM_014065579.1; XM_014059253.1; GDKU01023374.1; GDKU01023373.1; GDKU01023375.1; GDKU01023377.1 | Exon 1 lost in sequencing gap. Exon 28 partially lost in sequencing gap. Exons 4 and 5 absent from genomic sequence. |
| Pit Viper *(Protobothrops mucrosquamatus)* | Vip | Class: Reptilia Order: Squamata Family: Viperidae | Lizards and Snakes | NW_015386484.1 (+) | 254912 (1) | XM_015812804.1 | Full sequence coverage. Exon 4 absent from genomic sequence. |
| European Adder *(Vipera berus berus)* | Add | Class: Reptilia Order: Squamata Family: Viperidae | Lizards and Snakes | JTGP01063474.1 (-) JTGP01063473.1 (-) | JTGP01063474.1: 11323 (2) JTGP01063473.1: 33871 (9) | | Exon 1 lost in sequencing gap. Exon 4 absent from genomic sequence. |
| Timber Rattlesnake *(Crotalus horridus)* | Rts | Class: Reptilia Order: Squamata Family: Viperidae | Lizards and Snakes | LVCR01001158.1 (+) | 78946 (1) | | Full sequence coverage. Exon 4 absent from genomic sequence. |
| Jararaca Snake *(Bothrops jararaca)* | Jar | Class: Reptilia Order: Squamata Family: Viperidae | Lizards and Snakes | Only transcriptome sequence available | | GFJM01021698.1; GFJM01043838.1 | Partial sequence only. |

| | | | | | | | |
|---|---|---|---|---|---|---|---|
| Bazilian Lancehead Snake *(Bothrops moojeni)* | Blh | Class: Reptilia Order: Squamata Family: Viperidae | Lizards and Snakes | Only transcriptome sequence available | | GFWW01069819.1; GFWW01069818.1; GFWW01024308.1; GFWW01024307.1 | Partial sequence only. |
| King Cobra *(Ophiophagus hannah)* | Cob | Class: Reptilia Order: Squamata Family: Elapidae | Lizards and Snakes | AZIM01092440.1 (+) and AZIM01000051.1 (+) | AZIM01092440.1: 113 (1) AZIM01000051.1: 111033 (2) | GDRF01026118.1; GDRF01015782.1 | Exon 24 partially lost in sequencing gap. Exons 3 and 4 absent from genomic sequence. |
| Mainland Tiger Snake *(Notechis scutatus)* | Mts | Class: Reptilia Order: Squamata Family: Elapidae | Lizards and Snakes | NW_020716649.1 (-) | 6682002 (1) | XM_026671669.1 | Full sequence coverage. Exons 3 and 4 absent from genomic sequence. |
| Eastern Brown Snake *(Pseudonaj textilis)* | Ebs | Class: Reptilia Order: Squamata Family: Elapidae | Lizards and Snakes | NW_020769348.1 (+) | 8405278 (1) | XM_026708436.1 | Full sequence coverage. Exons 3 and 4 absent from genomic sequence. |
| Yellow-lipped Sea Krait *(Laticauda colubrina)* | Ylk | Class: Reptilia Order: Squamata Family: Elapidae | Lizards and Snakes | BHFR01000347.1 (+) | 1761311 (1) | | Full sequence coverage. Exons 3 and 4 absent from genomic sequence. |
| Painted Coral Snake *(Micrurus corallinus)* | Pcs | Class: Reptilia Order: Squamata Family: Elapidae | Lizards and Snakes | Only transcriptome sequence available | | IACJ01127362.1; IACJ01127361.1; IACJ01162889.1; IACJ01127359.1 | Partial sequence only. |
| South American Coral Snake *(Micrurus lemniscatus)* | Scs | Class: Reptilia Order: Squamata Family: Elapidae Sub-Family: | Lizards and Snakes | Only transcriptome sequence available | | IACK01205291.1; IACK01178129.1 | Partial sequence only. |

1. See Table S2 for species listed by alphabetized code
2. Species groups used for subsequent data analysis
3. Base position for start of sequence (initial exon #)
4. Supporting sequences can include other genomic sequence, RNA RefSeqs, EST sequences and transcriptome sequences
5. Curation Annotations

Full sequence coverage: Genomic sequence with supporting sequence includes all expected exons.
Lost in sequencing gap: Exons may be present but are masked in a sequencing gap (N) and not recovered in supporting sequences.
Absent from genomic sequence: Exons not present in spite of no apparent sequencing gaps.
Other unusual features may be noted.

Supplementary Table S2
Alphabetized Listing of Species by Three-letter Code

| Code | Species | Species Group |
|---|---|---|
| Aal | American Alligator *(Alligator mississippiensis)* | Archosaurs |
| Add | European Adder *(Vipera berus berus)* | Squamates |
| Agl | Asian Grass Lizard *(Takydromus sexlineatus)* | Squamates |
| Ano | Green Anolis Lizard *(Anolis carolinus)* | Squamates |
| Arm | Nine-Banded Armadillo *(Dasypus novemcinctus)* | Other Mammals |
| Bat | Chinese rufous horseshoe bat *(Rhinolophus sinicus)* | Other Mammals |
| Bby | Bushbaby *(Otolemur garnettii)* | Prosimians |
| Bdr | Bearded Dragon *(Pogona vitticeps)* | Squamates |
| Blh | Brazilian Lancehead Snake *(Bothrops moojeni*) | Squamates |
| Bon | Bonobo *(Pan paniscus)* | Great Apes |
| Bov | Bovine *(Bos taurus)* | Other Mammals |
| Bsm | Black Snub-Nosed Monkey *(Rhinopithecus bieti)* | Old World Monkeys |
| Bvr | Beaver *(Castor canadensis)* | Other Mammals |
| Bwh | Beluga Whale *(Delphinapterus leucas)* | Other Mammals |
| Cal | Chinese Alligator *(Alligator sinensis)* | Archosaurs |
| Can | Common Canary *(Serinus canaria)* | Archosaurs |
| Cap | Capuchin monkey *(Cebus capucinus imitator)* | New World Monkeys |
| Ckn | Chicken *(Gallus gallus)* | Archosaurs |
| Clb | Angola Colobus Monkey *(Colobus angolensis palliatus)* | Old World Monkeys |
| Cmp | Chimpanzee *(Pan troglodytes)* | Great Apes |
| Cob | King Cobra *(Ophiophagus hannah)* | Squamates |
| Col | Colugo (Malayan Flying Lemur) *(Galeopterus variegatus)* | Prosimian |
| Ctn | Chilean Tinamou *(Nothoprocta perdicaria)* | Archosaurs |
| Dck | Mallard Duck *(Anas platyrhynchos)* | Archosaurs |
| Dev | Tasmanian Devil *(Sarcophilus harrisii)* | Marsupials |
| Drl | Mandrill *(Mandrillus leucophaeus)* | Old World Monkeys |
| Drt | Fat-tailed Dunnart *(Sminthopsis crassicaudata)* | Marsupial |
| Dtr | Diamondback Terrapin *(Malaclemys terrapin terrapin)* | Testudines |
| Eag | Golden Eagle *(Aquila chrysaetos canadensis)* | Archosaurs |
| Ebs | Eastern Brown Snake *(Pseudonaj textilis)* | Squamates |
| Ech | Echidna *(Tachyglossus aculeatus)* | Monotremes |
| Ele | African Elephant *(Loxodonta africana)* | Other Mammals |
| Fal | Peregrine Falcon *(Falco peregrinus)* | Archosaurs |
| Flc | Collared Flycatcher *(Ficedula albicollis)* | Archosaurs |
| Gba | Gelada Baboon *(Theropithecus gelada***)** | Old World Monkeys |
| Gib | Gibbon *(Nomascus leucogenys)* | Great Apes |

| | | |
|---|---|---|
| Gor | Gorilla *(Gorilla gorilla gorilla)* | Great Apes |
| Grs | Garter snake *(Thamnophis sirtalis)* | Squamates |
| Gsm | Golden Snub-Nosed Monkey *(Rhinopithecus roxellana)* | Old World Monkey |
| Gtr | Green Sea Turtle *(Chelonia mydas)* | Testudines |
| Hgf | Helmeted Guineafowl *(Numida meleagris)* | Archosaurs |
| Hum | Human *(Homo sapiens sapiens)* | Great Apes |
| Ibs | Crested Ibis *(Nipponia nippon)* | Archosaurs |
| Jar | Jararaca Snake *(Bothrops jararaca)* | Squamates |
| Jgk | Japanese Gecko *(Gekko japonicus)* | Squamates |
| Jql | Japanese Quail *(Coturnix japonica)* | Archosaurs |
| Kap | Kakapo *(Strigops habroptilus)* | Archosaurs |
| Kbr | Koala Bear *(Phascolarctos cinereus)* | Marsupials |
| Krt | Kangaroo Rat *(Dipodomys ordii)* | Rodent |
| Lem | Gray Mouse Lemur *(Microcebus murinus)* | Prosimians |
| Mse | Mouse *(Mus musculus)* | Rodents |
| Mst | Marmoset *(Callithrix jacchus)* | New World Monkeys |
| Mts | Mainland Tiger Snake *(Notechis scutatus)* | Squamates |
| Nmn | Nancy Ma's Night Monkey *(Aotus nancymaae)* | New World Monkeys |
| Nmr | Naked Mole Rat *(Heterocephalus glaber)* | Rodents |
| Oba | Olive Baboon *(Papio anubis)* | Old World Monkeys |
| Opo | American Opossum *(Monodelphis domestica)* | Marsupials |
| Ora | Orangutan *(Pongo abelii)* | Great Apes |
| Ost | Australian Ostrich *(Struthio camelus australis)* | Archosaurs |
| Pcs | Painted Coral Snake *(Micrurus corallinus)* | Squamates |
| Plp | Platypus *(Ornithorhynchus anatinus)* | Monotremes |
| Png | Malayan Pangolin (Scaly Anteater *(Manis javanica)*) | Other Mammals |
| Por | Narrow-ridged Finless Porpoise *(Neophocaena* | Other Mammals |
| Ptr | Painted Turtle *(Chrysemys picta bellii)* | Testudines |
| Pyt | Burmese Python *(Python bivittatus)* | Squamates |
| Rab | Rabbit *(Oryctolagus cuniculus )* | Rodents |
| Rmc | Rhesus Macaque *(Macaca mulatta)* | Old World Monkeys |
| Rpg | Rock Pigeon *(Columba livia)* | Archosaurs |
| Rts | Timber Rattlesnake *(Crotalus horridus)* | Squamates |
| Scs | South American Coral Snake *(Micrurus lemniscatus)* | Squamates |
| Smc | Southern Pig-Tailed Macaque *(Macaca nemestrina)* | Old World Monkeys |
| Smn | Sooty Mangabey *(Cercocebus atys)* | Old World Monkeys |
| Sqm | Bolivian Squirrel Monkey *(Saimiri boliviensis boliviensis)* | New World Monkeys |
| Str | Red Eared Slider Turtle *(Trachemys scripta elegans)* | Testudines |
| Swh | Sperm Whale *(Physeter catodon)* | Other Mammals |
| Tin | White throated Tinamou *(Tinamus guttatus)* | Archosaurs |

| | | |
|---|---|---|
| Tsw | Tree Shrew *(Tupaia chinensis)* | Prosimians |
| Ttr | Goode's Thornscrub Tortoise *(Gopherus evgoodei)* | Testudines |
| Ver | Vervet (Green Monkey) *(Chlorocebus sabaeus)* | Old World Monkeys |
| Vip | Pit Viper *(Protobothrops mucrosquamatus)* | Squamates |
| Wal | Tammar Wallaby *(Macropus eugenii)* | Marsupials |
| Wom | Common Wombat *(Vombatus ursinus)* | Marsupials |
| Wtm | Wire-tailed Manakin *(Pipra filicauda)* | Archosaurs |
| Wts | White-throated Sparrow *(Zonotrichia albicollis)* | Archosaurs |
| YlK | Yellow-lipped Sea Krait *(Laticauda colubrina)* | Squamates |
| Zfn | Zebra Finch *(Taeniopygia guttata)* | Archosaurs |

Supplementary Table S3

Amino Acid Compositions and Bulk Properties of Amniote Tropoelastins[1]

(Residues/100)

A

| | Synapsids | | | | | | Sauropsids | | | | | | | | |
|---|---|---|---|---|---|---|---|---|---|---|---|---|---|---|---|
| | All Primates | | | All Other Mammals | | | Archosaurs | | | Testudines | | | Squamates | | |
| | Mean | SD | n | Mean | SD | n | Mean | SD | n | Mean | SD | n | Mean | SD | n |
| ala (A) | 21.7 | 0.86 | 23 | 20.7 | 1.11 | 19 | 17.0 | 0.86 | 18 | 15.5 | 0.26 | 5 | 14.0 | 1.92 | 13 |
| arg (R) | 0.8 | 0.11 | 23 | 1.0 | 0.21 | 19 | 0.8 | 0.13 | 18 | 1.1 | 0.13 | 5 | 1.3 | 0.14 | 13 |
| asn (N) | 0.0 | 0.00 | 23 | 0.0 | 0.05 | 19 | 0.1 | 0.07 | 18 | 0.0 | 0.00 | 5 | 0.2 | 0.12 | 13 |
| asp (D) | 0.1 | 0.06 | 23 | 0.2 | 0.15 | 19 | 0.0 | 0.07 | 18 | 0.0 | 0.00 | 5 | 0.0 | 0.05 | 13 |
| cys (C) | 0.3 | 0.01 | 23 | 0.3 | 0.02 | 19 | 0.3 | 0.02 | 18 | 0.3 | 0.01 | 5 | 0.0 | 0.00 | 13 |
| gln (Q) | 1.2 | 0.14 | 23 | 1.2 | 0.17 | 19 | 1.3 | 0.18 | 18 | 1.9 | 0.07 | 5 | 0.9 | 0.33 | 13 |
| glu (E) | 0.2 | 0.08 | 23 | 0.1 | 0.16 | 19 | 0.0 | 0.06 | 18 | 0.0 | 0.07 | 5 | 0.0 | 0.04 | 13 |
| gly (G) | 31.2 | 1.03 | 23 | 33.7 | 1.97 | 19 | 33.8 | 1.38 | 18 | 33.3 | 1.22 | 5 | 38.9 | 1.96 | 13 |
| his (H) | 0.0 | 0.00 | 23 | 0.0 | 0.00 | 19 | 0.0 | 0.00 | 18 | 0.0 | 0.00 | 5 | 0.0 | 0.00 | 13 |
| ile (I) | 2.1 | 0.31 | 23 | 2.3 | 0.55 | 19 | 2.2 | 0.47 | 18 | 2.1 | 0.14 | 5 | 3.5 | 0.67 | 13 |
| leu (L) | 5.5 | 0.41 | 23 | 5.7 | 0.51 | 19 | 5.8 | 0.53 | 18 | 6.5 | 0.10 | 5 | 5.9 | 1.11 | 13 |
| lys (K) | 4.8 | 0.20 | 23 | 4.9 | 0.27 | 19 | 4.7 | 0.23 | 18 | 4.9 | 0.05 | 5 | 4.1 | 0.45 | 13 |
| met (M) | 0.0 | 0.00 | 23 | 0.0 | 0.04 | 19 | 0.0 | 0.00 | 18 | 0.0 | 0.00 | 5 | 0.0 | 0.04 | 13 |
| phe (F) | 2.1 | 0.21 | 23 | 2.1 | 0.61 | 19 | 2.3 | 0.55 | 18 | 1.0 | 0.06 | 5 | 1.0 | 0.20 | 13 |
| pro (P) | 12.2 | 0.43 | 23 | 11.5 | 0.57 | 19 | 14.0 | 1.22 | 18 | 12.5 | 0.40 | 5 | 12.3 | 0.81 | 13 |
| ser (S) | 1.0 | 0.22 | 23 | 1.2 | 0.43 | 19 | 0.6 | 0.23 | 18 | 1.1 | 0.30 | 5 | 1.0 | 0.34 | 13 |
| thr (T) | 1.5 | 0.28 | 23 | 1.5 | 0.52 | 19 | 1.2 | 0.39 | 18 | 2.2 | 0.20 | 5 | 2.1 | 0.33 | 13 |
| trp (W) | 0.0 | 0.00 | 23 | 0.0 | 0.00 | 19 | 0.0 | 0.00 | 18 | 0.0 | 0.00 | 5 | 0.0 | 0.00 | 13 |
| tyr (Y) | 2.2 | 0.21 | 23 | 2.6 | 0.59 | 19 | 1.5 | 0.67 | 18 | 3.9 | 0.04 | 5 | 3.2 | 0.53 | 13 |
| val (V) | 13.3 | 1.01 | 23 | 11.0 | 1.64 | 19 | 14.4 | 1.79 | 18 | 13.6 | 0.42 | 5 | 11.7 | 1.53 | 13 |
| Predicted MW | 65045 | 2447 | 23 | 65389 | 2814 | 19 | 62507 | 3343.7 | 18 | 54901 | 897 | 5 | 66828 | 6120 | 13 |
| pI | 10.38 | 0.05 | 23 | 10.36 | 0.11 | 19 | 10.64 | 0.164 | 18 | 10.22 | 0.02 | 5 | 10.41 | 0.078 | 13 |
| GRAVY[2] | 0.682 | 0.046 | 23 | 0.565 | 0.084 | 19 | 0.645 | 0.056 | 18 | 0.511 | 0.014 | 5 | 0.481 | 0.071 | 13 |
| % non-polar[3] | 86.03 | 0.72 | 23 | 84.99 | 0.86 | 19 | 87.3 | 1.158 | 18 | 83.5 | 0.38 | 5 | 86.2 | 1.484 | 13 |
| D+E[4] | 0.26 | 0.11 | 23 | 0.29 | 0.25 | 19 | 0.07 | 0.096 | 18 | 0.03 | 0.074 | 5 | 0.03 | 0.085 | 13 |
| K+R[5] | 5.54 | 0.19 | 23 | 5.84 | 0.26 | 19 | 5.47 | 0.301 | 18 | 6.01 | 0.16 | 5 | 5.42 | 0.535 | 13 |
| Y+F[6] | 4.22 | 0.26 | 23 | 4.66 | 0.29 | 19 | 3.81 | 0.297 | 18 | 4.93 | 0.06 | 5 | 4.20 | 0.578 | 13 |

B

| | All Amniotes | | | All Synapsids | | | All Sauropsids | | |
|---|---|---|---|---|---|---|---|---|---|
| | Mean | StDev | n | Mean | StDev | n | Mean | StDev | n |
| ala (A) | 18.71 | 3.18 | 78 | 21.28 | 1.09 | 42 | 15.71 | 1.91 | 36 |
| arg (R) | 0.93 | 0.24 | 78 | 0.86 | 0.18 | 42 | 1.01 | 0.28 | 36 |
| asn (N) | 0.05 | 0.09 | 78 | 0.01 | 0.04 | 42 | 0.10 | 0.10 | 36 |
| asp (D) | 0.08 | 0.11 | 78 | 0.14 | 0.11 | 42 | 0.02 | 0.06 | 36 |
| cys (C) | 0.22 | 0.10 | 78 | 0.26 | 0.01 | 42 | 0.18 | 0.14 | 36 |
| gln (Q) | 1.21 | 0.31 | 78 | 1.20 | 0.16 | 42 | 1.21 | 0.42 | 36 |
| glu (E) | 0.09 | 0.11 | 78 | 0.14 | 0.12 | 42 | 0.03 | 0.06 | 36 |
| gly (G) | 33.81 | 2.95 | 78 | 32.33 | 1.98 | 42 | 35.53 | 2.98 | 36 |
| his (H) | 0.00 | 0.00 | 78 | 0.00 | 0.00 | 42 | 0.00 | 0.00 | 36 |
| ile (I) | 2.42 | 0.68 | 78 | 2.23 | 0.44 | 42 | 2.65 | 0.83 | 36 |
| leu (L) | 5.76 | 0.65 | 78 | 5.61 | 0.46 | 42 | 5.95 | 0.78 | 36 |
| lys (K) | 4.68 | 0.37 | 78 | 4.82 | 0.24 | 42 | 4.51 | 0.44 | 36 |
| met (M) | 0.00 | 0.02 | 78 | 0.01 | 0.03 | 42 | 0.00 | 0.02 | 36 |
| phe (F) | 1.88 | 0.65 | 78 | 2.08 | 0.43 | 42 | 1.66 | 0.78 | 36 |
| pro (P) | 12.47 | 1.19 | 78 | 11.85 | 0.60 | 42 | 13.20 | 1.31 | 36 |
| ser (S) | 0.95 | 0.37 | 78 | 1.07 | 0.35 | 42 | 0.80 | 0.34 | 36 |
| thr (T) | 1.58 | 0.50 | 78 | 1.52 | 0.40 | 42 | 1.64 | 0.60 | 36 |
| trp (W) | 0.00 | 0.00 | 78 | 0.00 | 0.00 | 42 | 0.00 | 0.00 | 36 |
| tyr (Y) | 2.40 | 0.84 | 78 | 2.35 | 0.46 | 42 | 2.45 | 1.14 | 36 |
| val (V) | 12.77 | 1.92 | 78 | 12.28 | 1.74 | 42 | 13.34 | 1.98 | 36 |
| Predicted MW | 64190 | 4464 | 78 | 65201 | 2592 | 42 | 63011 | 5771 | 36 |
| pI | 10.43 | 0.16 | 78 | 10.37 | 0.08 | 42 | 10.50 | 0.20 | 36 |
| GRAVY[2] | 0.600 | 0.097 | 78 | 0.629 | 0.088 | 42 | 0.567 | 0.098 | 36 |
| % non-polar[3] | 85.94 | 1.42 | 78 | 85.56 | 0.94 | 42 | 86.38 | 1.74 | 36 |
| D+E[4] | 0.17 | 0.19 | 78 | 0.27 | 0.18 | 42 | 0.05 | 0.09 | 36 |
| K+R[5] | 5.61 | 0.36 | 78 | 5.68 | 0.26 | 42 | 5.53 | 0.43 | 36 |
| Y+F[6] | 4.28 | 0.48 | 78 | 4.42 | 0.35 | 42 | 4.11 | 0.55 | 36 |

1 Compositions exclude signal peptides. Proline values are sum of proline and hydroxyprol

2 Grand Average of Hydropathy

3 Sum of Ala, Gly, Ile, Leu. Pro and Val

4 Sum of Asp and Glu

5 Sum of Lys and Arg

6 Sum of Tyr and Phe

Supplementary Table S4
Domain 1 (Signal Peptide) Sequence Alignment of Amniote Tropoelastins

| | Species Group | Species | Sequence |
|---|---|---|---|
| Synapsids | Great Apes | Hum | MAGLTAAAPRPGVLLLLL-----SILHPSRPG |
|  |  | Bon | MAGLTAAAPRPGVLLLLL-----SILHPSRPG |
|  |  | Cmp | MAGLTAAAPRPGVLLLLL-----SILHPSRPG |
| | | Gor | /////////////////////////////RPG |
|  |  | Ora | MAGLTAAAPRPGVLLLLL-----SILHPSRPG |
|  |  | Gib | MAGLTAAAPRPGVLLLLL-----SILHPSRPG |
|  | Old World Monkeys | Oba | MAGLTAAALRPGVLLLLL-----SILHPSRPG |
|  |  | Gba | MAGLTAAAPRPGVLLLLL-----SILHPSRPG |
|  |  | Drl | MAGLTAAAPRPGVLLLLL-----SILHPSRPG |
|  |  | Rmc | MAGLTAAAPRPGVLLLLL-----SILHPSRPG |
|  |  | Smc | MAGLTAAAPRPGVLLLLL-----SILHPSRPG |
|  |  | Ver | MAGLTAAAPRPGVLLLLL-----CILHPSRPG |
|  |  | Clb | MAGLTAAAPRPGVLLLLL-----SILHPSRPG |
|  |  | Gsn | MAGLTAAAQRPGVLLLLL-----SILHPSRPG |
|  |  | Smn | MAGLTAAAPRPGVLLLLL-----SILHPSRPG |
|  | New World Monkeys | Cap | MAGLTAAAPRPGVLLLLL-----SILHPSRPG |
|  |  | Sqm | ///////APRPGVLLLLL-----SILHPSRPG |
|  |  | Nmn | MAGLTAAVPRPGVLLLLL-----SILHPSRPG |
|  |  | Mst | MAGLTAAVPWPGVLLLLL-----SILHPSRPG |
|  | Prosimians | Bby | MAGLTAAALRPGVLLLLL-----SILHPSRPG |
|  |  | Lem | MAGLTAAALRPGVLLLLL-----SVLHSSRSG |
|  |  | Col | MAGLSAAAPQPGVLLLLL-----AVLHPSQPG |
|  |  | Tsw | MAGLTAATPRPGVLLLLL-----SILHPSKPG |
|  | Rodents and Lagomorphs | Mse | MAGLTAVVPQPGVLLILLL----NLLHPAQPG |
|  |  | Krt | MAGLSAAAQHPGFLLFLL-----SLLHPAQPG |
|  |  | Bvr | MAGLTAVAPQPGVLLLLL-----NLLHPVQPG |
| | | Nmr | MAGLTAAAPRPGPGALLALPLLLCLLRPAQPG |
|  |  | Rab | MAGVTAAAAPRPGLLLLLL----SLLHPSQPG |
|  | Other Mammals | Swh | MAGPTAAALRPGALLLLL-----CALHPSQPG |
|  |  | Bwh | MAGPTAAALWPGALLLLL-----CILHPSQPG |
|  |  | Por | MAGPTAAALWPGALLLLL-----CILHPSQPG |
|  |  | Bov | MAGLTAAARRPGVLLLLL-----CILQPSQPG |
|  |  | Bat | MAGPTAAALRPGVLLLLL-----SIVQPSQPG |
|  |  | Png | MAGLTAAALRPGVLLLLL-----SVVRPSQPG |
|  |  | Arm | MAGLTAAALRPGVLLLLL-----AALHPSRPG |
|  |  | Ele | MAGPTAAALRPGVLLLLL-----STLHLVQPG |
|  | Marsupials | Opo | MASRTALSLRPGVFLLLL-----SILHPTRQG |
|  |  | Dev | MASRTALSLRAGAFLLLLL----SILQPTRQG |
|  |  | Drt | MASRTALSLRTGACLLLLL----SILQPTRQG |
|  |  | Wal | MASRTALSLRPGVFLLLL-----SILHPTRQG |
|  |  | Kbr | MASRTALPLRPGVFLLLLL----SILQPTRQG |
|  |  | Wom | MASRTALPLRPGVILLLL-----SILQPTRQG |
|  | Monotremes | Plp | MATRTVGFCLLGAIFLLLLL------HPSEQG |
|  |  | Ech | MATRTVGFCLLGAIFLLLLL------HPSEQG |
| Sauropsids | Archosaurs | Can | MARQAAAPLLPGVLLLL------SILPATQQG |
|  |  | Flc | MARQAAAPLLPGVLLLL------SILPATQQG |
|  |  | Wts | MARQAAAPLLPGVLLLL------SILPATQQG |
|  |  | Zfn | MARQAAAPLLPGVLLLLL-----SILPATQQG |
|  |  | Wtm | MARQAAAPLLPGVLLLLF-----SILPATQQG |
|  |  | Kap | MARQAAAPLLPGVLLL-F-----SILPASQQG |
|  |  | Fal | MARQAAAPLLPGVLLL-F-----SILPATQQG |
|  |  | Kes | MARQAAAPLLPGVLLL-F-----SILPATQQG |
|  |  | Eag | MARRAAAPLLPGVLLL-F-----SILPASQQG |
|  |  | Ibs | MARQAAAPLLPGVLLL-F-----SILPASQQG |
|  |  | Rpg | MARQAAAPLLPGVLLLLF-----SILPASQQG |
|  |  | Ckn | MARQAAAPLLPGVLLL-F-----SILPASQQG |
|  |  | Jql | MARQAAAPLLPGVLLL-F-----SILPASQQG |
|  |  | Hgf | MARQAAAPLLPGVLLL-F-----SILPASQQG |
|  |  | Dck | MAKQAAAPLLPGVLLL-F-----SILPASQQG |
|  |  | Ost | MARQPAAPLFPGVLLL-F-----SILPASQQG |
|  |  | Tin | MARQPAAPLFPGVLLL-F-----SILPASQQG |
|  |  | Ctn | MARQPAAPLFPGVLLL-F-----SILRASQQG |
|  |  | Cal | MARQLAASLFPGVLLLL------SILQASWQG |
|  |  | Aal | MARQLAASLFPGVLLLL------SILQASWQG |
|  | Testudines | Str | MARQLAASLLPGVLLLL------AILPASRQG |
|  |  | Dtr | MARQLAASLLPGVLLLL------AILPASRQG |
|  |  | Ptr | MARQLAASLLPGVLLLL------AILPASRQG |
|  |  | Ttr | MARQLAASLLPGVLLLLL-----AILPTSRQG |
|  |  | Gtr | MARQLAASLLPGVLLLL------AILPASRQG |
| | Squamates | Ano | MAKPWTSSLLSGAAGLLLLLL--SSLPASWQG |
|  |  | Agl | MARLRTPSFLPGAVTVLLL----CTLPASWQG |
| | | Jgk | MASLRTPASLPRLLRGVAFLLL-AGLPASWQG |
| | | Pyt | //////////////PGAPVLRL-SGPRASWQG |
|  |  | Vip | MARLRTSFLLPGVLVLLL-----SSLPASWQG |
|  |  | Rts | MARLRTSFLLPGVLVLLL-----SSLPVSWQG |
|  |  | Jar | MARLRTSFLLPGVLVLLL-----SSLPVSWQG |
|  |  | Blh | MARLRTSFLLPGVLALLL-----SSLPVSWQG |
|  |  | Cob | MARLRTSSLLPGVLVLLL-----SSLPASWQG |
|  |  | Mts | MARLRTSSLLPGVLVLLL-----SSLPASWQG |
|  |  | Ebs | MARLRTSSLLPGVLVLLL-----SSLPASWQG |
|  |  | Ylk | MARLRTSSLLPGVLVLLL-----SSLPASWQG |
|  |  | Pcs | MARLRTSSLLPGVLVLLL-----SSLPASWQG |

Notes
Spaces inserted for alignment purposes indicated as '-'
Residue positions lost to sequencing gaps indicated as '/'
In all cases /SignalP-6.0 predicted signal peptide cleavage sites before the final G of Domain 1

Supplementary Table S5

Relative Arrangement of micRNA Target Sites and Polyadenylation Sites in 3'utrs of Amniote Tropoelastins

| | | Species | # PolyA Sites | Bases to 1st pA | Bases to 2nd pA | Bases to 3rd pA | Bases to 4th pA | Bases to 5th pA | Arrangement of micRNA and polyA sites |
|---|---|---|---|---|---|---|---|---|---|
| Synapsids | Great Apes | Hum | 2 | 951 | 1181 | | | | m29-m181-m29-m29p-m101-pA-pA |
| | | Cmp | 2 | 953 | 1185 | | | | m29-m181-m29-m29p-m101-pA-pA |
| | | Bon | 2 | 953 | 1184 | | | | m29-m181-m29-m29p-m101-pA-pA |
| | | Gor | 2 | 954 | 1186 | | | | m29-m181-m29-m29p-m101-pA-pA |
| | | Ora | 2 | 953 | 1182 | | | | m29-m181-m29-m29p-m101-pA-pA |
| | Old World Monkeys | Oba | 2 | 951 | 1185 | | | | m29-m181-m29-m101-pA-pA |
| | | Gba | 2 | 949 | 1178 | | | | m29-m181-m29-m101-pA-pA |
| | | Rmc | 2 | 939 | 1171 | | | | m29-m181-m29-m101-pA-pA |
| | | Smc | 2 | 939 | 1170 | | | | m29-m181-m29-m101-pA-pA |
| | | Ver | 2 | 950 | 1180 | | | | m29-m181-m29-m101-pA-pA |
| | | Clb | 2 | 952 | 1188 | | | | m29-m181-m29-m101-pA-pA |
| | | Gsn | 2 | 952 | 1186 | | | | m29-m181-m29-m101-pA-pA |
| | | Bsn | 2 | 952 | 1182 | | | | m29-m181-m29-m101-pA-pA |
| | | Smn | 2 | 951 | 1182 | | | | m29-m181-m29-m29p-m101-pA-pA |
| | New World Monkeys | Cap | 2 | 930 | 1161 | | | | m29-m181-m29-m29p-m101-pA-pA |
| | | Sqm | 2 | 920 | 1157 | | | | m29-m181-m29-m29p-m101-pA-pA |
| | | Nmn | 2 | 944 | 1176 | | | | m29-m181-m29-m29p-m101-pA-pA |
| | Prosimians | Bby | 2 | 937 | 1157 | | | | m29-m181-m29-m101-pA-pA |
| | | Col | 2 | 965 | 1194 | | | | m29-m181-m29-m29p-m101-pA-pA |
| | | Lem | 2 | 928 | 1155 | | | | m29-m181-m29-m101-pA-pA |
| | Rodents and Lagamorphs | Mse | 2 | 915 | 1162 | | | | m29-m181-m29-m29p-m101-pA-pA |
| | | Krt | 5 | 715 | 964 | 1181 | 1232 | 1261 | m29-m181-m29-m29p-m101-pA-pA-pA-pA-pA |
| | | Bvr | 2 | 956 | 1175 | | | | m29-m181-m29-m29p-m101-pA-pA |
| | | Nmr | 4 | 865 | 1062 | 1405 | 1413 | | m29-m181-m29-m29p-m101-pA-pA-pA-pA |
| | | Rab | 2 | 934 | 1174 | | | | m29-m181-m29-m29p-m101-pA-pA |
| | Other Mammals | Swh | 2 | 956 | 1185 | | | | m29-m181-m29-m29p-m101-pA-pA |
| | | Bwh | 2 | 954 | 1192 | | | | m29-m181-m29-m101-pA-pA |
| | | Bov | 2 | 955 | 1191 | | | | m29-m181-m29-m29p-m101-pA-pA |
| | | Bat | 2 | 944 | 1199 | | | | m29-m181-m29-m29p-m101-pA-pA |
| | | Png | 5 | 924 | 1013 | 1148 | 1524 | 1704 | m29-m181-m29-m29p-m101-pA-pA-pA-pA-pA |
| | | Ele | 1 | 901 | | | | | m29-m181-m29-m29p-m101-pA |
| | Marsupials | Wal | 3 | 1246 | 1476 | 2604 | | | m29-m181-m29-m29p-m101-pA-pA |
| | | Kbr | 2 | 1281 | 1516 | | | | m29-m181-m29-m29p-m101-pA-pA |
| | | Wom | 2 | 1255 | 1490 | | | | m29-m181-m29-m29p-m101-pA-pA |
| | Monotremes | Plp | 2 | 1184 | 1406 | 3066 | | | m29-m181-m29-m29p-m29p-m101-pA-pA-pA |
| | | Ech | 3 | 1162 | 1364 | 2575 | | | m29-m181-m29-m29p-m101-pA-pA-pA |
| Sauropsids | Archosaurs | Can | 1 | 563 | | | | | m29-m181-m29-pA |
| | | Flc | 2 | 203 | 563 | | | | m29-m181-pA-m29-pA |
| | | Wts | 1 | 174 | | | | | m29-m181-pA-m29-pA |
| | | Zfn | 2 | 195 | 560 | | | | m29-m181-pA-m29-pA |
| | | Wtm | 3 | 210 | 527 | 1088 | | | m29-m181-pA-m29-pA-pA |
| | | Kap | 2 | 184 | 516 | | | | m29-m181-pA-m29-pA |
| | | Fal | 2 | 215 | 560 | | | | m29-m181-pA-m29-pA |
| | | Eag | 2 | 215 | 571 | | | | m29-m181-pA-m29-pA |
| | | Rpg | 2 | 194 | 537 | | | | m29-m181-pA-m29-pA |
| | | Ckn | 2 | 175 | 492 | | | | m29-m181-pA-m29-pA |
| | | Jql | 2 | 184 | 505 | | | | m29-m181-pA-m29-pA |
| | | Hgf | 2 | 175 | 505 | | | | m29-m181-PA-m29-PA |
| | | Dck | 2 | 191 | ? | | | | m29-m181-PA-m29-n-PA |
| | | Tin | ? | ? | ? | | | | m29-m181-n |
| | | Ctn | ? | ? | ? | | | | m29-m181-m29 |
| | | Cal | 2 | 271 | 1269 | | | | m29-m181-pA-m29-m29p-m101-pA |
| | | Aal | 2 | 271 | 1269 | | | | m29-m181-pA-m29-m29p-m101-pA |
| | Testudines | Str | 1 | 1024 | | | | | m29-m181-m29-m101-pA |
| | | Dtr | 1 | 1023 | | | | | m29-m181-m29-m101-pA |
| | | Ptr | 1 | 1024 | | | | | m29-m181-m29-m101-pA |
| | | Ttr | 1 | 1014 | | | | | m29-m181-m29-m101-pA |
| | | Gtr | 1 | 1013 | | | | | m29-m181-m29-m101-pA |
| | Squamates | Ano | 9 | 368 | 460 | 630 | 746 | 854 | m29-m181-m29-pA-pA-pA-pA-pA-pA-pA-pA-pA |
| | | Bdr | ? | ? | ? | ? | | | m29-m181-n-m29p-pA-pA-pA |
| | | Jgk | 2 | 764 | 777 | | | | m29-m181p-m29-m29p-pA-pA |
| | | Pyt | 4 | 740 | 753 | 763 | 1428 | | m29-m181-m29-m29p-pA-pA-pA-pA |
| | | Grs | 3 | 750 | 759 | 773 | | | m29-m181-m29-m29p-pA-pA-pA |
| | | Vip | 3 | 741 | 750 | 764 | | | m29-m181-m29-m29p-pA-pA-pA |
| | | Add | 3 | 740 | 749 | 763 | | | m29-m181-m29-m29p-pA-pA-pA |
| | | Rts | 3 | 739 | 748 | 762 | | | m29-m181-m29-m29p-pA-pA-pA |
| | | Jar | ? | ? | | | | | m29-m181-m29 |
| | | Blh | 2 | 718 | 727 | | | | m29-m181-m29-m29p-pA-pA |
| | | Cob | 3 | 772 | 782 | 796 | | | m29-m181-m29-m29p-pA-pA-pA |
| | | Mts | 2 | 784 | 802 | | | | m29-m181-m29-m29p-pA-pA |
| | | Ebs | 3 | 753 | 762 | 776 | | | m29-m181-m29-m29p-pA-pA-pA |
| | | Ylk | 3 | 767 | 776 | 790 | | | m29-m181-m29-m29p-pA-pA-pA |

m29: micRNA29 target site; m29p: micRNA29 target site incomplete by 1 flanking base
m181: micRNA181 target site; m101: micRNA101 target site
pA: polyadenylation site; n: gap in available sequence

#### Supplementary Table S6
##### Presence of Exons 34 and 35 in Genomic Sequences of Primate Tropoelastins

| Species Group | Species | Exon 34 Genomic Sequence | Exon 34 RefSeq Sequence | Exon 34 TSA/EST Sequence | Exon 34 Acceptor Site[1] | Exon 34 Donor Site[1] | Exon 35 Genomic Sequence | Exon 35 RefSeq Sequence | Exon 35 TSA/EST Sequence | Exon 35 Acceptor Site[1] | Exon 35 Donor Site[1] |
|---|---|---|---|---|---|---|---|---|---|---|---|
| Great Apes | Human | **absent** | NP | NP | - | - | **absent** | NP | NP | - | - |
| | Bonobo | **absent** | NP | - | - | - | **absent** | NP | - | - | - |
| | Chimpanzee | **absent** | NP | NP | - | - | **absent** | NP | NP | - | - |
| | Gorilla | **absent** | NP | - | - | - | **absent** | NP | - | - | - |
| | Orangutan | **absent** | NP | - | - | - | **absent** | NP | - | - | - |
| | Gibbon | gap[2] | - | - | - | - | gap[2] | - | - | - | - |
| Old World Monkeys | Olive Baboon | present | P | NP | 0.99 | 0.98 | present | P | NP | 0.88 | 1.00 |
| | Gelada Baboon | present | P | - | 0.99 | 0.98 | present | P | - | 0.88 | 1.00 |
| | Mandrill | gap[2] | - | - | - | - | present | NP | - | 0.88 | 1.00 |
| | Rhesus Monkey | present | P | - | 0.99 | 0.94 | present | P | NP | 0.87 | 1.00 |
| | Southern Pig-tailed Macaque | present | P | - | 0.99 | 0.98 | present | P | - | 0.88 | 1.00 |
| | Green Monkey | present | - | - | 0.90 | 0.98 | present | - | - | 0.88 | 1.00 |
| | Angolan Colobus Monkey | **absent** | NP | - | - | - | **absent** | NP | - | - | - |
| | Golden Snub-nosed Monkey | present | P | - | 0.99 | 0.98 | present | P | - | 0.88 | 1.00 |
| | Black Snub-nosed Monkey | present | - | - | 0.99 | 0.98 | present | - | - | 0.88 | 1.00 |
| | Sooty Mangabey | present | NP | - | 0.99 | 0.98 | present | P | - | 0.88 | 1.00 |
| New World Monkeys | Capuchin Monkey | present | P | - | 0.99 | 0.98 | present | P | - | 0.95 | 1.00 |
| | Bolivian Squirrel Monkey | present | - | - | 0.99 | 0.98 | present | - | - | 0.97 | 1.00 |
| | Nancy Ma Night Monkey | present | P | - | 0.99 | 0.98 | present | P | - | 0.92 | 1.00 |
| | Marmoset | present | P | - | 1.00 | 0.98 | present | P | P | 0.92 | 1.00 |
| Prosimians | Bushbaby | present | P | - | 0.99 | 0.98 | present | P | - | 0.94 | 1.00 |
| | Grey Mouse Lemur | present | P | - | 0.99 | 0.99 | present | P | - | 0.99 | 1.00 |
| | Tree Shrew | present | P | - | 0.99 | 0.98 | present | P | - | 0.70 | 1.00 |
| | Colugo | present | - | - | 0.99 | 0.99 | **absent** | - | - | - | - |

1. Splice site probabilities (0-1) predicted using NNSplice 0.9, http://www.fruitfly.org/seq_tools/splice.html

2. Sequence of exons 34 and 35 lost in sequencing gap

P predicted in supporting sequence

NP not predicted in supporting sequence

\- no data

### Supplementary Table S7
#### Expression of Exon 22 in Primate Tropoelastins

| Species Group | Species | Genomic Sequence | Acceptor Site[1] | Donor Site[1] | RefSeq Isoforms ( predicting exon 22/total) | TSA/EST Sequence |
|---|---|---|---|---|---|---|
| Great Apes | Human | GAAGAGVLGGLVPGAPGAVPGVPGTGGVP | <0.01 | 0.98 | 1/13 | - |
| | Bonobo | GAAGAGVLGGLVPGAPGAVPGVPGTGGVP | 0.92 | 0.98 | 1/13 | - |
| | Chimpanzee | GAAGAGVLGGLVPGAPGAVPGVPGTGGVP | 0.92 | 0.89 | 0/6 | NP |
| | Gorilla | GAAGAGVLGGLVPGAPGAIPGVPGTGGVP | 0.90 | 0.98 | 0/4 | - |
| | Orangutan | GAAGAGVLGGLVPGAPGAVPGVPGTGGVP | 0.76 | 0.98 | 0/2 | - |
| | Gibbon | GAAGAGVLGGLVPGAPGVVPGVPGIGGVP | 0.83 | 0.89 | 1/1 | - |
| | | | | | 3/39 | |
| Old World | Olive Baboon | GAAGAGVLGGLVPGAGGIVPGVPGVGGVP | 0.90 | 0.98 | 7/8 | NP |
| Monkeys | Gelada Baboon | GAAGAGVLGGLVPGAGGVVPGVPGVGGVP | 0.90 | 0.98 | 1/1 | - |
| | Mandrill | GAAGAGVLGGLVPGAGGVVPGVPGVGGVP | 0.90 | 0.98 | 0/1 | - |
| | Rhesus Monkey | GAAGAGVLGGLVPGAGGVVPGVPGVGGVP | - | - | 7/7 | - |
| | Southern Pig-tailed Macaque | GAAGAGVLGGLVPGAGGVVPGVPGVGGVP | 0.90 | 0.98 | 6/10 | - |
| | Green Monkey | GAAGAGVLGGLVPGAGGIVPGVPGVGGVP | 0.90 | 0.98 | 0/1 | - |
| | Angolan Colobus Monkey | GAAGAGVLGGLVPGAGGVVPGVPGVGGVP | 0.85 | 0.98 | 0/1 | - |
| | Golden Snub-nosed Monkey | GAAGAGVLGGLVPGAGGVVPGVPGVGGVP | 0.90 | 0.98 | 8/10 | - |
| | Black Snub-nosed Monkey | GAAGAGVLGGLVPGAGGVVPGVPGVGGVP | 0.90 | 0.98 | - | - |
| | Sooty Mangabey | GAAGAGVLGGLVPGAGGVVPGVPGVGGVP | 0.90 | 0.98 | 7/10 | - |
| | | | | | 36/49 | |
| New World | Capuchin Monkey | GAGGAGVLGGLVPGAGVAVPGVPGVGGVP | 0.71 | 0.99 | 13/13 | - |
| Monkeys | Bolivian Squirrel Monkey | gap[2] | - | - | 0/1 | - |
| | Nancy Ma's Night Monkey | GAGGAGVLGGLVPGAGVAVPGVPGTGGVP | 0.7 | 0.99 | 1/1 | - |
| | Marmoset | GAGGAGVLGGLVPGAGVAVPGVPGTGGVP | 0.74 | 0.99 | 0/1 | P |
| | | | | | 14/16 | |
| Prosimians | Bushbaby | GAAGAAALGGLVPGAVPGVPGAVPGVPGAVPGVPGTGGVP | 0.99 | 0.99 | 3/3 | - |
|  | Grey Mouse Lemur | GAAGAGALGGLVPG-----------VPGAVPGVPGAEGIP | 0.95 | 0.98 | 8/8 | - |
|  | Colugo | GPGGVGALGGLVPG-------------GAVPGVPGIGGVP | 0.98 | 0.99 | 1/1 | - |
| | Tree Shrew | GAGGAVPGVSGVPG-VPGVPG-VPGVPG-VPGVPGTGGVP | 0.95 | 0.98 | - | - |
| | | | | | 12/12 | |

1. Splice site probabilities (0-1) predicted using NNSplice 0.9, (http://www.fruitfly.org/seq_tools/splice.html)
2. Sequence of the exon region lost in sequencing gap

P predicted in supporting sequence

NP not predicted in supporting sequence

- no data

Supplementary Table S8

Exon 4a in Synapsid Tropoelastins

| Species Group | Species | Domain 4a Sequence | Acceptor Site[1] | Donor Site[1] | RefSeq Sequence | TSA/EST Sequence |
|---|---|---|---|---|---|---|
| Great Apes | Human | DARILGAFGA | 0.99 | 0.99 | NP | NP |
| | Bonobo | DAGILGAFGA | 0.99 | 0.99 | NP | - |
| | Chimpanzee | DAGILGAFGA | 0.99 | 0.99 | NP | NP |
| | Gorilla | DSGILGAFGA | 0.99 | 0.99 | NP | - |
| | Orangutan | DAGILGAFGA | 0.99 | <0.01 | NP | NP |
| | Gibbon | GAGIFGAFGA | 0.99 | 0.99 | NP | - |
| Old World Monkeys | Olive Baboon | GAGILGAFGA | <0.01 | 0.99 | NP | NP |
| | Gelada Baboon | GAGILGAFGA | <0.01 | 0.99 | NP | - |
| | Mandrill | GAGILGAFGA | <0.01 | 0.99 | NP | - |
| | Rhesus Monkey | GAGILGAFGA | <0.01 | 0.99 | NP | NP |
| | Southern Pig-tailed Macaque | GAGILGAFGA | <0.01 | 0.99 | NP | - |
| | Green Monkey | GAGILGAFGA | 1.00 | 0.99 | NP | - |
| | Angolan Colobus Monkey | GAGILGAFGA | 1.00 | 0.95 | NP | - |
| | Golden Snub-nosed Monkey | GAGILGAFGA | 1.00 | 0.99 | NP | - |
| | Black Snub-nosed Monkey | GAGILGAFGT | 1.00 | 0.98 | - | - |
| | Sooty Mangabey | GAGLLGAFGA | 0.68 | 1.00 | NP | - |
| New World Monkeys | Capuchin Monkey | GAGLLGAFGA | 1.00 | 0.99 | P | - |
| | Bolivian Squirrel Monkey | GAGILGAFGT | 0.99 | 0.99 | NP | - |
| | Nancy Ma's Night Monkey | GAGLLGAFGA | 1.00 | 0.99 | NP | - |
| | Marmoset | GAGLLGAFGA | 0.99 | 0.99 | P | NP |
| Prosiumians | Bushbaby | GARIFREFLA | <0.01 | 0.99 | NP | - |
| | Grey Mouse Lemur | **absent** | - | - | NP | - |
| | Tree Shrew | GAGLLGTFGA | 1.00 | 0.99 | P | NP |
| | Colugo | GAGALGAFGA | 0.99 | 0.96 | P | - |
| Rodents | Mouse | GAGLLGTFGA | 0.97 | 1.00 | P | P |
| | Kangaroo Rat | GAGV-GAFGA | 1.00 | 0.99 | P | - |
| | Beaver | GAGLLGAFGA | 0.99 | 1.00 | P | NP |
| | Naked Mole Rat | GAGLLGTFGI | 1.00 | 0.99 | P | NP |
| Lagomorphs | Rabbit | GAAIPGAFGA | 1.00 | 0.99 | P | P |
| Other Mammals | Sperm Whale | GAGLLGAFGP | 0.99 | 0.99 | P | - |
| | Beluga Whale | GAGLLGAFGP | 0.99 | 1.00 | P | P |
| | Narrow-ridged Porpoise | GAGLLGAFGP | 0.99 | 1.00 | - | P |
| | Bovine | GVGLLGAFGP | 1.00 | <0.01 | NP | NP |
| | Chinese Rufous Horseshoe Bat | GAGLLGAFGP | 0.99 | 1.00 | P | P |
| | Malayan Pangoln | GAGLLGAFGP | 0.99 | 1.00 | P | P |
| | Nine-Banded Armadillo | GAGLFGAFGA | 1.00 | 0.99 | P | - |
| | African Elephant | GVGLLGPFGT | 0.99 | <0.01 | NP | - |
| Marsupials | American Opossum | gap[2] | - | - | - | - |
| | Tasmanian Devil | **absent** | - | - | - | NP |
| | Fat-tailed Dunnart | gap[2] | - | - | - | - |
| | Tammar Wallaby | gap[2] | - | - | - | - |
| | Koala Bear | **absent** | - | - | NP | - |
| | Common Wombat | **absent** | - | - | NP | - |
| Monotremes | Platypus | **absent** | - | - | NP | - |
| | Echidna | **absent** | - | - | NP | - |

1. Splice site probabilities (0-1) predicted using NNSplice 0.9, http://www.fruitfly.org/seq_tools/splice.html

2. Sequence of the exon region lost in sequencing gap

P predicted in supporting sequences

NP not predicted in supporting sequences

\- no data

#### Supplementary Table S9
#### Exon 5a in Tropoelastins of Non-Placental Mammals

| Species Group | Species | Domain 5a Sequence | Acceptor Site[1] | Donor Site[1] | RefSeq Sequence | TSA/EST Sequence |
|---|---|---|---|---|---|---|
| Marsupials | American Opossum | **absent** | - | - | - | - |
| | Tasmanian Devil | GPGGLGA | 0.99 | 1.00 | - | P |
| | Fat-tailed Dunnart | GPGGLGG | [2] | [2] | - | P |
| | Tammar Wallaby | **gap**[3] | - | - | - | - |
| | Koala Bear | GPGGLGA | 0.99 | 0.99 | P | - |
| | Common Wombat | **absent** | - | - | NP | - |
| Monotremes | Platypus | **absent** | - | - | NP | - |
| | Echidna | **absent** | - | - | NP | - |

1. Splice site probabilities (0-1) predicted using NNSplice 0.9, http://www.fruitfly.org/seq_tools/splice.html
2. TSA sequence only. No splice site informaton
3. Sequence of the exon region lost in sequencing gap

P predicted in supporting sequence
NP not predicted in supporting sequence
\- no data

#### Supplementary Table S10
#### Truncation of Domain 13 in Avian Tropoelastins

| | Species Group | Species | Domain 13 Sequence |
|---|---|---|---|
| SYNAPSIDS | Great Apes | Human | GGY-GLPYTTGKLPY |
| | | Chimpanzee | GGY-GLPYTTGKLPY |
| | | Orangutan | GGY-GLPYTTGKLPY |
| | Old World Monkeys | Olive Baboon | GGY-GLPYSTGKLPY |
| | | Rhesus Macaque | GGY-GLPYSTGKLPY |
| | | Black Snub-nosed Monkey | GGY-GLPYSTGKLPF |
| | New World Monkeys | Capuchin Monkey | GGY-GLPYSTGKLPF |
| | | Marmoset | GGY-GLPYSTGKLPF |
| | Prosimians | Bushbaby | GGY-GLPYTTGKLPY |
| | | Tree Shrew | GGY-GLPYSTGKLPY |
| | Rodents | Mouse | GGY-GLPYTNGKLPY |
| | Lagomorphs | Rabbit | GGY-GLPYSTGKLPY |
| | Other Mammals | Sperm Whale | GGY-GLPYSTGKLPY |
| | | Chinese Rufous Horseshoe Bat | GGY-GLPYSTGKLPY |
| | Marsupials | American Opossum | GGY-GLPYSTGKLPY |
| | | Common Wombat | GGY-GLPYSTGKLPY |
| | Monotremes | Platypus | GGY-GLPYSTGKLPY |
| SAUROPSIDS | Birds | Collared Flycatcher | GGY-GLPYTN[1] |
| | | Zebra Finch | GGY-GLPYTN |
| | | Kakopo | GGY-GLPYST |
| | | Peregrine Falcon | GGY-GLPYST |
| | | Crested Ibis | GGY-GLPYTT |
| | | Chicken | GGY-RLPFVN |
| | | Australian Ostrich | GGY-GLPYST |
| | | Chilean Tinamou | GGY-GLPYST |
| | Alligators/ Crocodiles | Chinese Alligator | GGY-GLPYSTGKLPY |
| | | American Alligator | GGY-GLPYSTGKLPY |
| | Turtles | Red-eared Slider Turtle | GGY-GLPYSTGKLPY |
| | | Green Sea Turtle | GGY-GLPYSTGKLPY |
| | Lizards | Asian Grass Lizard | GSYYGLPYGAGKGPY |
| | Snakes | Burmese Python | GSYYGLPYGAGKLAF |
| | | King Cobra | GSYYGNSYGAGKLTF |

1. Truncation was present in all avian species, but absent from all other Amniote species

#### Supplementary Table S11
#### Truncation of Domain 36 in Squamate Tropoelastins

| | Species Group | Species | Domain 36 Sequence |
|---|---|---|---|
| Synapsids | Great Apes | Human | GGA-CL-GKACGRKRK |
| | | Chimpanzee | GGA-CL-GKACGRKRK |
| | | Orangutan | GGA-CL-GKACGRKRK |
| | Old World Monkeys | Olive Baboon | GAA-CL-GKSCGRKRK |
| | | Rhesus Macaque | GAA-CL-GKSCGRKRK |
| | | Black Snub-nosed Monkey | GGA-CL-GKSCGRKRK |
| | New World Monkeys | Capuchin Monkey | AGA-CL-GKACGRKRK |
| | | Marmoset | AGA-CL-GKACGRKRK |
| | Prosimians | Bushbaby | GGA-CL-GKSCGRKRK |
| | | Tree Shrew | GGA-CV-GKACGRKRK |
| | Rodents | Mouse | GGG-CF-GKSCGRKRK |
| | Lagomorphs | Rabbit | GGA-CL-GKSCGRKRK |
| | Other Mammals | Sperm Whale | GGA-CL-GKPCGRKRK |
| | | Chinese Rufous Horseshoe Bat | GGA-CL-GKSCGRKRK |
| | Marsupials | American Opossum | GGA-CL-GKMCGRKRK |
| | | Common Wombat | GGA-CL-GKLCGRKRK |
| | Monotremes | Platypus | GGA-CS-GKYCGRKRK |
| Sauropsids | Birds | Collared Flycatcher | GGAGCAAGKYCGRKRK |
| | | Kakopo | GGVGCAQGKYCGRKRK |
| | | Peregrine Falcon | GGVGCAQGKYCGRKRK |
| | | Chicken | GGVGCAQGKYCGRKRK |
| | | Chilean Tinamou | GGVGCAQGKYCGRKRK |
| | Alligators/Crocodiles | Chinese Alligator | GGAGC-QGKYCGRKRK |
| | Turtles | Red-eared Slider Turtle | GGVGCAQGKYCGRKRK |
| | | Goode's Thornscrub Tortise | GGVGCAQGKYCGRKRK |
| | | Green Sea Turtle | GGVGCAQGKYCGRKRK |
|  | Lizards | Anolis LIzard | GAL--------GRRRK[1] |
|  |  | Asian Grass Lizard | GGI--------GRKRK |
|  | Snakes | Burmese Python | RGV--------GRKRK |
|  |  | Garter Snake | RGV--------GRKRK |
|  |  | Pit Viper | RGV--------GRKRK |
|  |  | Timber Rattle Snke | RGV--------GRKRK |
|  |  | King Cobra | RGV--------GRKRK |
|  |  | Eastern Brown Snake | RGV--------GRKRK |
|  |  | Yellow-lipped Sea Krait | RGV--------GRKRK |

1. Truncation was present in all squamate species (lizards and snakes), but absent from other Amniote species

Supplementary Table S12
Examples of Recent Replication of Exon/ Domain Sequences of Some Amniotes
A. Avian (Passeriformes birds)

| Species | RefSeq | Domain # | Cross-linking Domain | Domain # | Hydrophobic Domain |
|---|---|---|---|---|---|
| Common Canary | XM_050981640.[illegible] | 15 | GVGAQAAAAKAAAKL | 16 | GAGVLPGVGGIPGVAPGVGIGGVPGVG |
| *(Serinus canaria)* | (variant X1) | 17 | VGGPAAAAAKAAAKAGAF | 18r1 | GPGA-----------VPGVGGVPGLVPGVGGVPGAVPGVGGVPGVA |
| | | 19r1 | GVPSAAAAAKAAKY | 18r2 | GAGVPGIGVGGVPGLVPGVGGVPGLVPGVGGVPGAVPGVGGVPGVA |
| | | 19r2 | GVPSAAAAAKAAKY | 18r3 | GAGVPGVGVGGVPGLVPGVGGVPGLVPGVGGVPGAVPGVGGVPGVA |
| | | 19r3 | GVPSAAAAAKAAKY | 18r4 | GAGVPGVGVGGVPGLVPGVGGVPGLVPGVGGVPGAVPGVGGVPGVA |
|  |  | 19r4 | GVPSAAAAAKAAKY | 18r5 | GAGVPGIGVGGVPGL--------------------VPGVGGVPGVA |
| | | 19r5 | GVPSAAAAAKAAKY | 18r6 | GAGVPGVGVGGVPGLVPGVGGVPGLVPGVGGVPGAVPGVGGVPGVA |
| | | 19r6 | GVPSAAAAAKAAKY | 18r7 | GAGVPGVGVGGVPGLVPGVGGVPGLVPGVGGVPGAVPGVGGVPGVA |
| | | 19r7 | GVPSAAAAAKAAKY | 18r8 | GAGVPGVGVGGVPGLVPGVGGVPGLVPGVAGVPGAVPGVGGVPGVA |
| | | 19r8 | GVPSAAAAAKAAKY | 20 | GAGVPGIAGVPGVPGVPGGPGVPGVPGVPGAVPGVG |
| | | 23 | VGGLAAAAAAKAAAKAAAI | 24 | GAGRVPGVGVPGVGVPAVGVPGLVPGVGVP |
| | | 25 | GGPAAAAKAAAKAAKY | 26 | GAGGLAPGVGGLAPGVGGLAPGVGGLAPGVGGLVPGVGGVP |
| | | 27 | GVGGPAAAAKAAAKAAKF | 28 | GAGVGGVPGVVPGVGGVPGVTPGVGGVPGLVPGVGVPGTGILPGA |
| | | 29a | GIPQVGVQPGAKPPKF | 29b | GVPGVGVPGVGGLP |
| Collared Flycatcher *(Ficedula albicollis)* | XM_016303078. (variant X1) | 15 | GVGAQAAAAKAAAKL | 16 | GAGVLPGVGGIPGVAPGVGVGGVPGVG |
|  |  | 17 | VGGPAAAAAAAAAKAAAKAGAF | 18r1 | GPGA-----------VPGVGGVPGLVPGVGGVPGAVPGVGGVPGVA |
| | | 19 r1 | GVPSAAAAAKAAKY | 18r2 | GAGVPGVGVGGVPGLVPGVGGVPGLVPGVGGVPGAVPGVGGVPGVA |
| | | 19r2 | GVPSAAAAAKAAKY | 18r3 | GAGVPGVGVGGVPGLVPGVGGVPGLVPGVGGVPGAVPGVGGVPGVA |
| | | 19r3 | GVPSAAAAAKAAKY | 18r4 | GAGVPGVGVGGVPGLVPGVGGVPGLVPGVGGVPGAVPGVGGVPGVA |
| | | 19r4 | GVPSAAAAAKAAKY | 18r5 | GAGVPGVGVGGVPGLVPGVGGVPGLVPGVGGVPGAVPGVGGVPGVA |
| | | 19r5 | GVPSAAAAAKAAKY | 18r6 | GAGVPGVGVGGVPGLVPGVGGVPGLVPGVGGVPGAVPGVGGVPGVA |
| | | 19r6 | GVPSAAAAAKAAKY | 20 | GAGVPGIAGVPGVPGVPGVPGVPGVPGVPGVPGVPGAVPGVG |
| | | 23 | VGGPAAAAAAKAAAKAAAI | 24 | GAGRVPGVGVPGVGVPGVGVPGVGVPGVGVPGLVPGVGVP |
| | | 25 | GGPAAAAKAAAKAAKY | 26 | GAGGLAPGVGGLVPGVGGLVPGVGGLVPGVGGVP |
| | | 27 | GVGGPAAAAKAAAKAAKF | 28 | GAGVGGVPGVVPGVGGVPGVTPGVGGVPGLVPGVGVPGTGILPGA |
| | | 29a | GIPQVGVQPGAKPPKF | 29b | GVPGVGVPGVGGLP |

Only sequences from the Replicated Regions of tropoelastins (domains 15-29b, Figure 2) are shown. Apparent domain replications are shaded.

Supplementary Table S12
Examples of Recent Replication of Exon/ Domain Sequences of Some Amniotes
B. Testudines (Turtles)

| Species | RefSeq | Domain # | Cross-linking Domain | Domain # | Hydrophobic Domain |
|---|---|---|---|---|---|
| Painted Turtle | XM_042845711.1 | 15 | GVGAQVAAAKAAAKY | 16 | GAGVVPGAAGIPGVGGLGGLVPGVGGLPGVAGVPGVA |
| (*Chrysemys* | (variant 1) | 17 | absent from genomic sequence | 18 | absent from genomic sequence |
| *picta bellii*) | | 19 r1 | GAGSPAAAAAAAAAKAAKY | 20r1 | GAGGVGGLVPGVGGVPGVVPGVGGVPGLVPGVGGVPGVG |
| | | 19r2 | AGLPAGAAAAAAAKAAKY | 20r2 | GAGGVGGLVPGVGGVPGVVPGVGGVPGVVPGVGGFPGVG |
| | | 23 | AISPAAAAAAKAAAKAAAY | 24 | GAGRVPGVAPGVGGLVPGIGGGLVPGVGGGLVPGVGGALVPGVG |
| | | 25 | GVVSPAAAAKAAAKAAKY | 26 | GAGGVGGLVPGGVGGLVPGGVGGLVPGGVGGLVPGGVGGIP |
| | | 27 | GLLSPAAAAKAAAKAAKY | 28 | GAGVAPGVGGVPGLVPGAGRVPVATPGAGGVPVVTPGTGGFVPGAGRVPGTGIVPGV |
| | | 29a | GIPQLGVQPGAKPPKY | 29b | absent from genomic sequence |
| Goode's | XM_030536861.1 | 15 | GVGAQVAAAKAAAKY | 16 | GAGVVPGAAGIPGVGGLGGLVPGVGGLPGVAGVPGVA |
| Thornscrub Turtle | (variant 1) | 17 | absent from genomic sequence | 18 | absent from genomic sequence |
| *(Gopherus* |  | 19r1 | GAESPAAAAAAAAAAKAAKY | 20r1 | GAAGVGGLVPGVAGVPGVVPGVPGVVPGVP-------GVPGVVPGVGGVPGVG |
| *evgoodei)* |  | 19r2 | AGIPAGAAAAAAAAKAAKY | 20r2 | GAAGVGGLVPGVAGVPGVVPGVPGVVPGVP-------GVPGVVPGVGVVPGVG |
|  |  | 19r3 | AGIPAG-AAAAAAAKAAKY | 20r3 | GAAGVGGLVPGVAGVPGVVPGVPGVVPGVP-------GVPGVVPGVGGVPGVG |
|  |  | 19r4 | AGIPAG-AAAAAAAKAAKY | 20r4 | GAAGVGGLVPGVAGVPGVVPGVPGVVPGVP-------GVPGVVPGVGGVPGVG |
| | | 19r5 | AGIPAG-AAAAAAAKAAKY | 20r5 | AAAGVGGLVPGVAGVPGVVPGVPGVVPGVPGVVPGVGGIPGVVPGVGGVPGVG |
| | | 23 | AVSPAAAAAAKAAAY | 24 | GAGRIPGVAPGVGGLVPGVGGVLVPGVG |
| | | 25 | GVVSPAAAAKASAKAAKY | 26 | GAGGVGGLVPGGVRGLVPGGVGGLVPGGVGGIP |
| | | 27 | GLSPAAAKAAAKAAKY | 28 | GAGLAPGVGGVPGLVPGVGRLPVATPGAGRLPVVTPGTGGVPVFVPGAGRVPGTGIIPGA |
| | | 29a | GIPQLGVQPGAKPPKY | 29b | absent from genomic sequence |

Only sequences from the Replicated Regions of tropoelastins (domains 15-29b, Figure 2) are shown. Apparent domain replications are shaded.

Figure S1

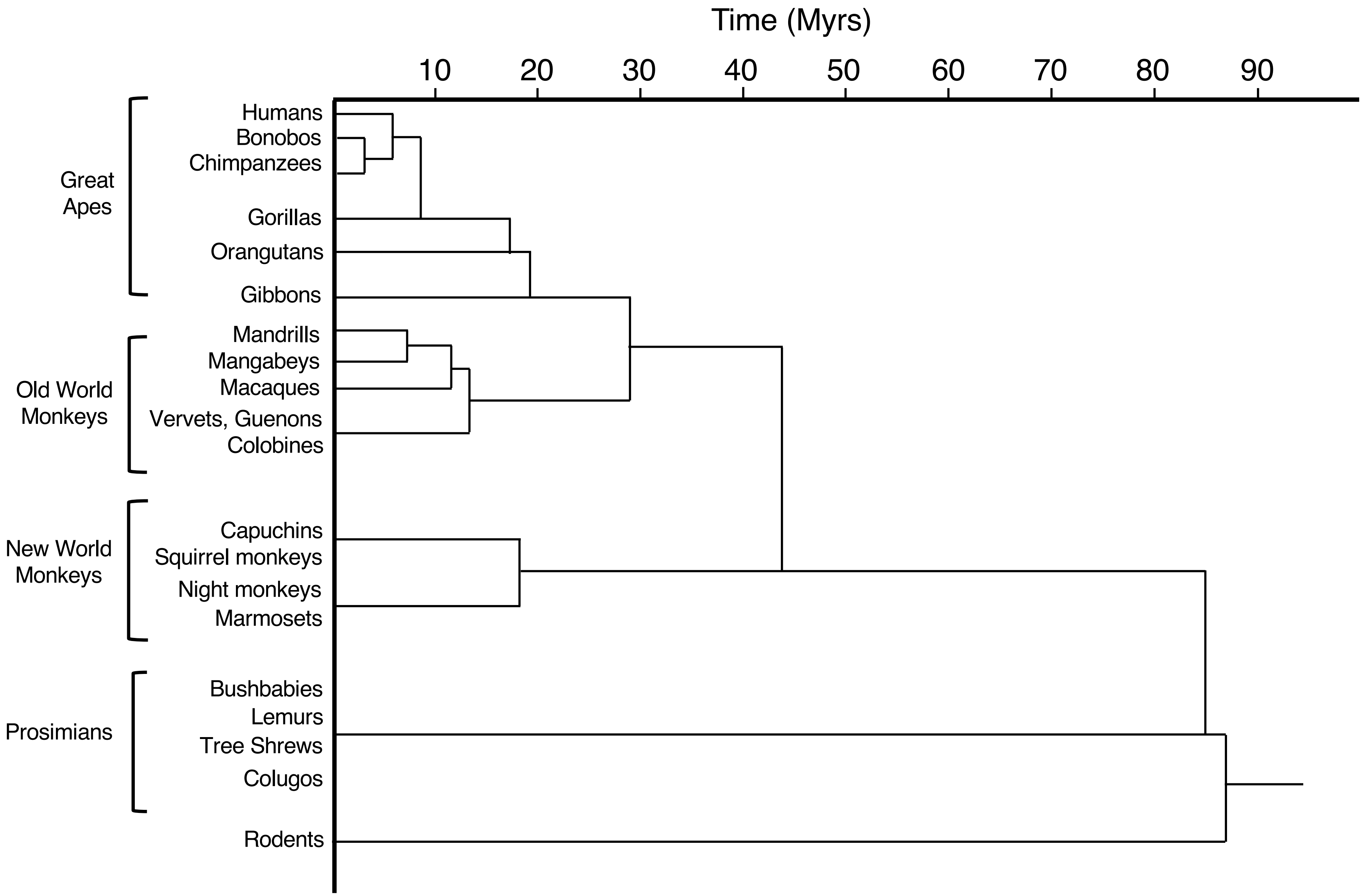
Time (Myrs)
10
20
30
40
50
60
70
80
90
Great Apes
Humans
Bonobos
Chimpanzees
Gorillas
Orangutans
Gibbons
Old World Monkeys
Mandrills
Mangabeys
Macaques
Vervets, Guenons
Colobines
New World Monkeys
Capuchins
Squirrel monkeys
Night monkeys
Marmosets
Prosimians
Bushbabies
Lemurs
Tree Shrews
Colugos
Rodents

Figure S2

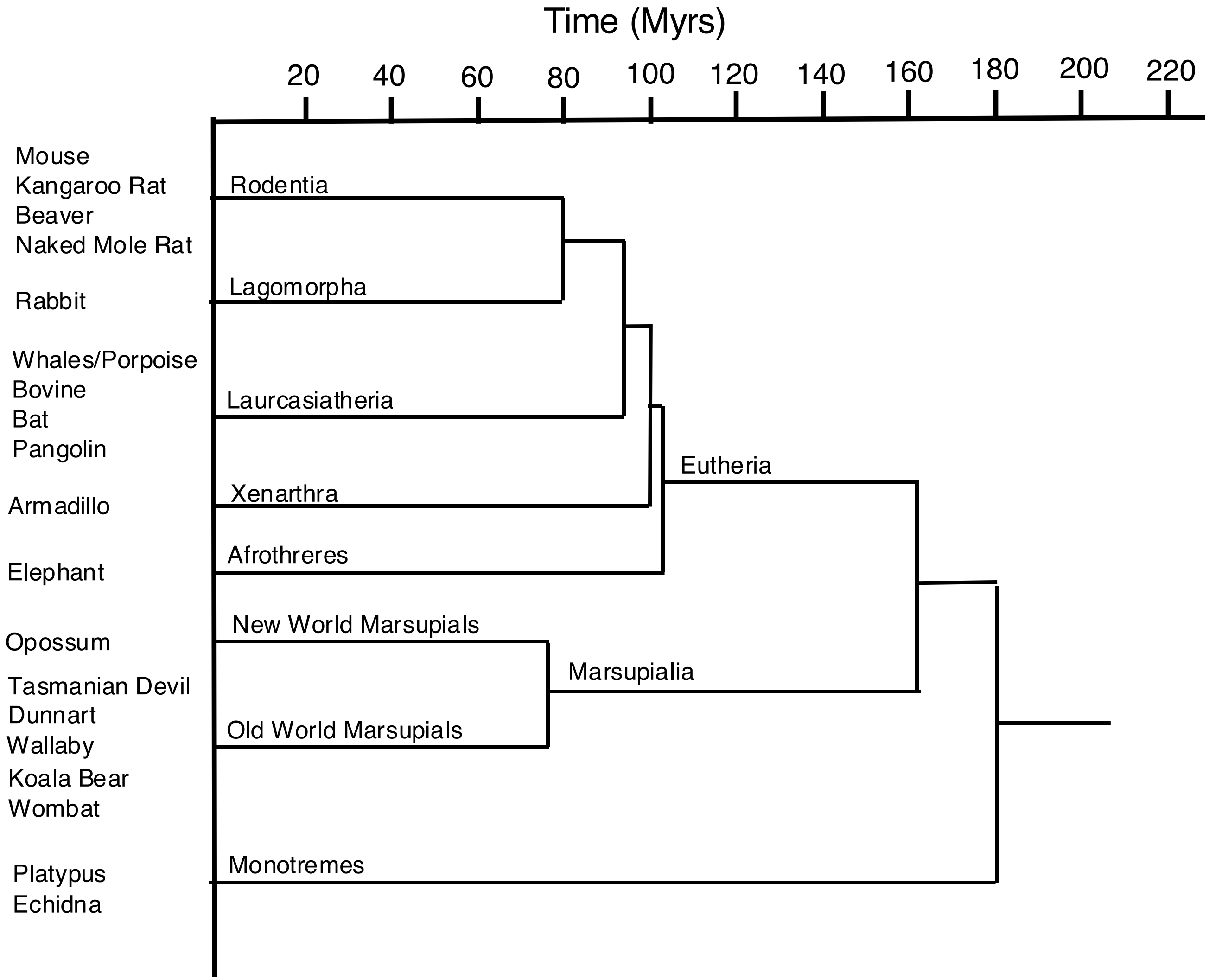
Time (Myrs)
20
40
60
80
100
120
140
160
180
200
220
Mouse
Kangaroo Rat
Beaver
Naked Mole Rat
Rabbit
Whales/Porpoise
Bovine
Bat
Pangolin
Armadillo
Elephant
Opossum
Tasmanian Devil
Dunnart
Wallaby
Koala Bear
Wombat
Platypus
Echidna
Rodentia
Lagomorpha
Laurcasiatheria
Xenarthra
Afrothreres
Eutheria
New World Marsupials
Marsupialia
Old World Marsupials
Monotremes

Figure S3

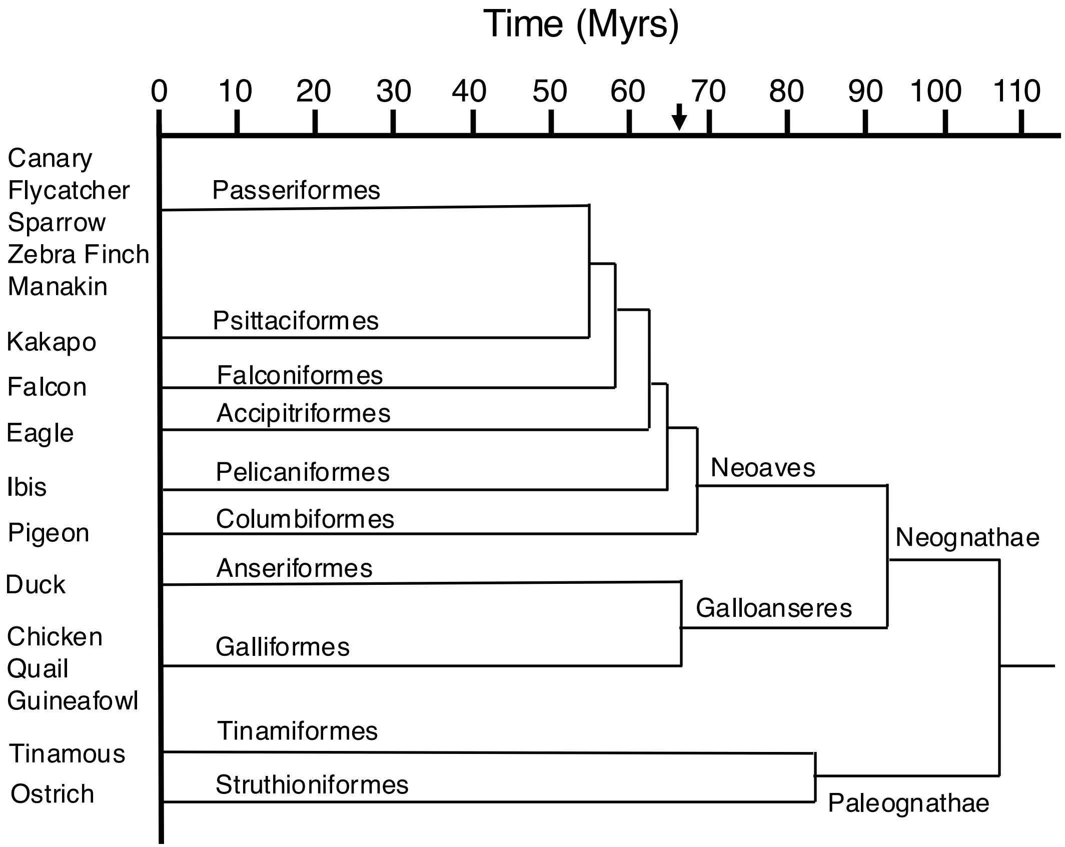
Time (Myrs)
0
10
20
30
40
50
60
70
80
90
100
110
Canary
Flycatcher
Sparrow
Zebra Finch
Manakin
Kakapo
Falcon
Eagle
Ibis
Pigeon
Duck
Chicken
Quail
Guineafowl
Tinamous
Ostrich
Passeriformes
Psittaciformes
Falconiformes
Accipitriformes
Pelicaniformes
Columbiformes
Anseriformes
Galliformes
Tinamiformes
Struthioniformes
Neoaves
Neognathae
Galloanseres
Paleognathae

Figure S4

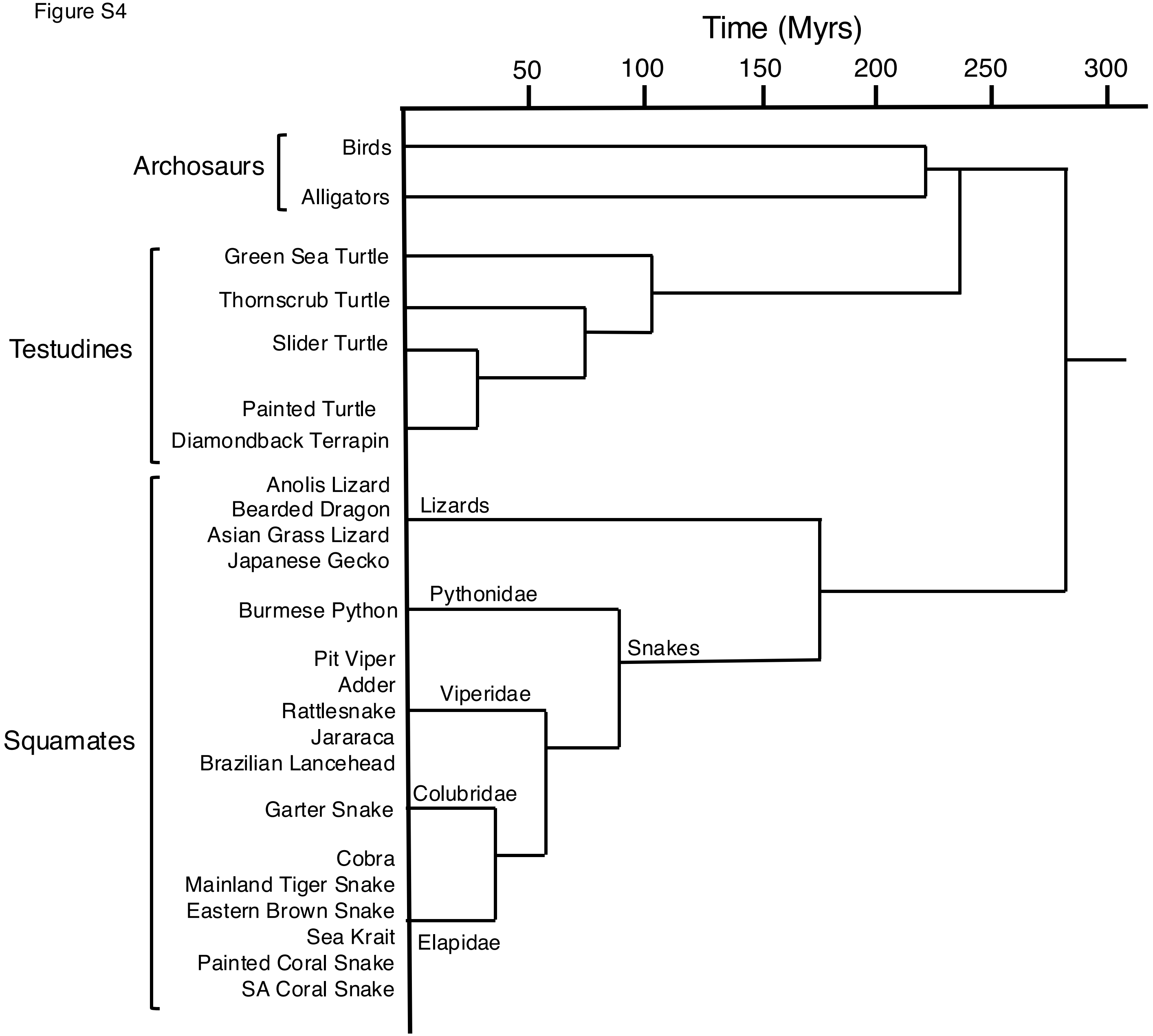

Time (Myrs)
50
100
150
200
250
300
Archosaurs
Birds
Alligators
Testudines
Green Sea Turtle
Thornscrub Turtle
Slider Turtle
Painted Turtle
Diamondback Terrapin
Squamates
Anolis Lizard
Bearded Dragon
Asian Grass Lizard
Japanese Gecko
Lizards
Burmese Python
Pythonidae
Snakes
Pit Viper
Adder
Rattlesnake
Jararaca
Brazilian Lancehead
Viperidae
Garter Snake
Colubridae
Cobra
Mainland Tiger Snake
Eastern Brown Snake
Sea Krait
Painted Coral Snake
SA Coral Snake
Elapidae

Figure S5

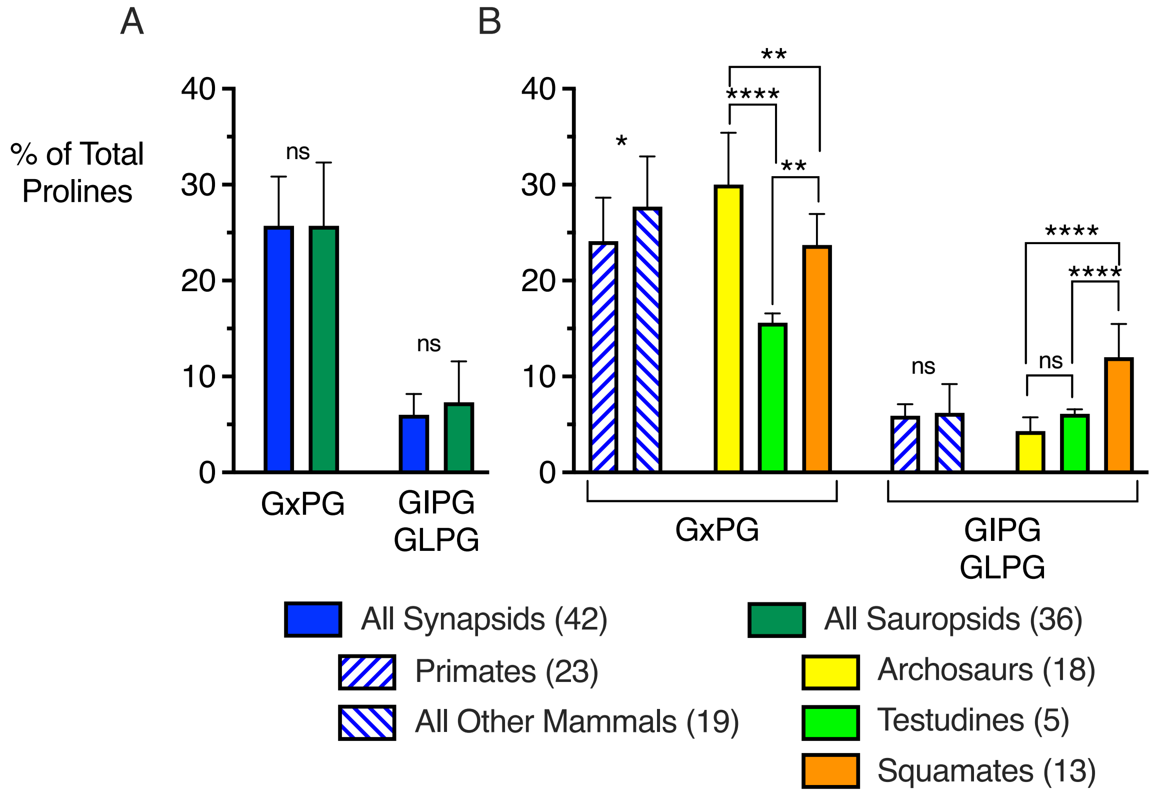
A
B
% of Total Prolines
40
30
20
10
0
ns
ns
GxPG
GIPG
GLPG
*
**
****
**
****
****
ns
ns
GxPG
GIPG
GLPG
All Synapsids (42)
All Sauropsids (36)
Primates (23)
All Other Mammals (19)
Archosaurs (18)
Testudines (5)
Squamates (13)

Figure S6

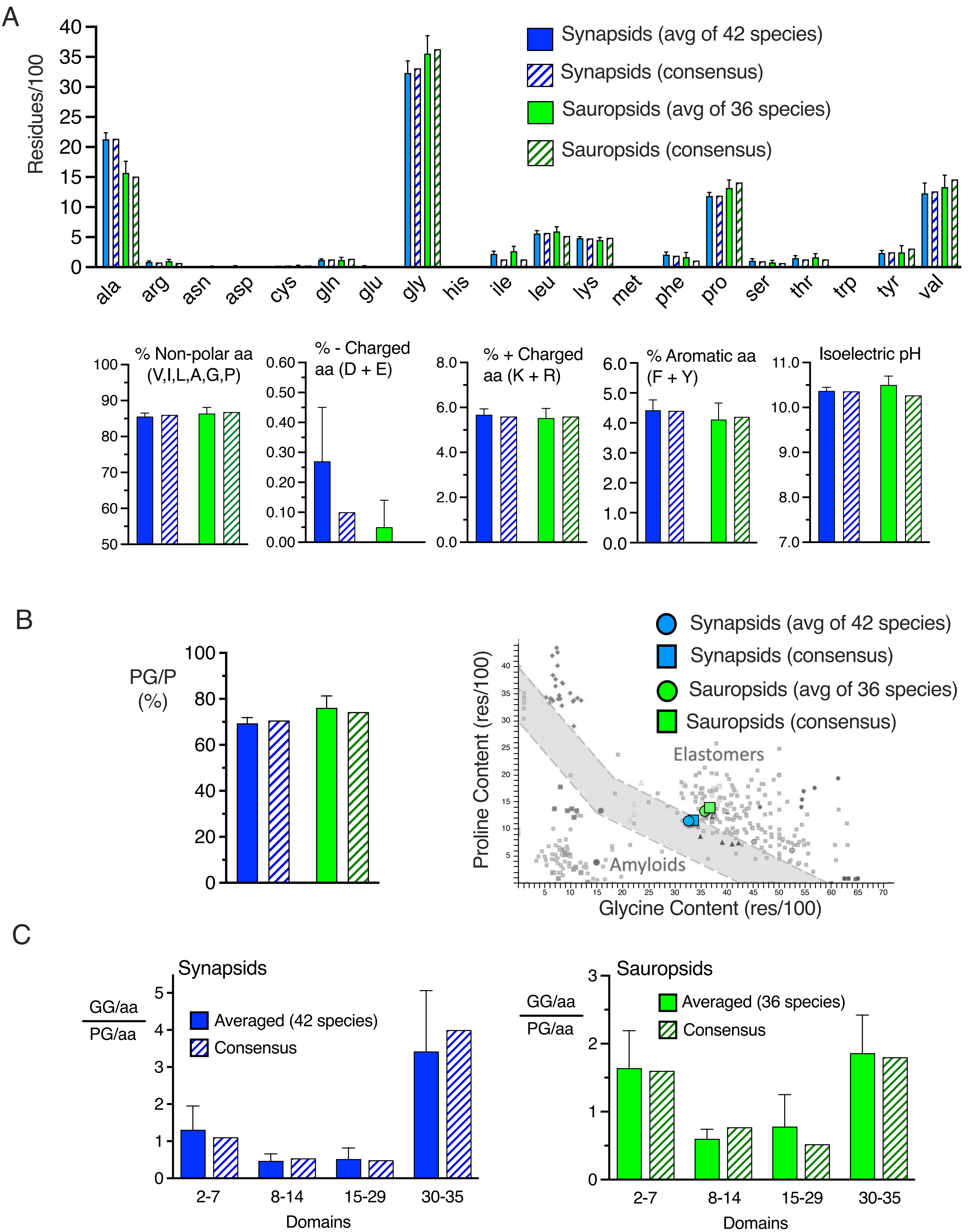
A
Residues/100
Synapsids (avg of 42 species)
Synapsids (consensus)
Sauropsids (avg of 36 species)
Sauropsids (consensus)
ala arg asn asp cys gln glu gly his ile leu lys met phe pro ser thr trp tyr val
% Non-polar aa (V,I,L,A,G,P)
% - Charged aa (D + E)
% + Charged aa (K + R)
% Aromatic aa (F + Y)
Isoelectric pH
B
PG/P (%)
Synapsids (avg of 42 species)
Synapsids (consensus)
Sauropsids (avg of 36 species)
Sauropsids (consensus)
Elastomers
Amyloids
Proline Content (res/100)
Glycine Content (res/100)
C
Synapsids
GG/aa / PG/aa
Averaged (42 species)
Consensus
Domains
2-7
8-14
15-29
30-35
Sauropsids
GG/aa / PG/aa
Averaged (36 species)
Consensus
Domains

Figure S7

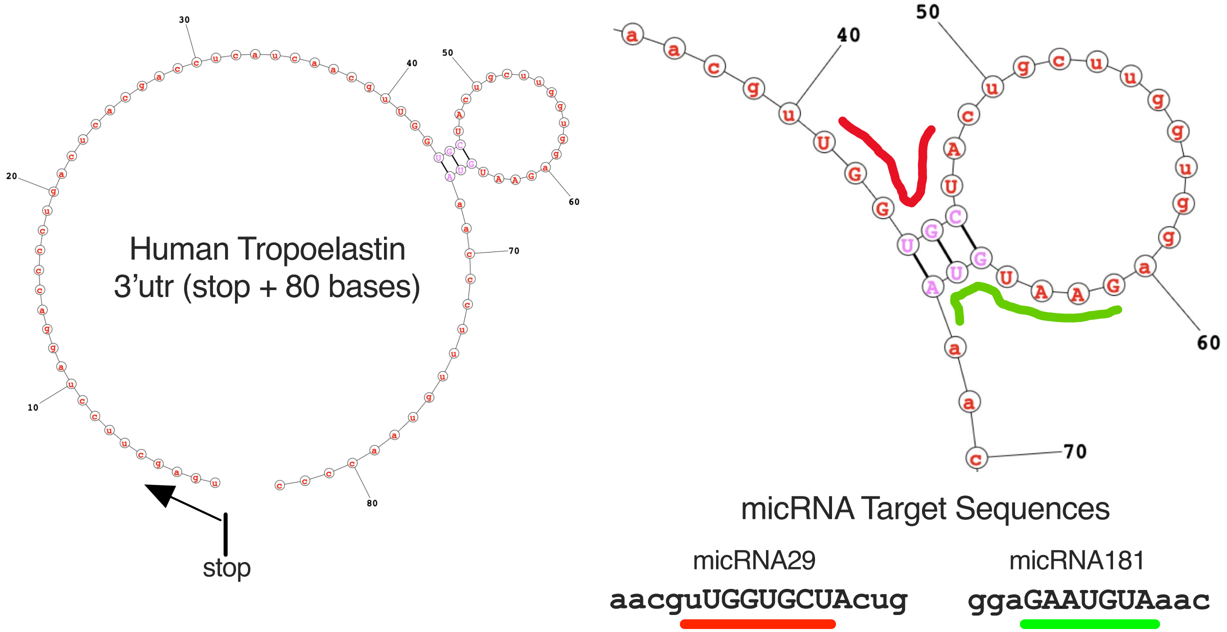
Human Tropoelastin
3'utr (stop + 80 bases)
stop
micRNA Target Sequences
micRNA29
aacguUGGUGCUAcug
micRNA181
ggaGAAUGUAaac
